# A *Drosophila* glial cell atlas reveals a mismatch between detectable transcriptional diversity and morphological diversity

**DOI:** 10.1101/2022.08.01.502305

**Authors:** Inês Lago-Baldaia, Maia Cooper, Austin Seroka, Chintan Trivedi, Gareth T. Powell, Stephen Wilson, Sarah D. Ackerman, Vilaiwan M. Fernandes

## Abstract

Morphology is a defining feature of neuronal identity. Like neurons, glia display diverse morphologies, both across and within glial classes, but are also known to be morphologically plastic. Here, we explored the relationship between glial morphology and transcriptional signature using the *Drosophila* central nervous system, where glia are categorized into five main classes (outer and inner surface glia, cortex glia, ensheathing glia, and astrocytes), which show within-class morphological diversity. We analysed and validated single cell RNA sequencing data of *Drosophila* glia in two well-characterized tissues from distinct developmental stages, containing distinct circuit types: the embryonic ventral nerve cord (motor) and the adult optic lobes (sensory). Our analysis identified a new morphologically and transcriptionally distinct surface glial population in the ventral nerve cord. However, many glial morphological categories could not be distinguished transcriptionally, and indeed, embryonic and adult astrocytes were transcriptionally analogous despite differences in developmental stage and circuit type. While we did detect extensive within-class transcriptomic diversity for optic lobe glia, this could be explained entirely by glial residence in the most superficial neuropil (lamina) and an associated enrichment for immune-related gene expression. In summary, we generated a single-cell transcriptomic atlas of glia in *Drosophila*, and our extensive *in vivo* validation revealed that glia exhibit more diversity at the morphological level than was detectable at the transcriptional level. This atlas will serve as a resource for the community to probe glial diversity and function.

## Introduction

Nervous systems contain more distinct cell types than any other organ. This cellular diversity underlies the complexity and multifunctionality of circuits and processing networks in the brain and, thus, defines the breadth of an animal’s behavioural repertoire. Not surprisingly, categorising neural cell types has long been, and continues to be, a major endeavour in the field. Although much emphasis has been placed on categorising neuronal diversity, we know much less about the extent of glial diversity. Given their pivotal roles in every aspect of nervous system development and function (Allen and Lyons, 2018; Lago-Baldaia et al., 2020), understanding glial diversity is also imperative.

Morphological diversity among glia has been documented alongside that of neurons for over a century (Ramón y Cajal, 1899). This morphological heterogeneity exists not only between broad glial classes (*i.e.* astrocytes, oligodendrocytes, Schwann cells and microglia) but also within classes (Foerster et al., 2019; Harty and Monk, 2017; Kamen et al., 2022; Khakh and Sofroniew, 2015). It has long been appreciated that mammalian astrocytes from different brain regions vary in morphology (Lanjakornsiripan et al., 2018; Ramón y Cajal, 1899; reviewed in detail in Zhou et al., 2019), and recent advances in RNA-sequencing (RNA-seq) technologies have revealed regionalized molecular diversity in astrocytes and other central nervous system (CNS) glial cell classes (*e.g.* oligodendrocyte progenitor cells and microglia) (Batiuk et al., 2020; Bayraktar et al., 2020; Chai et al., 2017; Dimou and Gallo, 2015; Grabert et al., 2016; Marques et al., 2018; Spitzer et al., 2019). Confoundingly, astrocytes are known to be highly plastic cells. Most notably, in response to injury, astrocytes undergo a process called astrogliosis, become “reactive,” and alter their morphology dramatically (Escartin et al., 2021). It is clear that astrocyte reactivity represents a change in cell state due to underlying differences in environment. Thus, in healthy conditions, it is difficult to distinguish whether the morphological diversity of astrocytes is a consequence of cell-fate diversity and/or cell-state. In other words, what is the relationship between glial morphology and transcriptional profile?

*Drosophila* glia share several key morphological and functional attributes with their vertebrate counterparts, including maintaining neurotransmitter and ionic homeostasis, providing trophic support for neurons, acting as immune cells, and modifying neural circuit function (Bittern et al., 2021; Freeman, 2015; Freeman and Doherty, 2006; Lago-Baldaia et al., 2020). In the *Drosophila* CNS, neuropils contain synaptic connections, while neuronal cell bodies are located at the cortex, around the periphery of neuropils; axon tracts connect different neuropils to each other. *Drosophila* glia can be categorised based on morphology and by their association with these anatomical structures as either outer or inner surface glia, cortex glia, ensheathing glia, or astrocytes (Figures 1 and 2). Surface glia comprise two sheet-like glia called the perineurial and subperineurial glia (Freeman, 2015; Pogodalla et al., 2022). Together these form a double-layered surface that spans the nervous system, which acts as a blood (or hemolymph)-brain barrier (BBB) (Freeman, 2015; Pogodalla et al., 2022). Cortex glia envelop neuronal cell bodies in cortical regions of the CNS, whereas ensheathing glia can wrap axonal tracts between neuropils (*aka* tract ensheathing glia) or wrap neuropil borders (Edwards and Meinertzhagen, 2010; Freeman, 2015). Astrocytes also inhabit neuropil regions with ensheathing glia, but extend many fine projections into the neuropil to associate with neuronal synapses, akin to vertebrate astrocytes (Freeman, 2015; Stork et al., 2014). Although *Drosophila* has a simplified nervous system with reduced numbers of glia relative to mammals, striking morphological diversity exists between and within glial cell classes during development and in the adult (Edwards et al., 2012; Edwards and Meinertzhagen, 2010; Kremer et al., 2017; Peco et al., 2016). For example, in the highly ordered visual system, astrocytes of distinct morphologies can be found across neuropils and within the same neuropil (Edwards et al., 2012; Richier et al., 2017). Whether these morphological categories correspond to distinct subclasses with unique transcriptional profiles and functions is not known.

**Figure 1.**
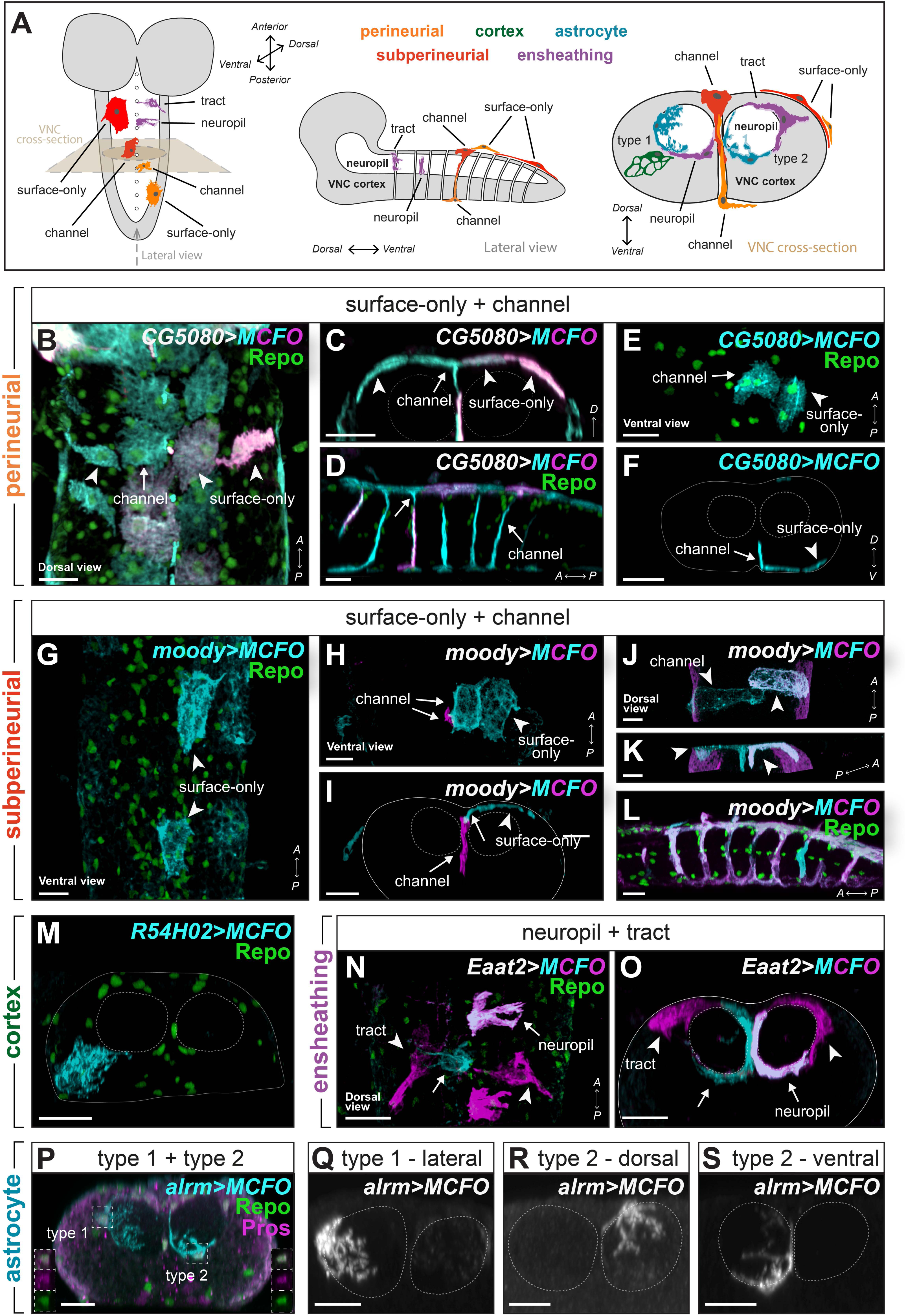
Morphologies of newly hatched VNC glia. **(A)** Schematics of the embryonic CNS along different axes with the five glial classes indicated. Cortex (neuronal cell bodies) indicated in grey. **(B)** Dorsal surface view of the VNC showing surface-only and channel perineurial and subperineurial glial cells on the surface with characteristic fibrous morphology and loose tiling. **(C)** Cross-sectional view of the VNC showing the same cells as in (B) now showing the tapered projection from a (dorsal) channel perineurial glial cell that tiles with neighbouring surface-only perineurial glial cells. **(D)** Lateral view of the VNC showing channel perineurial glia on the ventral and dorsal surfaces, each sending a single projection with ventral channel perineurial glia sending longer processes than their dorsal counterparts. **(E)** Ventral surface view of the VNC showing surface-only and channel perineurial and subperineurial glial that tile with each other loosely. **(F)** Cross-sectional view of the VNC showing the same cells as in (C) now showing the tapered projection from a (dorsal) channel perineurial glial cell that tiles with a neighbouring surface-only perineurial glial cell. **(G)** Surface view of the VNC showing polygonal-shaped surface-only subperineurial glia on the surface. **(H)** Surface view of the VNC showing polygonal-shaped surface-only subperineurial glia (cyan) on the surface and an underlying channel-only subperineurial glial cell (magenta). **(I)** Cross-sectional view of the VNC showing the same cells as in (H). **(J-K)** Surface view (J) and oblique view (K) of the VNC showing two ventral channel subperineurial glia (cyan and magenta) with long extensions towards the neuropil. **(L)** Lateral view of the VNC showing channel subperineurial glia on the ventral and dorsal surfaces, sending projections along the channels, and channel-only subperineurial glia, which reside along the channels only, both forming tube-like structures. **(M)** Cross-sectional view of the VNC showing a single cortex glial cell in the cortical region forming a membranous, honeycomb-like structure. **(N-O)** Lateral view (N) and cross-sectional view (O) of the VNC showing ensheathing glial cells at the border of the cortex and neuropil. Tract ensheathing cells extend longer or shorter processes through axon tracts entering the neuropil, while neuropil ensheathing cells only extend processes along the neuropil border. **(P)** Cross-sectional view of the VNC showing a type 1 astrocyte and a type 2 astrocyte sending processes into the neuropil. Type 1 astrocyte processes were highly ramified, whereas type 2 astrocyte processes were less ramified. Co-expression of Pros and Repo indicated astrocyte identity (insets). **(Q-S)** Cross-sectional views of the VNC showing single astrocytes in greyscale, from dorsal, lateral and ventral nucleus positions, with corresponding morphological type 1 or 2 indicated. Green marks Repo, and cyan and magenta mark MCFO clones in all panels, except (P) with Pros in magenta. Dashed lines outline the neuropil and full lines outline the VNC. Clones represent samples at 0 h after larval hatching. All scale bars represent 10 µm.

**Figure 2.**
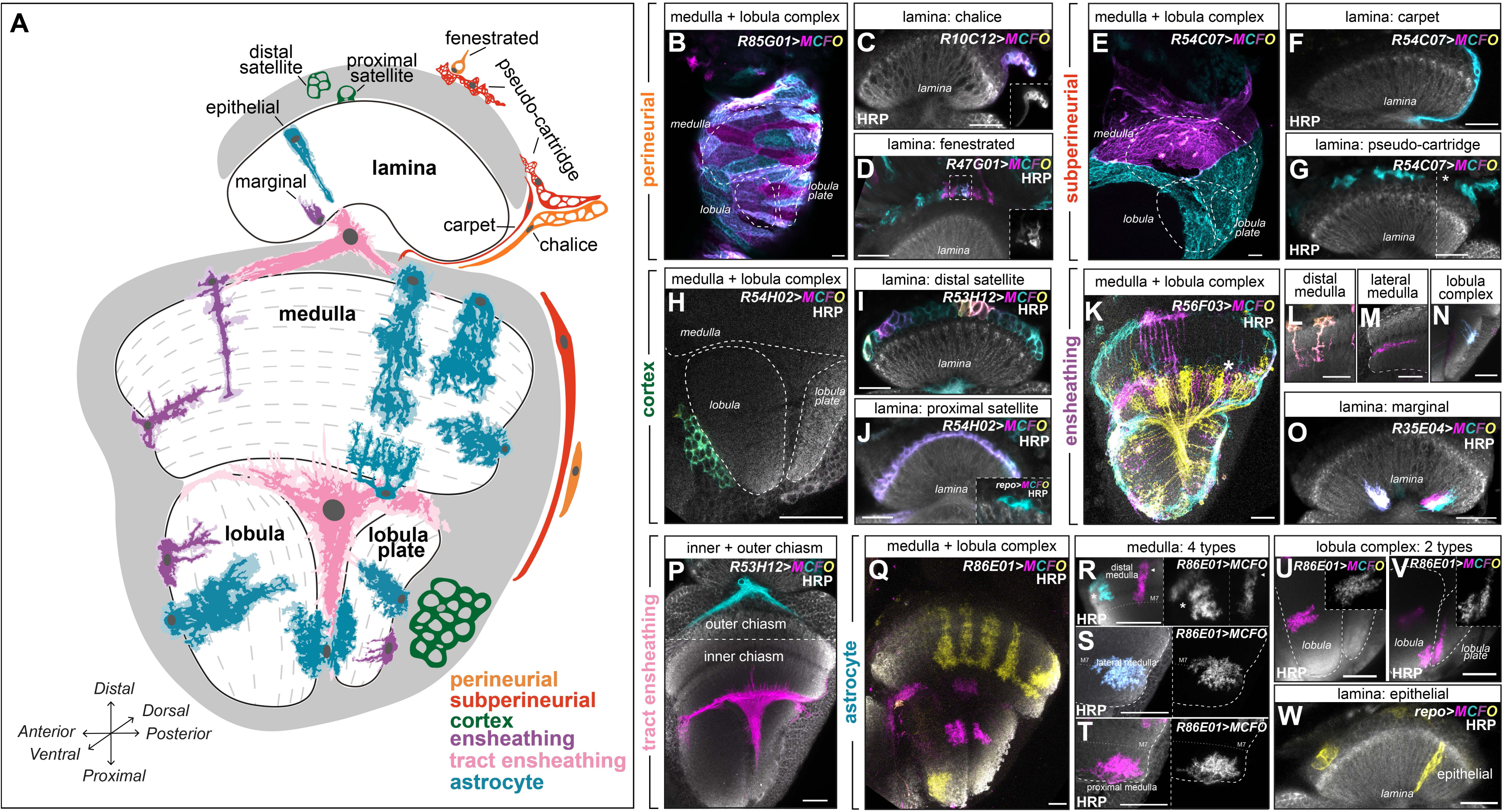
Morphologies of glia in the adult optic lobe. **(A)** Schematic of the cross-section of the adult optic lobe and its four neuropils: lamina, medulla, lobula and lobula plate with glial classes indicated. Dashed lines indicate the layers of the specific neuropil. Cortex (neuronal cell bodies) indicated in Grey. **(B)** Maximum projection showing MCFO clones of perineurial glial cells covering the medulla, lobula, and lobula plate. Cells were oblong-shaped and tiled together. **(C)** A cross-sectional view of the lamina showing a chalice glial cell (a type of lamina perineurial glia at the rim of the lamina cortex and neuropil). **(D)** A cross-sectional view of the lamina showing fenestrated glial cells (a type of lamina perineurial glia), separating the compound eye from the lamina. **(E)** Maximum projection showing MCFO clones of subperineurial glia that covered the medulla, lobula and lobula plate as large squamous cells that tiled together. **(F)** A cross-sectional view of the lamina showing a single carpet glia (a type of lamina subperineurial glia), along the rim of the lamina cortex and neuropil. **(G)** A cross-sectional view of the lamina showing pseudocartridge glia (a type of lamina subperineurial glia), as irregularly shaped cells above the lamina cortex. **(H)** A cross-sectional view of the medulla, lobula and lobula plate showing a single cortex glial cell in the lobula cortex with typical membranous and honeycomb-like morphology. **(I)** A cross-sectional view of the lamina showing distal satellite glia (a type of lamina cortex glia), excluded from the most proximal region of the lamina cortex. Note that inner chiasm glia were also labelled by this driver (bottom). **(J)** A cross-sectional view of the lamina showing proximal satellite glia in the most proximal region of the lamina. Inset shows a single proximal satellite glia labelled by *repo-Gal4*. **(K)** A cross-sectional view of the medulla, lobula and lobula plate showing MCFO clones labelling ensheathing glia and a subset of neurons (asterisk). **(L)** Examples of ensheathing glia in the distal medulla sending primary projections from the surface of the neuropil to layer M7. **(M)** An example of an ensheathing glial cell in the lateral medulla sending a primary from the outer neuropil surface inwards along layer M7. **(N)** An example of an ensheathing glial cell in the lobula plate neuropil sending several short processes with minimal secondary branches into the neuropil. **(O)** A cross-sectional view of the lamina showing marginal glia (a type of lamina-specific ensheathing glia), which project partway into the lamina neuropil from the distal neuropil surface. **(P)** A cross-sectional view of the optic lobe showing two morphologically distinct tract ensheathing glia (called chiasm glia) located in the outer chiasm between the lamina and medulla, or the inner chiasm between the medulla, lobula, and lobula plate. Outer chiasm glia did not send projections into the neuropils, but the inner chiasm glia projected into all three neuropils. The dashed line separates images in the same brain on different z-stacks. **(Q)** A cross-sectional view of the medulla, lobula and lobula plate showing astrocytes. **(R)** Examples of distal medulla astrocytes with short and long morphologies. The short distal medulla astrocyte (asterisk) projected to layer M6, while the long distal medulla astrocyte (arrowhead) projected to layer M8. **(S)** An example of a lateral medulla astrocyte projecting into the neuropil laterally along the M7/8 layers. **(T)** An example of a proximal medulla astrocyte (also called chandelier glia) projecting from the proximal surface of the medulla, up to layers M9 and M10. **(U,V)** Examples of astrocytes in the lobula and lobula plate, which **(U)** projected into the lobula neuropil from the cortex-neuropil border or **(V)** projected into both the lobula and pobula plate neuropils. **(W)** Example of an epithelial glial cell (right; the lamina astrocyte population), which projected across the entire distal-proximal neuropil length. Cyan, yellow, magenta and purple mark the multi-color flip-out (MCFO) clones, while white labels horseradish peroxidase (HRP labels the neuropils) in the main panels. Insets show the MCFO clones in greyscale. Dashed lines are used to outline neuropil borders or separate an inset. All scale bars represent 20 µm.

Here, we leverage the simple and tractable *Drosophila* nervous system to explore the relationship between glial morphological diversity and transcriptional signatures. We generated and validated a single cell atlas of all glial classes in two distinct *Drosophila* circuits— the embryonic ventral nerve cord and the adult optic lobe. We chose these circuits as they are well-characterized with many existing tools and reagents, they span two different circuit types (sensorimotor *versus* pure sensory), and they represent two distinct developmental stages. We identified several new glial morphological categories but found no clear correlation between glial morphological diversity and detectable transcriptional diversity. Moreover, we found that class-specific transcriptomes were conserved from embryo to adult, despite changes in circuit location and developmental stage. One exception was the glia of the optic lobe lamina, which accounted for the majority of glial diversity at a transcriptional level. The lamina and its associated glia lie in close proximity to an environmental interface, positioned immediately below photoreceptors of the compound eye; these glia were enriched for gene expression associated with immune-related functions. Our data suggest that within-class (*e.g.* astrocytes) glial morphological categories cannot be assumed to correspond to transcriptionally distinct subclasses. Instead, we propose that glia adopt different morphological and functional states in response to cues from their local environment.

## Results

### Morphological diversity of embryonic *Drosophila* glia

To determine the relationship between glial morphology and transcriptional signature, we began by characterising glial morphology across distinct *Drosophila* brain regions and developmental stages. We focused on glia in the ventral nerve cord (VNC), akin to the vertebrate spinal cord, in the late embryonic (stage 17) CNS. The developing VNC contains five major glial classes: astrocytes, ensheathing glia, cortex glia, and two types of surface glia, which are all neuroectodermal in origin and express the marker *reversed polarity (repo).* (Freeman, 2015). We used enhancer trap Gal4 drivers expressed in each of these glial cell classes to sparsely label cells using the multi-colour flip-out (MCFO) cassette. We then visualized single glial cell morphology at 0 hours after larval hatching (0h ALH) to assess morphological diversity both within and between classes (Figure 1). Note that in addition to the five major glial classes described above, the VNC contains a distinct class called the midline glia, which are a transient population found only during embryonic and larval stages (Perz, 1994; Sonnenfeld and Jacobs, 1995a, 1995b; Stollewerk et al., 1996). Although midline glia express *wrapper*, otherwise known as a cortex glia marker (Banerjee et al., 2017; Noordermeer et al., 1998; Stork et al., 2009), they do not resemble cortex glia in form or function but instead ensheath commissural axons and play critical roles in axon guidance and VNC morphogenesis (Jacobs, 2000). Moreover, unlike the other major glial classes described above, midline glia are mesectodermal in origin (Kosman et al., 1991; Thomas et al., 1988) and do not express the pan-glial marker *repo* or the broad glial marker *glial cells missing (gcm)* (Jacobs, 2000). Midline glia have been characterised extensively by several groups (Hartenstein, 2011; Hidalgo, 2003; Jacobs, 2000; Kearney et al., 2004; Vasenkova et al., 2006; Wheeler et al., 2006), therefore, given their distinct origin and the ambiguity surrounding their functional classification, we instead focused our analyses on *repo+* glia.

#### VNC surface glia

The *Drosophila* CNS is bathed in circulating hemolymph, which contacts the VNC at its main surface and along dorsoventral channels that perforate the VNC along the midline between pairs of longitudinal connectives and neighbouring neuromeres (Evans et al., 2010; Ito et al., 1995). To analyse surface glia morphology in the VNC, we used the *CG5080-Gal4* and *moody-Gal4* drivers to generate MCFO clones. Previous reports identified two classes of surface glia, perineurial on the outer surface of the VNC (*CG5080*+) and subperineurial glia (*moody*+) lying below perineurial glia (Freeman, 2015; Pogodalla et al., 2022). When we generated clones in the VNC with *CG5080-Gal4*, we observed two distinct perineurial glial cell morphologies. We found loosely tiled cells at the periphery of the VNC, displaying a fibrous morphology, characteristic of what has been described for perineurial glia previously (Pogodalla et al., 2022). Here, we refer to these as surface-only perineurial glia (Figure 1A-F). *CG5080-Gal4* also labelled cells located along the midline of the dorsal and ventral surfaces of the VNC, which when viewed at the surface, tiled with and resembled the fibrous morphology of surface-only perineurial glia (Figure 1B-F; N=140 clones from N=13 brains). However, in contrast to the surface-only perineurial glia, each of these cells also sent a single tapered projection inward along the dorsoventral channel (Figure 1C,D,F and Figure 1 – figure supplement 1A-C; N=51 clones from N=13 brains). Ventral cells projected further than their dorsal counterparts and were present along the entire anterior-posterior axis of the VNC, whereas cells projecting from the dorsal surface were observed with lower frequency towards more posterior positions of the VNC (Figure 1D and Figure 1 – figure supplement 1A). To confirm that these channel-associated cells belong to the perineurial glial class, we used *CG5080-Gal4* to label individual cells while co-labelling all glia with *repo-LexA>LexAop-myr::tdTomato.* We observed that the channel-associated cells occupied the outermost glial surface of the VNC (Figure 1 – figure supplement 1B,C). Taken together with the fact that these cells tiled with surface-only perineurial glia, these data argue that they are a subclass of perineurial glia. Therefore, we refer to them hereafter as ‘channel perineurial glia’. Thus, perineurial glia could be characterised into two morphologically distinct populations: the surface-only perineurial glia, which have been described previously, and the channel perineurial glia, which we describe here (Figure 1A-F and Figure 1 – figure supplement 1).

Two sub-classes of subperineurial glia were described previously based on lineage relationships, molecular markers, and nuclear positions: (i) subperineurial glia located at the main surface of the VNC (hereafter referred to as surface-only subperineurial glia) and ‘channel glia’, which line the surface of the dorsoventral channels (Beckervordersandforth et al., 2008; Ito et al., 1995). To avoid ambiguity in terminology and to distinguish channel glia from the newly identified channel perineurial glia, hereafter we refer to previously named channel glia as ‘channel subperineurial glia’. Both surface-only and channel subperineurial glia were previously reported to express *moody* (Beckervordersandforth et al., 2008), and accordingly, we recovered both surface-only and channel subperineurial glial clones with *moody-Gal4* (N=101 clones from N=12 brains) (Figure 1A, G-L). Intriguingly, although we observed some glia that were associated with the surface exclusively (69.3% of total clones; Figure 1B, H,I) and others that lined the channels exclusively (8.9% of total clones; Figure 1H,I), we also observed a continuum of glial morphologies between these two extremes (Figure 1J-L). Importantly, glia with intermediate morphologies (*i.e.* which associated with both the surface and the channels; 21.8% of total clones), tiled tightly with neighbouring surface-only subperineurial cells (Figure 1H,I). Thus, subperineurial glia could be characterised roughly into three morphologies: the surface-only subperineurial glia, the channel-only subperineurial glia those with intermediate morphologies *i.e.* the surface and channel-associated subperineurial glia (Figure 1G-L); note that hereafter we refer to the latter two morphologies collectively as ‘channel subperineurial glia’.

#### VNC cortex glia, ensheathing glia, and astrocytes

*Drosophila* have three glial cell classes that collectively perform the functions of vertebrate astrocytes: cortex glia, ensheathing glia and astrocytes (Freeman, 2015). To analyse their morphology, we used the *R54H02(wrapper fragment)-Gal4*, *Eaat2-Gal4*, and *alrm-Gal4* drivers to generate MCFO clones, respectively. In the VNC, cortex glia (*wrapper*+) varied in size but exhibited one general morphology. In brief, all cortex glia extended a large membrane sheet to envelop neighbouring neuronal cell bodies. Furthermore, each cortex glia extended processes to contact the apical and basal surfaces of the cortex, which were occupied by other glial cell membranes (Figure 1A,M; N=295 clones from N=13 brains) (Freeman, 2015).

Two morphological categories of VNC ensheathing glia have been described previously: (i) ensheathing glia associated with the ventral and medial neuropil, which wrap the border between the cortex and neuropil with no projections extending into the neuropil itself; hereafter referred to as neuropil ensheathing glia, and (ii) ensheathing glia associated with the dorsal neuropil and the proximal regions of intersegmental nerves (the latter are also known as ‘ensheathing/wrapping glia’ or ‘tract ensheathing glia’) (Kremer et al., 2017; Peco et al., 2016). We recovered both morphological categories of ensheathing glia with *Eaat2-Gal4* (79.6% of clones were neuropil-only ensheathing glia and 20.4% of clones were tract ensheathing glia; Figure 1A,N,O). We note that tract ensheathing glia varied in the degree to which they associated with the neuropil and axon tracts (See examples in Figure 1N,O; N=190 clones from N=11 brains).

Using *alrm-Gal4* we observed two morphological categories when we labelled astrocytes (*alrm*+) by MCFO (Figure 1P-S), which we named type 1 and type 2 (Figure 1A,P-S; N=117 clones from 36 brains). Type 1 astrocytes extended highly ramified processes throughout the neuropil (Figure 1A,P,Q), and represented 37 ± 6 SEM% of clones. Type 2 astrocytes elaborated some processes within the neuropil, but also extended a single radial process along the border of the neuropil and cortex (Figure 1A,P,R,S), and represented 62 ± 4 SEM % of clones. Type 2 astrocytes were distinguished from ensheathing glia by nuclear expression of Prospero (Figure 1P) (Peco et al., 2016). The VNC contains a single neuropil regionalised into dorsal motor and ventral sensory processing domains (Grueber et al., 2007; Landgraf et al., 2003; Mark et al., 2021; Merritt and Whitington, 1995; Zlatic et al., 2009). We wondered whether type 1 and type 2 astrocytes displayed regional associations that might indicate morphological specialisation to distinct circuit-types.

Indeed, Peco et al., (2016) previously reported that astrocyte cell bodies could be consistently allocated to one of three distinct regions around the neuropil: dorsal, lateral and ventral, and that the neuropil domains covered by astrocytes from these regions were also stereotyped corresponding to either the dorsal-medial, lateral or ventral-medial domains. To probe for astrocytic morphological specialisation between the dorsal (motor) and ventral (sensory) processing domains of the neuropil we used astrocyte cell body position along the neuropil to allocate them to either the dorsal, lateral or ventral domains (as in Peco et al., 2016; Figure 1 – figure supplement 2A; Figure 1 – Videos 1-3; N=117 clones from 36 brains) and correlated these positions to type 1 or type 2 morphologies (Figure 1 – figure supplement 2B). We found that astrocytes with cell body positions along the dorsal (motor) and ventral (sensory) neuropil border were predominantly type 2, whereas laterally positioned astrocytes were predominantly type 1 (lateral vs dorsal and lateral vs ventral, *p*<0.0001, dorsal vs ventral, *p*=0.593; Fisher’s exact test). Thus, type 1 and type 2 astrocytes show regional preferences, but these did not correlate with sensory versus motor circuit types. To further quantify their morphological differences, we measured total cell volume and the number of primary branches from the cell body (Figure 1 – figure supplement 2C,D). Unsurprisingly, lateral astrocytes (more ramified type 1 morphology) occupied larger volumes with more primary branches on average compared to either dorsal or ventral astrocytes (type 2 morphology) (total volume: lateral vs dorsal. *P*=0.0002, and lateral vs ventral, *p*=0.004; number of branches: lateral vs dorsal. *P*=0.0005, and lateral vs ventral, *p*=0.0097; Mann-Whitney U-test; Figure 1 – figure supplement 2C,D). However, dorsal and ventral astrocytes did not differ from each other in their cell volumes or the number of primary branches (total volume, *p*=0.101; number of branches: *p*=0.398; Mann-Whitney U-test; Figure 1 – figure supplement 2C,D).

In summary, within the early larval VNC, cortex could be defined by a single stereotyped morphology, whereas perineurial, subperineurial, ensheathing and astrocyte glial classes each contained morphologically distinct subpopulations (Figure 1 and Supplementary file 1).

### Optic lobe glia display morphological diversity within glial classes

In addition to the relatively simple VNC, we also focused on the adult optic lobe, which is more structurally complex than the VNC with four distinct neuropils called the lamina, medulla, lobula and lobula plate (Figure 2A). Several other groups have characterised the morphology of each of the five major glial classes (perineurial, subperineurial, cortex, ensheathing and astrocyte) present in the optic lobe in detail (Edwards et al., 2012; Kremer et al., 2017; Richier et al., 2017). Therefore, we used previously characterised Gal4 lines (Chotard and Salecker, 2007; Edwards et al., 2012; Kremer et al., 2017) (Figure 2 – figure supplement 1) to generate MCFO clones to visualise glial morphologies and validate morphological diversity within each class (Summarized in Figure 2; see Materials and Methods for more details).

#### Optic lobe surface glia

Briefly, we used *R85G01-Gal4* to label perineurial glia that covered the medulla, lobula, and lobula plate neuropils and observed morphological homogeneity across these three neuropils (Figure 2A, B) (Kremer et al., 2017). We used *R10C12-Gal4* to label chalice glia, a putative perineurial glial population found at the margins of the lamina neuropil (Figure 2A, C). Finally, we used *R47G01-Gal4* to label fenestrated glia, specialised perineurial glia found over the lamina cortex (Figure 2A, D) (Kremer et al., 2017). We then used *R54C07-Gal4* to label subperineurial glia of the optic lobe (Kremer et al., 2017). Subperineurial glia of the same morphology covered the medulla, lobula, and lobula plate neuropils (Figure 2A, E). In the lamina, *R54C07-Gal4* also labelled carpet glia, a specialised subperineurial glia found at the margins of the lamina neuropil (Edwards et al., 2012; Kremer et al., 2017) and pseudo-cartridge glia, a specialised subperineurial glia adjacent to the lamina cortex (Figure 2A, F,G). Thus, we validated each of the previously annotated surface glial subtypes (Edwards et al., 2012; Kremer et al., 2017).

#### Optic lobe cortex glia, ensheathing glia and astrocytes

To label optic lobe cortex glia, we used *R54H02-Gal4 (*Gal4 driven by a fragment of the *wrapper* locus) and *R53H12-Gal4* (Kremer et al., 2017). *R54H02-Gal4* labelled cortex glia in the medulla, lobula and lobula plate, which were morphologically indistinguishable across the three neuropils, and the proximal satellite glia, a lamina-specific cortex glia located in the proximal lamina (Figure 2A, H, J) (Kremer et al., 2017). *53H12-Gal4* labelled the distal satellite glia, another lamina-specific cortex glia population, located in the distal lamina (Figure 2A, J) (Kremer et al., 2017).

To visualize optic lobe ensheathing glia, we used *R56F03-Gal4, R35E04-Gal4,* and *R53H12-Gal4* (Kremer et al., 2017). *R56F03-Gal4* labelled lamina, medulla, lobula, and lobula plate ensheathing glia (Kremer et al., 2017). The medulla exhibited two ensheathing glial morphologies: one in the distal medulla, which sent a primary process to layer M7 and one in the lateral medulla which sent a primary process along layer M7 (Figure 2A, K-M)(Kremer et al., 2017). Ensheathing glia of the lobula and lobula plate showed more complex branching patterns compared to the medulla and were less stereotyped (Figure 2A, K, N). *R56F03-Gal4* also labelled the marginal glia, a lamina-specific ensheathing glia (not shown) (Kremer et al., 2017), which we labelled specifically by *R35E04-Gal4* (Figure 2A, O). Finally, *R53H12-Gal4* labelled chiasm glia, a specialised ensheathing glia (tract ensheathing glia), which ensheaths tracts of neuronal projections between neuropils. Chiasm glia displayed two distinct morphologies: one associated with the chiasm between the lamina and medulla (outer chiasm), and the other associated with the chiasm between the medulla, lobula and lobula plate chiasm (inner chiasm) (Figure 2A, P)(Edwards et al., 2012; Kremer et al., 2017).

Finally, to label optic lobe astrocytes we used *R86E01-Gal4* (Kremer et al., 2017). In the medulla, we observed four distinct astrocyte morphologies as previously reported: two in the distal medulla, a third in the lateral medulla, and a fourth in the proximal medulla (also called chandelier glia) (Figure 2A, Q-T)(Edwards et al., 2012; Kremer et al., 2017; Richier et al., 2017). In addition, the lobula and lobula plate were together populated by three morphologically distinct astrocyte populations (Figure 2A, Q, U, V). Finally, *R86E01-*Gal4 also labelled astrocytes of the lamina, called epithelial glia (not shown; shown instead with *repo-Gal4*) (Figure 2A, W).

Since astrocytes displayed the most within-class morphological diversity of all the glial classes in the optic lobe, often with multiple stereotyped morphologies occupying the same neuropil, we sought to characterise them further. The medulla, lobula and lobula plate neuropils display synaptic stratifications (layers), which arise because of diverse neuronal arborization patterns (Fischbach and Dittrich, 1989). Different neurons project to different layers and in this way restrict the partners with whom they synapse in a layer-specific manner. Thus, distinct layers encode distinct features of visual information (Fischbach and Dittrich, 1989). We found that astrocytes which differed dramatically in shape sometimes had similar whole cell volumes and similar numbers of primary branches (Figure 2 – figure supplement 2A-C). Therefore, to better assess whether astrocyte morphological categories displayed layer-specific and, therefore, circuit-specific associations, we quantified their volumes across neuropil layers (Figure 2 – figure supplement 2D,E). This analysis revealed that each astrocyte morphological category displayed a clear preference in the layers it covered (Figure 2 – figure supplement 2D,E; Figure 2 - Videos 1-8).

In sum, all major classes of optic lobe glia showed some regionalized morphological differences (Figure 2A and Supplementary file 2). Additionally, for some glial classes — cortex glia in the lamina, ensheathing glia in the medulla, and astrocytes in the medulla, lobula and lobula plate — we observed morphologically distinct populations in close proximity to each other within the same neuropil. Thus, optic lobe glia exhibited much more morphological diversity than VNC glia, prompting us to ask how transcriptional diversity differs between glia at embryonic and adult stages. Furthermore, are distinct glial morphologies within broad glial classes associated with unique transcriptional signatures?

### A transcriptomic atlas of embryonic and young adult Drosophila glia

To correlate glial cell morphology and transcriptional identity during development, we performed scRNA-seq on late-stage *Drosophila* embryos (stage 17). Whole embryos were dissociated into a single cell suspension, filtered, prepared using the 10X Genomics single cell pipeline, and sequenced via Illumina sequencing (see Materials and Methods). At this stage and through larval stage 1, we quantified an average of 528 ± 9 SEM glial cells in the CNS (labelled by the glial-specific marker Reversed polarity, or Repo) that were localized in the VNC, and 188 ± 9 SEM glial cells in the brain lobes (N=8 brains per anatomical region). Thus, our dataset was enriched for VNC glia, which make up 74% of the CNS glia at this time (p<.0001, one-way ANOVA). Following sequencing, cells were clustered based on differential gene expression using Seurat, and glial clusters were computationally isolated by expression of the pan-glial markers *repo,* and the astrocyte-specific markers *GABA transporter (Gat)* and *astrocytic leucine-rich repeat molecule (alrm)*, except for the midline glia, which lack *repo* expression (Jacobs, 2000) (see Materials and Methods and Figure 3 for further details). We bioinformatically isolated midline glia based on *wrapper* and *single minded (sim)* expression (Crews et al., 1988; Noordermeer et al., 1998) and performed hierarchical cluster analysis on midline glia, all *repo+* glial clusters and neuronal clusters (see Materials and Methods and Figure 3 – figure supplement 1A-F for more details). This analysis revealed that midline glia formed an outgroup to neuronal and *repo+* glial clusters; perhaps not surprisingly given their distinct (mesectodermal) origin (Kosman et al., 1991; Thomas et al., 1988) (See Source data file 1 for genes with enriched expression in midline glia). We therefore continued all further analyses focusing on *repo+* glia only but nonetheless validated *wrapper* expression in midline glia *in vivo* using *wrapper-Gal4* (carried on a BAC insertion; Banerjee et al., 2017) (Figure 3 – figure supplement 1G,H).

**Figure 3.**
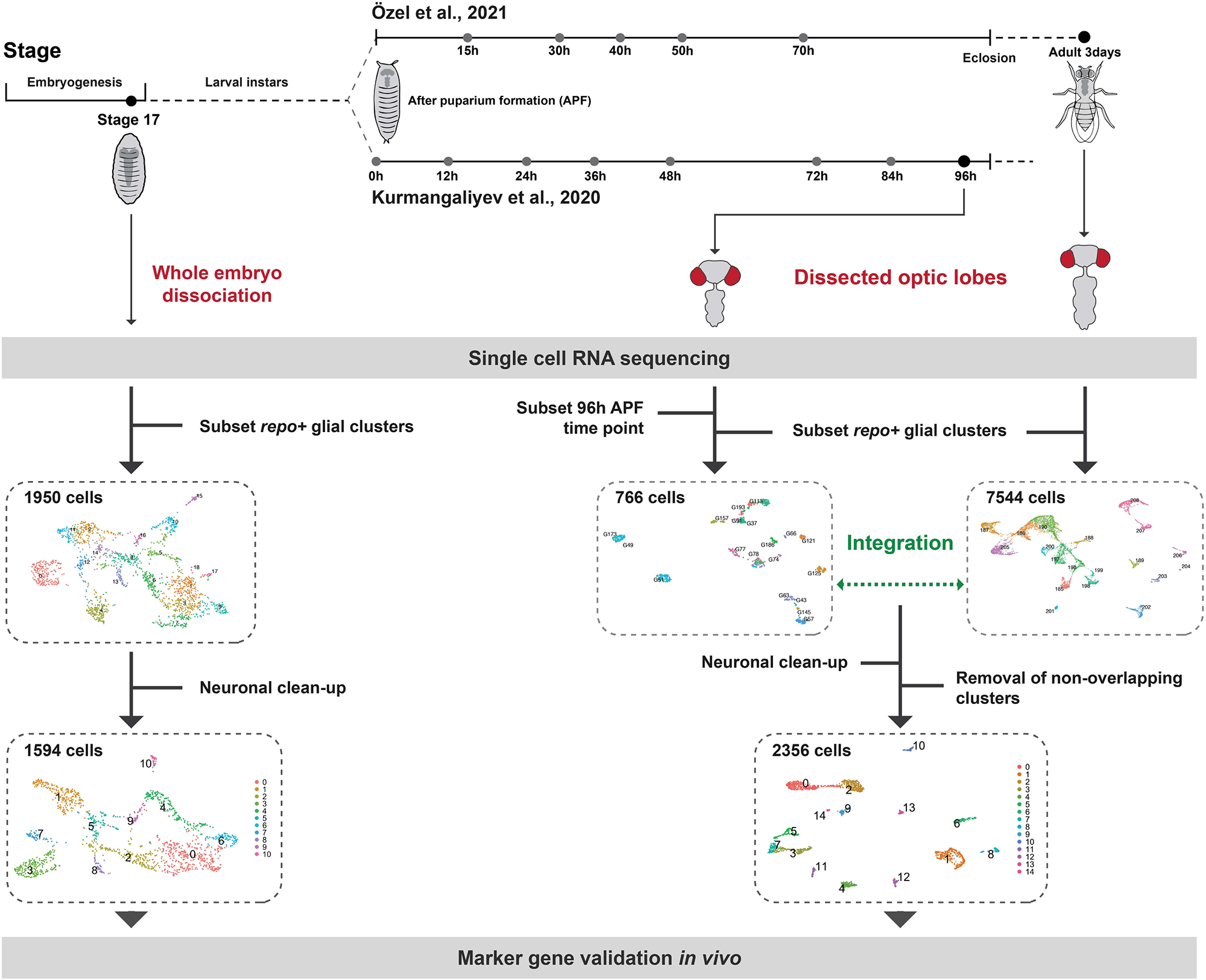
Summary of the experimental and computational workflow used in this study. **(Left)** We performed scRNA-seq on whole stage 17 embryos. We then subsetted *repo+* glial cells and eliminated neuronal contamination (see Materials and Methods). This resulted in 1594 cells and 11 clusters, which we proceeded to annotate by validating marker gene expression *in vivo*. **(Right)** We isolated and integrated *repo+* glial clusters from two published datasets of dissected optic lobes (Kurmangaliyev et al., 2020; Özel et al., 2021)(see also Figure 3 – figure supplement 1). We eliminated neuronal contamination and clusters that contained cells from only the Özel et al. dataset (see Materials and Methods) (see also Figure 3 – figure supplements 2 and 3). This resulted in 2356 cells and 15 glial clusters, which we went on to annotate by comparisons with cell-type-specific bulk RNA sequencing datasets and by validating marker gene expression *in vivo*.

For the optic lobes, we used data from two recent studies that performed scRNA-seq on whole optic lobe tissue at the young adult stage and throughout pupal development (Kurmangaliyev et al., 2020; Özel et al., 2021). Despite slight differences in experimental approaches between these studies (see Figure 3 for summary), both analyses generated 19 glial clusters for the young adult optic lobes (Figure 3– figure supplement 2A,B); however, since both studies focused on neuronal development, glial clusters were not annotated or analysed (Kurmangaliyev et al., 2020; Özel et al., 2021). Since glial cells constitute only 10-15% of neural cells in the *Drosophila* nervous system (Kremer et al., 2017), we sought to increase the number of cells analysed to maximise our ability to identify rare cell types. Therefore, we isolated glial cells based on their original annotation in each study (*i.e. repo* expression) and then combined the closest developmental stages using the Seurat integration pipeline to remove batch effects between libraries (Figure 3 and Materials and Methods). Here, we focused primarily on the 3-day-old adult dataset from Özel et al. (2020) (7544 cells) integrated with the 96 hours after puparium formation (APF) dataset from Kurmangaliyev et al. (2020) (766 cells); hereafter referred to as the young adult dataset (Figure 3 and Figure 3– figure supplement 2C,D).

On Uniform Manifold Approximation and Projection (UMAP) visualisations, we noticed that many clusters were connected by streams of cells, which expressed *Resistant to dieldrin (Rdl), Frequenin 1 (Frq1),* and *Nckx30C* (Figure 3 – figure supplement 3A, top). These streams also co-expressed *embryonic lethal abnormal vision (elav)* and *found in neurons (fne),* whose expression is known to be enriched pan-neuronally (Figure 3 – figure supplement 3A, bottom). Indeed, on UMAP visualisations *Rdl, Frq1* and *Nckx30C* were expressed pan-neuronally (Figure 3– figure supplement 3B). We also detected *elav, fne, Rdl, Frq1,* and *Nckx30C* co-expressed in a subset of the embryonic dataset (Figure 3– figure supplement 3C,D). These data suggested that the *Rdl, Frq1* and *Nckx30C* expressing glial cells could represent neuronal contamination. Therefore, we used *in situ* hybridization chain reaction (HCR) and MCFO clonal analyses to assess the expression of these genes in adult optic lobes and newly-hatched larval VNCs, respectively, but failed to detect any expression in glial cells (Figure 3– figure supplement 4A-D). As glia interact closely with neurons throughout life, we hypothesised that the cells in our datasets that co-express *Rdl, Frq1* and *Nckx30C* represented either glial cells contaminated by neuronal transcripts or neuronal cells contaminated by glial transcripts. Therefore, to circumvent potential clustering artefacts, we removed cells expressing high levels of *Rdl, Frq1* and *Nckx30C* (see Materials and Methods; Figure 3– figure supplement 4E-H,J-N). In addition, we noticed that a few cells expressed high levels of *Hemolectin* (*Hml*), a hemocyte-specific marker (Goto et al., 2003), which likely indicated contamination from a few stray hemocytes; therefore, we eliminated these cells also (see Materials and Methods). Lastly, others have reported that glial clustering is sensitive to batch effects (Kurmangaliyev et al., 2020; Özel et al., 2021; Simon and Konstantinides, 2021), and to further minimise these, we eliminated clusters to which the Kurmangaliyev dataset contributed fewer than 1% of the total number of cells in the cluster (Figure 3– figure supplement 4I). Following this elimination and reclustering (see Materials and Methods) our datasets clustered into 11 embryonic glial cell clusters and 15 adult optic lobe glial clusters (Figure 3, Figure 3– figure supplement 4N and Figure 5– figure supplement 1A).

### Annotating embryonic glial clusters by validating marker gene expression *in vivo*

Based on known marker genes, we were able to make predictions about the identity of most of the embryonic clusters (Figure 4A,B). For example, cells in cluster 1 strongly expressed the gene *wrapper*, which is a known cell type-specific marker of cortex glia (Coutinho-Budd et al., 2017). To validate the identity of each embryonic glial cluster, we acquired Gal4 lines for marker genes that were significantly enriched in one or more clusters (Figure 4A,B, see Materials and Methods for complete list) and used these lines to generate MCFO clones (Figure 4C-H and Figure 4– figure supplements 1-3). Embryos were heatshocked between 6-10h after egg laying (prior to gliogenesis (Hartenstein et al., 1998)), and larvae were dissected at 0 h after larval hatching. These MCFO analyses revealed near perfect specificity for the predicted glial cell type (Figure 4B and Figure 4– figure supplements 1-3). Indeed, by morphology and marker gene expression, 100% of *CG6126-Gal4* MCFO clones and 17.9 ± 1.8 SEM% of *pippin-Gal4* MCFO clones were perineurial glia (see below for further resolution of perineurial glial clusters). 100% of *CG10702-Gal4* MCFO clones, 80 ± 8.9 SEM% *Ntan1-Gal4* MCFO clones, and 100% of *moody-Gal4* MCFO clones were exclusive to subperineurial glia. Furthermore, 98 ± 1 SEM% of *Eaat2-Gal4* MCFO clones were ensheathing glia, 96 ± 1.3 SEM% of *R54H02(wrapper fragment)-Gal4* MCFO clones were cortex glia, and 98 ± 1.5 SEM% of *alrm-Gal4* MCFO clones were astrocytes, consistent with their expression in the scRNA-seq data (Figure 4 and Figure 4– figure supplements 1-3). Importantly, drivers inserted in genes that showed expression across multiple clusters always produced MCFO clones with morphologies that matched the predicted cluster identities. In other words, a gene that showed expression in both the predicted astrocyte and ensheathing glial clusters produced both astrocyte and ensheathing glia clones (Figure 4 – figure supplements 1-3). In this way, we annotated 6 of the 11 clusters as surface-only perineurial glia, channel perineurial glia, subperineurial glia (including both surface-only and channel subperineurial glia), cortex glia, ensheathing glia (including both neuropil ensheathing and tract ensheathing) and astrocytes, and uncovered novel marker genes for the major glial classes (and subclasses; summarised in Figure 4B). *Lobe (L)*, a known marker of peripheral nervous system glia (Lassetter et al., 2021), was expressed by three of the remaining unannotated clusters (clusters #0, #6 and #8). We therefore annotated these clusters as peripheral nervous system glia (PNSg_1, 2 and 3) but did not validate markers for these *in vivo*. In this way we extended previous single cell atlases, which included glial cells of the larval CNS and adult VNC (Allen et al., 2020; Corrales et al., 2022), by validating marker gene expression *in vivo* and resolving surface glial classes into perineurial (surface-only and channel) and subperineurial classes, which were indistinguishable in previous datasets.

**Figure 4.**
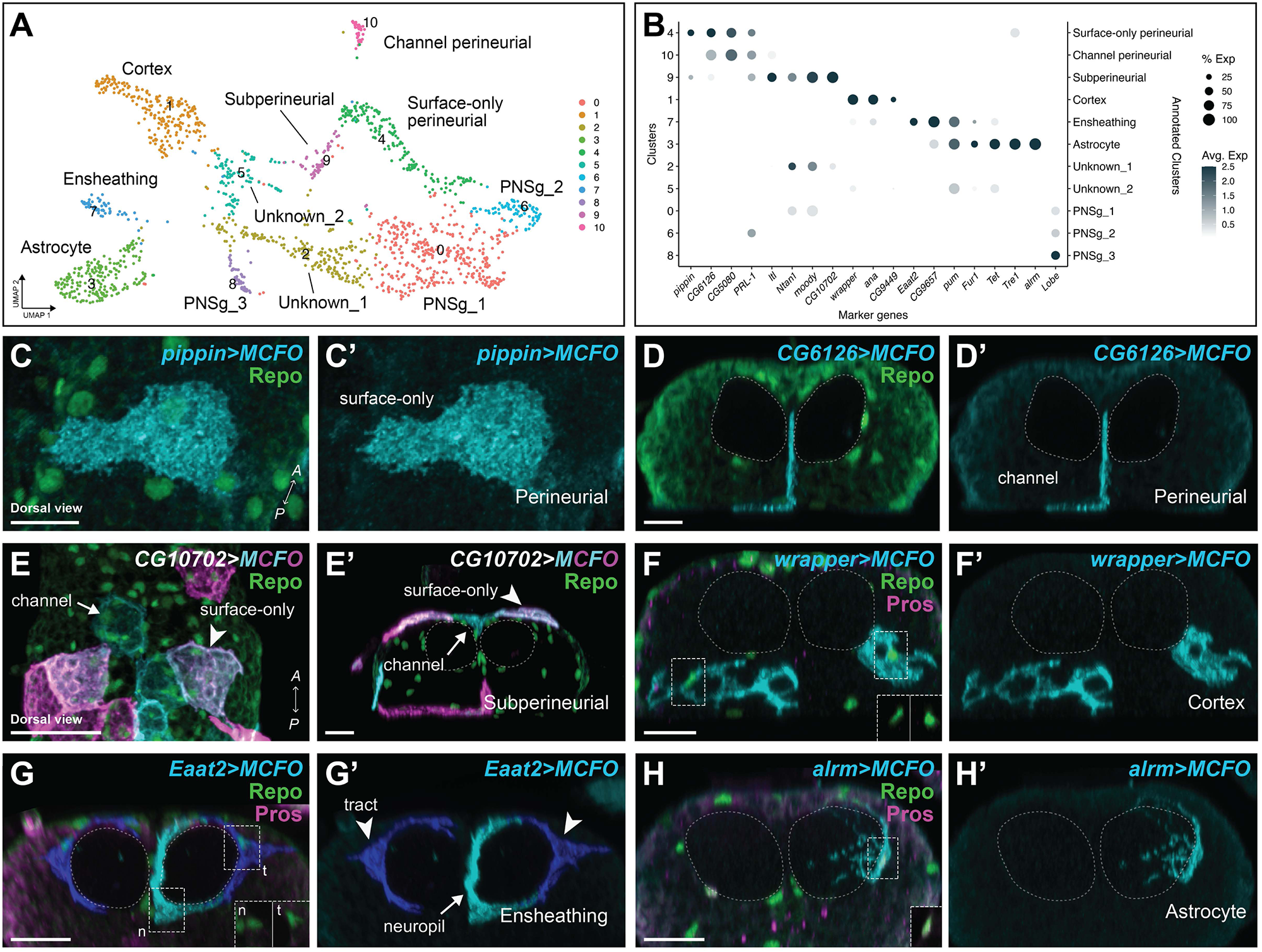
Annotation of embryonic glial clusters in the VNC. **(A)** UMAP of the 11 embryonic glial clusters labelled with both the cluster number (left) and our annotation of the specific glial class (and subtype; right) based on marker gene validation. **(B)** Expression plot of all marker genes selected for annotation, which were validated *in vivo* (except for *Lobe)*. The size of the dot represents the percentage of cells with expression in each cluster, while the colour of the dot represents the level of average expression in the cluster. **(C-H)** MCFO clones (0 h after larval hatching) generated with the Gal4 lines indicated belonged to the main embryonic glial subtypes as accessed by their morphology. **(C)** Surface-only perineurial (N=269 clones from N=11 brains), **(D)** the newly defined perineurial glial subtype, termed channel perineurial glia (N=51 clones from N=13 brains), **(E)** subperineurial, both channel and surface-only (N=95 clones from N=12 brains), **(F)** cortex (N=295 clones from N=13 brains), **(G)** ensheathing, both tract (t) and neuropil (n) (N=190 clones from N=11 brains) and **(H)** astrocyte (N=117 clones from N=36 brains). All clones labelled in cyan, with Repo in green and Prospero in magenta. Insets in (F-H) show Prospero and Repo in glial nuclei in *alrm>MCFO* clones in (H) were positive for Prospero. See Figure 4– figure supplements 1-3 for additional *in vivo* marker gene validation. Dashed lines outline the neuropils. Prime panels show MCFO clones alone. Scale bars are 10 µm.

**Figure 5.**
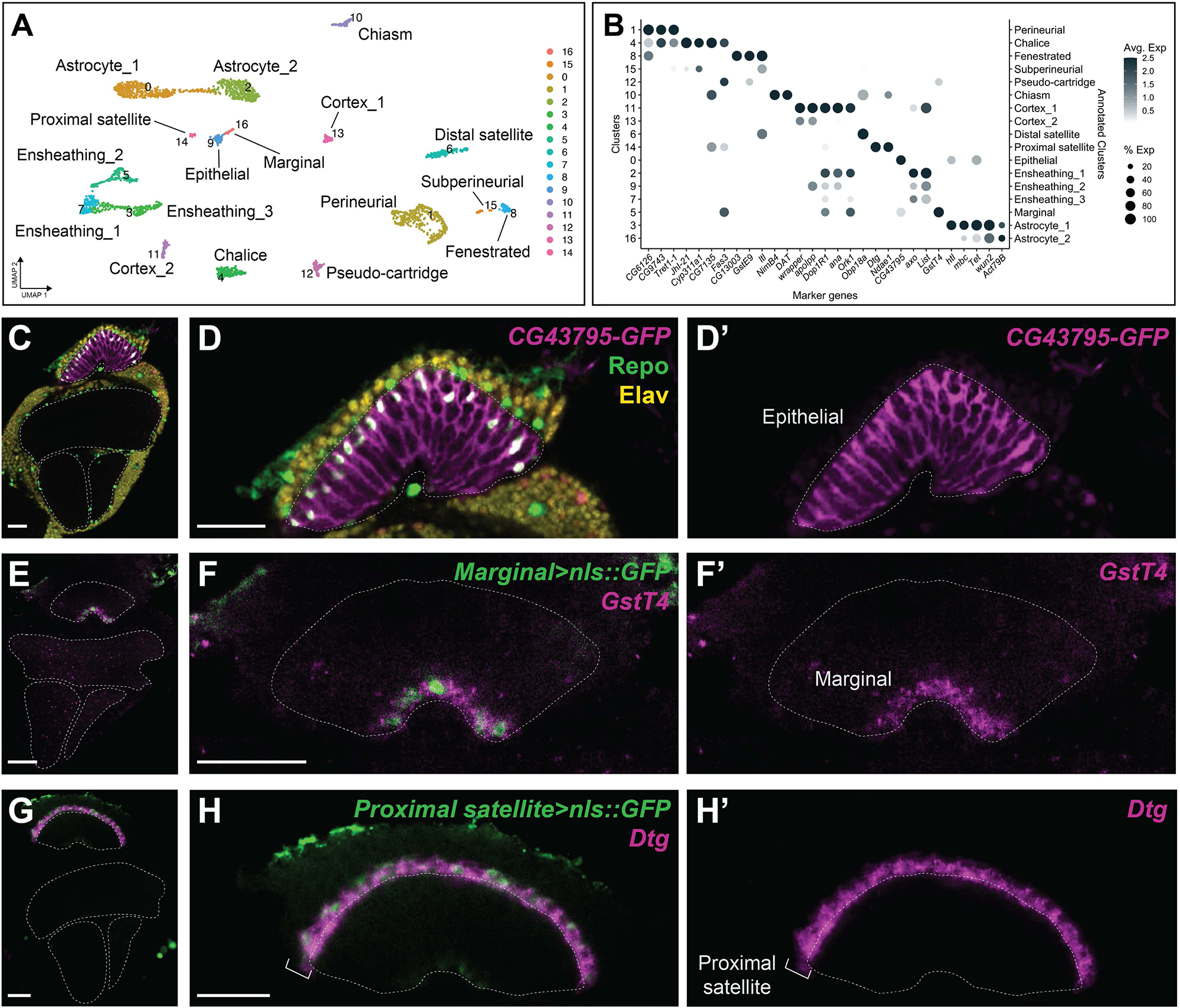
Annotation of young adult optic lobe glial clusters. **(A)** UMAP of the 17 clusters obtained for the young adult optic lobe dataset, labelled with both the cluster number (left) and our annotation of the specific glial class based on marker gene expression validated *in vivo (*right). **(B)** Expression plot of all marker genes selected for annotation and validated *in vivo*. The size of the dot represents the percentage of cells with expression in each cluster, while the colour of the dot represents the level of average expression in the cluster. **(C-H)** Three examples of marker gene validation for the whole optic lobe, focussed on the lamina: **(C,D)** *CG43795* gene trap drove GFP expression (magenta) in epithelial glia specifically. Repo in green and Elav in yellow. **(E, F)** *GstT4* and **(G, H)** *Dtg* (both in magenta) expression were detected by *in situ* HCR in marginal and proximal satellite glia, respectively. GFP was driven by the indicated glial-Gal4 in green. See Figure 5– figure supplements 1,3-5 for additional *in vivo* marker gene validation. Dashed lines outline the neuropils. Single focal planes in (C-H). Scale bars are 20 µm in (C,E,G) and 5 µm in (D,F,H).

#### Perineurial glia morphologies were transcriptionally distinct

Our previous clonal analysis unveiled two perineurial glia morphologies: surface-only perineurial glia and channel perineurial glia (Figure 1A-F and Figure 1 – figure supplement 1). Validating cluster marker expression *in vivo* revealed that surface-only and channel perineurial glia were transcriptionally distinct, with cluster #4 corresponding to surface-only perineurial glia and cluster #10 corresponding to channel perineurial glia. Clusters #4 and #10 were located adjacent to each other on the UMAP. *PRL-1* and *pippin* were enriched in cluster #4 (Figure 4B) and Gal4 drivers for these markers predominantly labelled clones with surface-only perineurial glial morphology (84.6% of 16 brains contained exclusively surface-only perineurial glia clones labelled by *PRL-1-Gal4*; 87.5% of 13 brains contained exclusively surface-only perineurial glia clones labelled by *pippin-Gal4*; Figure 4B,C and Figure 4 – figure supplement 1C,D,J). By contrast, cluster #10 expressed high levels of *CG6126* and *CG5080*, but low expression of *PRL-1* and *pippin* (Figure 4B). Consistent with these expression patterns, 100% of brains (N=11 brains) contained both surface-only and channel perineurial glia clones labelled by *CG6126-Gal4*, and 92.9% of brains (N=13 brains) contained both surface-only and channel perineurial glia clones labelled by *CG5080-Gal4* (Figures 1B-F, Figure 4D and Figure 4 – figure supplement 1J). Thus, the two perineurial glia morphologies are transcriptionally distinct.

#### Transcriptional profiles for subperineurial glia, ensheathing glia and astrocytes did not distinguish morphological categories

Our prior clonal analyses revealed morphological heterogeneity in the subperineurial glia population, along a continuum from surface-only to channel-only (Figure 1H-L), as well as two morphological categories within ensheathing glia: neuropil-only and tract-associated (Figure 1N,O), and two morphological categories within astrocytes: type 1 and type 2 (Figure 1P-S). Interestingly, while our annotations revealed separate transcriptional clusters corresponding to surface-only perineurial glia and channel perineurial glia (Figure 4A-D and Figure 4– figure supplement 1), we could resolve just one transcriptional cluster for subperineurial glia based on strong expression of *CG10702, Ntan1* and *moody* (cluster# 9; Figure 4B,E), one transcriptional cluster for ensheathing glia based on strong expression of *Eaat2* (cluster #7; Figure 1N,O and Figure 4B,G), and one transcriptional cluster for astrocytes based on strong expression of *alrm* (cluster #3; Figure 1P-S, Figure 4B). We questioned whether any of the markers enriched in our astrocyte cluster might distinguish type 1 from type 2 astrocytes. To this end, we generated MCFO clones under the control of enhancers for genes expressed in the astrocyte cluster. All drivers gave rise to clones containing both type 1 and type 2 astrocytes at the expected proportions based on our *alrm* MCFO study (∼38% type 1 and ∼62% type 2), with the exception of *pum*, which gave a slightly higher proportion of type 1 astrocytes (52%), but still contained both clones (Figure 4A,B,H and Figure 4 – figure supplements 2,3). Similarly, we failed to identify any markers that distinguished the subperineurial glial morphologies or the ensheathing glial morphologies (Figure 4E,G and Figure 4 – figure supplements 1-S3). Thus, although distinct perineurial glia morphologies corresponded to distinct transcriptional signatures, the same was not true for subperineurial, ensheathing, or astrocyte morphologies, with the caveat that scRNA-seq may fail to detect genes that are expressed at low levels.

Overall, these data suggest that morphological diversity cannot be equated with transcriptional diversity, at least at present levels of detection. As the developing *Drosophila* VNC contains a single, simple, neuropil, which may not accurately represent the diversity of more complex brain regions, we next turned to the more complex adult *Drosophila* optic lobe to assess how accurately cellular identity can be gauged by glial morphology.

### Annotating young adult optic lobe glial clusters

#### Comparisons to glial cell-type specific bulk RNA sequencing datasets

To annotate adult optic lobe glia, we compared glial cell-type specific transcriptomes (obtained from bulk-RNA sequencing of FACS-purified glial-types), which were published for the proximal satellite, epithelial and marginal glia (Davis et al., 2020), to the integrated young adult optic lobe scRNA-seq dataset. This approach matched the proximal satellite glia with cluster #14 (Pearson correlation = 0.243; Figure 5– figure supplement 1A,C), however, both the epithelial and marginal glia showed the highest Pearson correlation with cluster #9 (0.277 and 0.3, respectively; Figure 5– figure supplement 1A,D,E), a small cluster made up of only 82 cells. Since the clustering algorithm we used has a known tendency to group together rare cell types while splitting apart abundant cell-types artificially, we hypothesized that clusters containing few cells may contain more than one rare cell-type (Lancichinetti and Fortunato, 2011; Simon and Konstantinides, 2021). To determine if cluster #9 was comprised of a mixture of cells belonging to epithelial and marginal glial cell types that were artificially merged because of their rarity, we analysed cluster #9 in isolation and found that it could be divided into two distinct subclusters (Figure 5– figure supplement 1F). We identified 22 genes that were differentially expressed (by at least 4-fold) between the two subclusters (Figure 5– figure supplement 2). We found that marginal-and epithelial-specific marker genes, identified from the FACS-purified transcriptomes, segregated perfectly between the two subclusters (*e.g. GstT4* for marginal glia and *CG43795* for epithelial glia; Figure 5– figure supplement 1G,H). Indeed, when we plotted the expression of these marker genes on the original UMAP, we observed a clear spatial segregation among the cells of cluster #9, supporting the hypothesis that cluster #9 contained a heterogeneous cell population made up of both epithelial and marginal glial cells (Figure 5– figure supplement 1I,J). Therefore, we used the subclusters to manually divide cluster #9 into two clusters (renamed cluster #9 and cluster #16), with (new) cluster #9 likely corresponding to the epithelial glia and cluster #16 likely corresponding to the marginal glia (Figure 5– figure supplement 1K).

To test for heterogeneity in other clusters, we subclustered each in isolation and examined the differential gene expression between subclusters. To ensure that any subclusters uncovered in this manner represented real cellular heterogeneity rather than artificial differences due to over-clustering, we examined the number of genes expressed differentially between them (see Materials and Methods and Figure 5– figure supplement 2). As with cluster #9, cluster #8 could also be divided into two subclusters, which segregated on the main UMAP. Therefore, we manually divided it (renamed cluster #8 and cluster #15) (Figure 5– figure supplement 1L-Q). All other subclusters were deemed to be products of over-clustering artefacts, as few genes were expressed differentially between them (Figure 5– figure supplement 2). Clusters #11, #12, #14, #15 and #16 contained fewer than 3 cells belonging to either the Özel or Kurmangaliyev datasets, rendering integration, and therefore subclustering analysis, impossible.

#### Validating marker gene expression in vivo

Next, we identified marker genes enriched in each cluster that would enable us to annotate all the clusters exhaustively (see Materials and Methods and Figure 5B). We validated the expression of 31 marker genes *in vivo* using available enhancer trap lines, antibodies, and by *in situ* hybridization chain reaction (HCR) (Figure 5C-H and Figure 5– figure supplements 1,3-5). Since whole cell morphologies were not always visible when transcripts were visualized by HCR, we validated marker gene expression in specific glial-types by visualising transcript expression by HCR with glial-type Gal4 lines driving GFP (Figure 2– figure supplement 1). In this way, we annotated all 17 clusters with 13 unique glial cell identities — fenestrated glia (lamina-specific perineurial), pseudocartridge glia (lamina-specific subperineurial), chalice glia (lamina-specific perineurial), distal satellite glia (lamina-specific cortex), proximal satellite glia (lamina-specific cortex), epithelial glia (lamina-specific astrocyte), marginal glia (lamina-specific ensheathing), chiasm glia (tract ensheathing), and medulla, lobula and lobula plate perineurial, subperineurial, cortex, ensheathing and astrocyte glia — with some glial identities mapping to multiple clusters (Figures 5C-H, Figure 5– figure supplements 3-5). Hereon, non-lamina glial classes are referred to as general perineurial, general subperineurial, general cortex, general ensheathing and general astrocyte glia.

We confirmed that clusters #9, #16 and #14 corresponded to epithelial, marginal, and proximal satellite glia, respectively, consistent with our previous comparisons to the glial cell-type specific transcriptomes (Figure 5B-H, Figure 6B and Figure 5– figure supplements 4J and 5G,H,K). Carpet glia did not appear to be represented as a unique cluster, indicating that either they are transcriptionally indistinguishable from another type of surface glia or that they were not sampled in these datasets. The latter is more likely since only two carpet glia are present in each optic lobe and they can be uniquely labelled with their own driver lines (Ho et al., 2019), suggesting that they are a distinct cell type. As well, inner and outer chiasm glia could not be resolved into distinct clusters though unique driver lines do distinguish between them (Edwards et al., 2012). Instead, both mapped to cluster #10. Given that these are also rare cells, it is possible that cluster #10 contains a heterogeneous population, but that insufficient cells were sampled to resolve them in the present dataset. Thus, our annotations revealed that morphological categories associated with the lamina neuropil across all glial classes were transcriptionally distinct.

**Figure 6.**
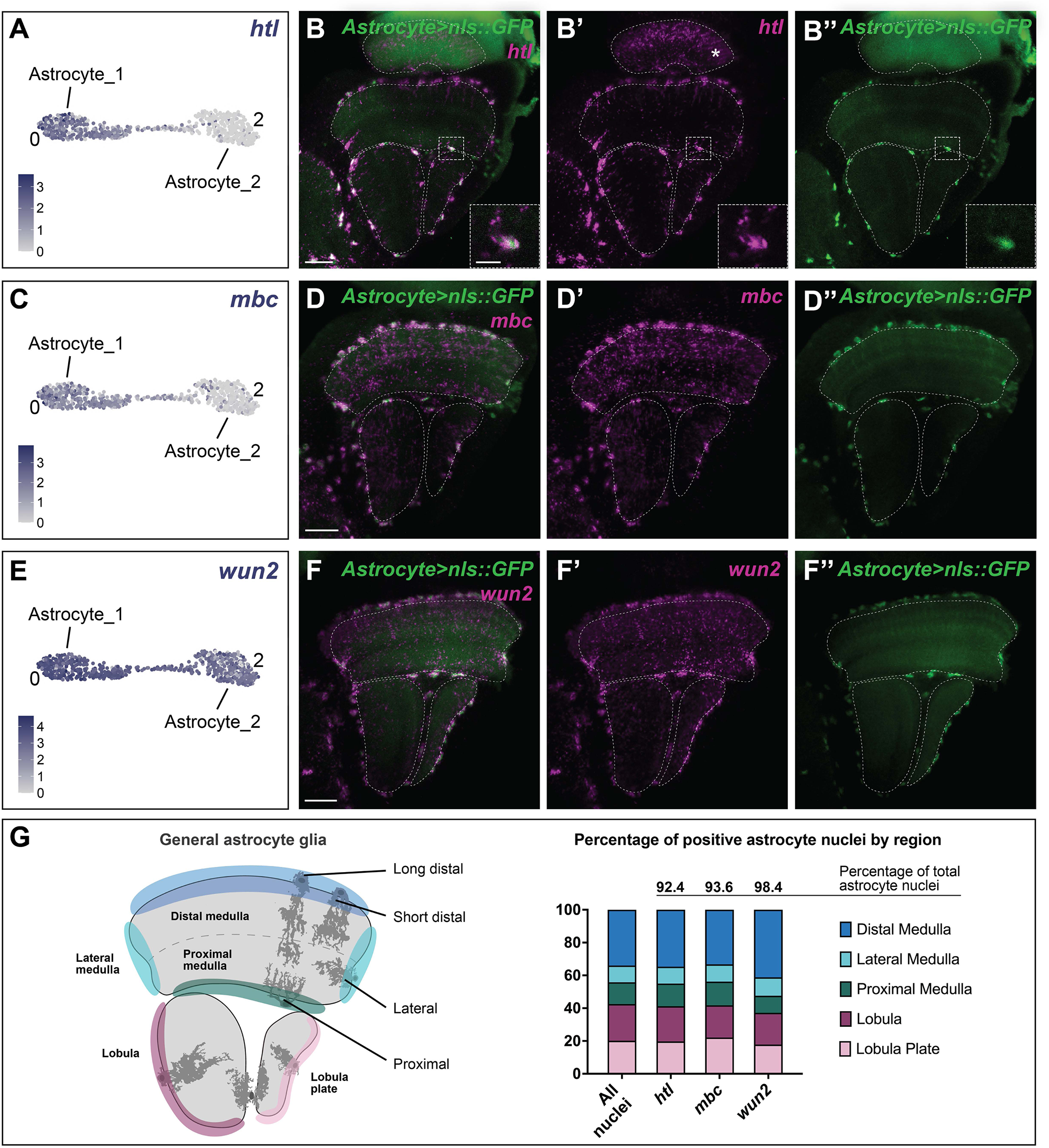
Only one cluster within the astrocyte cluster multiplet corresponded to homeostatic astrocytes detected *in vivo*. **(A)** *htl* expression levels plotted on a UMAP showing only cluster #0 (Astrocyte_1) and #2 (Astrocyte_2). Each dot represents a single cell, and the colour represents the level of expression as indicated. **(B)** *Astrocyte(R86E01)>nls::GFP* adult optic lobe in which all astrocyte nuclei were labelled with GFP and where *htl* expression (magenta) was detected by *in situ* HCR in most GFP positive nuclei. **(C)** UMAP of clusters #0 and #2 showing *mbc* expression. **(D)** *Astrocyte(R86E01)>nls::GFP* adult optic lobe where *mbc* expression (magenta) was detected by *in situ* HCR in most GFP (green) positive nuclei. **(E)** UMAP of cluster #0 and #2 showing *wun2* expression. **(F)** Maximum projection of *Astrocyte(R86E01)>nls::GFP* adult optic lobe where *wun2* expression (magenta) was detected by *in situ* HCR in most GFP (green) positive astrocyte nuclei of the medulla, lobula and lobula plate. Dashed lines outline the neuropils and scale bars are 20 µm in (B,D,F). **(G)**. Quantification of *htl, mbc* and *wun2* positive (medulla, lobula and lobula plate nuclei) astrocyte nuclei (*i.e.* GFP positive nuclei) from (B,D,F). More than 90% of astrocyte nuclei were positive for the three genes. Different regions of the medulla, lobula and lobula plate were defined, as these captured some differences in astrocyte morphotypes. The percentage of positive nuclei within each region was quantified and no significant difference (Chi-square test; *htl p*=.90, *mbc p*=.89, *wun2 p*=.33) was observed between the distribution of positive nuclei and the distribution of all nuclei in all five regions.

### General optic lobe glia do not subcluster by neuropil location

While astrocytes could be categorised into eight distinct morphologies distributed across and within neuropils (including epithelial glia of the lamina) (Figure 2A, Q-W), we only identified three transcriptional clusters of putative astrocytes. We annotated cluster #9 as the epithelial glia (astrocytes of the lamina), whereas clusters #0 and #2 co-expressed well-known astrocyte markers including *Glutamine synthetase 2* (*Gs2*), *ebony* (*e*), *nazgul* (*naz*), *alrm*, and *Eaat1* (and did not express *Eaat2,* a known ensheathing glia marker; Figure 6-supplementary S1). We wondered whether Clusters #0 and #2, which were connected to each other on the UMAP visualization, corresponded to different morphological subtypes of astrocytes or to astrocytes of different neuropils or neuropil regions. However, most differentially expressed marker genes showed enriched expression in cluster #0 compared to cluster #2; *e.g. heartless (htl),* which encodes an Fibroblast Growth Factor receptor and *myoblast city (mbc)* were expressed in cluster #0 only. We could not find unique marker genes for cluster #2 (Figure 6A,C). Therefore, we examined the distribution of astrocytes across and within neuropils (excluding the lamina) using a pan-astrocyte-specific Gal4 to drive GFP expression (*R86E01>GFP*). Since *htl* and *mbc* were expressed only in cluster #0 and not cluster #2, we then compared the distribution of GFP-positive astrocytes by region (as indicated in Figure 6G) to the distribution of *htl* expressing astrocytes. Surprisingly, although *htl* and *mbc* were expressed only in one of the two astrocyte clusters, we detected their expression in 92.4% and 93.6% of all astrocytes, respectively, and there was no bias in their regional distribution (Figure 6A-D,G). The same was true for *wunen 2 (wun 2), a* marker expressed across both clusters #0 and #2 (Figure 6E,F,G). Therefore, both clusters mapped indistinguishably to general astrocytes, with no apparent correspondence to neuropil or morphology. Similar to clusters #0 and #2, when we validated marker gene expression *in vivo,* we were also unable to distinguish between clusters #3, #5 and #7, which all appeared to correspond to general ensheathing glia (*i.e.* ensheathing glia of the medulla, lobula and lobula plate; Figure 5– figure supplement 4G-J). Likewise, clusters #11 and #13 both mapped to general cortex glia (*i.e.* cortex glia in the optic lobe excluding the lamina; Figure 5– figure supplement 3B-D). Thus, transcriptional heterogeneity of the general optic lobe glia is not regionally defined.

### Transcriptional diversity in the general optic lobe glia reflects cellular state and not cellular identity

What then is the distinction between these clusters? Across model systems, glia appear to be more susceptible to stress than neurons during tissue dissociation (Kurmangaliyev et al., 2020; Marsh et al., 2022; Özel et al., 2021; Simon and Konstantinides, 2021). Indeed, compared with neuronal clusters, glial clusters within the Özel and Kurmangaliyev optic lobe datasets were reported to be enriched for cells with low total gene counts per cell, enriched in mitochondrial transcripts, features of low-quality transcriptomes (Kurmangaliyev et al., 2020; Özel et al., 2021; Simon and Konstantinides, 2021). While standard cut-offs (see Materials and Methods) for total and mitochondrial gene counts were used to filter out low-quality transcriptomes, we speculated that some of the glial clusters which mapped to the same cell-type (hereafter referred to as ‘cluster multiplets’) likely represented the same cell-type split by cell state (*e.g.* based on transcriptome quality or cellular stress). Therefore, we examined the total number of genes, the total number of reads and the proportion of mitochondrial genes relative to the total number of genes for each cluster within a multiplet (Figure 6–supplementary S2A-C). Although ensheathing glial clusters (#3, #5 and #7) and cortex glial clusters (#11 and #13) appeared to separate by transcriptome quality, we found no clear indication that general astrocyte clusters (#0 and #2) differed in this way (Figure 6– supplementary S2A-C). Therefore, we performed Gene Ontology (GO) enrichment analysis on the genes differentially expressed within the general astrocyte clusters with a 1.2-fold cut-off.

Interestingly, this revealed that a wide range of GO terms (from signal transduction and morphogenesis to growth and taxis) were enriched for cluster #0, whereas GO terms associated with mitochondrial regulation, autophagy and metabolic processes were enriched for cluster #2 (Figure 6– supplementary S2D). These data suggested that cells of cluster #2 (astrocyte_2) are unlikely to be a distinct cell type from those of cluster #0 (astrocyte_1), but instead may be cells in distinct metabolic states. Altogether these data suggest that general astrocyte, ensheathing and cortex clusters may be segregating based on transcriptome quality and/or cellular state, possibly driven by tissue dissociation. Furthermore, our data indicate that clusters #0 (astrocyte_1), #3 (ensheathing_1) and #11 (cortex_1) likely correspond to a more homeostatic state of general astrocyte, ensheathing and cortex glia, respectively. We hypothesize that the generation of cluster multiplets occurred in the adult optic lobes due to the requirement to dissect these tissues prior to dissociation, resulting in greater tissue stress compared to our whole embryo dissociations.

Since our data indicated that cluster #0 (astrocyte_1) most likely corresponded to homeostatic astrocytes of the medulla, lobula, and lobula plate, we subclustered it to further probe for any potential heterogeneity (see Materials and Methods and Figure 7A). Although we obtained three subclusters in this way, they differed only subtly from each other. We could identify only one marker gene (*Actin 79B; Act79B*) that was differentially expressed by at least 4-fold among the three subclusters (Figure 5– figure supplement 2 and Figure 7B). Specifically, subclusters #0 and #2 showed enriched expression of *Act79B* relative to subcluster #1 (Figure 7B). Although 96% of the cells in cluster #0 (astrocyte_1; homeostatic general astrocytes) expressed *Act79B* according to the scRNA-seq data, *in vivo* it was expressed very sparsely, in only 10.9% of medulla, lobula and lobula plate astrocytes (Figure 7C,D). These cells were found more frequently in the proximal medulla, the lobula, and lobula plate (Figure 7C,D). *Act79B* encodes what is thought to be a muscle-specific isoform of Actin (Dohn and Cripps, 2018; Ohshima et al., 1997) and therefore is unlikely to be a cell-fate determinant; we speculate instead that *Act79B* may be expressed by cells in a particular state of cytoskeletal remodelling (*e.g.* during process extension or growth). The apparent mismatch between the proportion of *Act79B-*expressing general astrocytes in scRNA-seq and *in vivo* may suggest that tissue dissociation pushes cells towards a state where *Act79B* is upregulated, but that this state is relatively rare under homeostatic conditions (*i.e. Act79B* may label a transient cell-state under homeostatic conditions) (Figure 7C,D). In sum, the unique and stereotyped morphologies of astrocytes of the medulla, lobula, and lobula plate cannot be assigned to unique transcriptional signatures within the present depth of sequencing limits and therefore are unlikely to represent distinct subclasses of astrocytes. Furthermore, any hidden heterogeneity that we have been able to uncover within general astrocytes appears to correspond to a cell-state rather than cell-type or identity.

**Figure 7.**
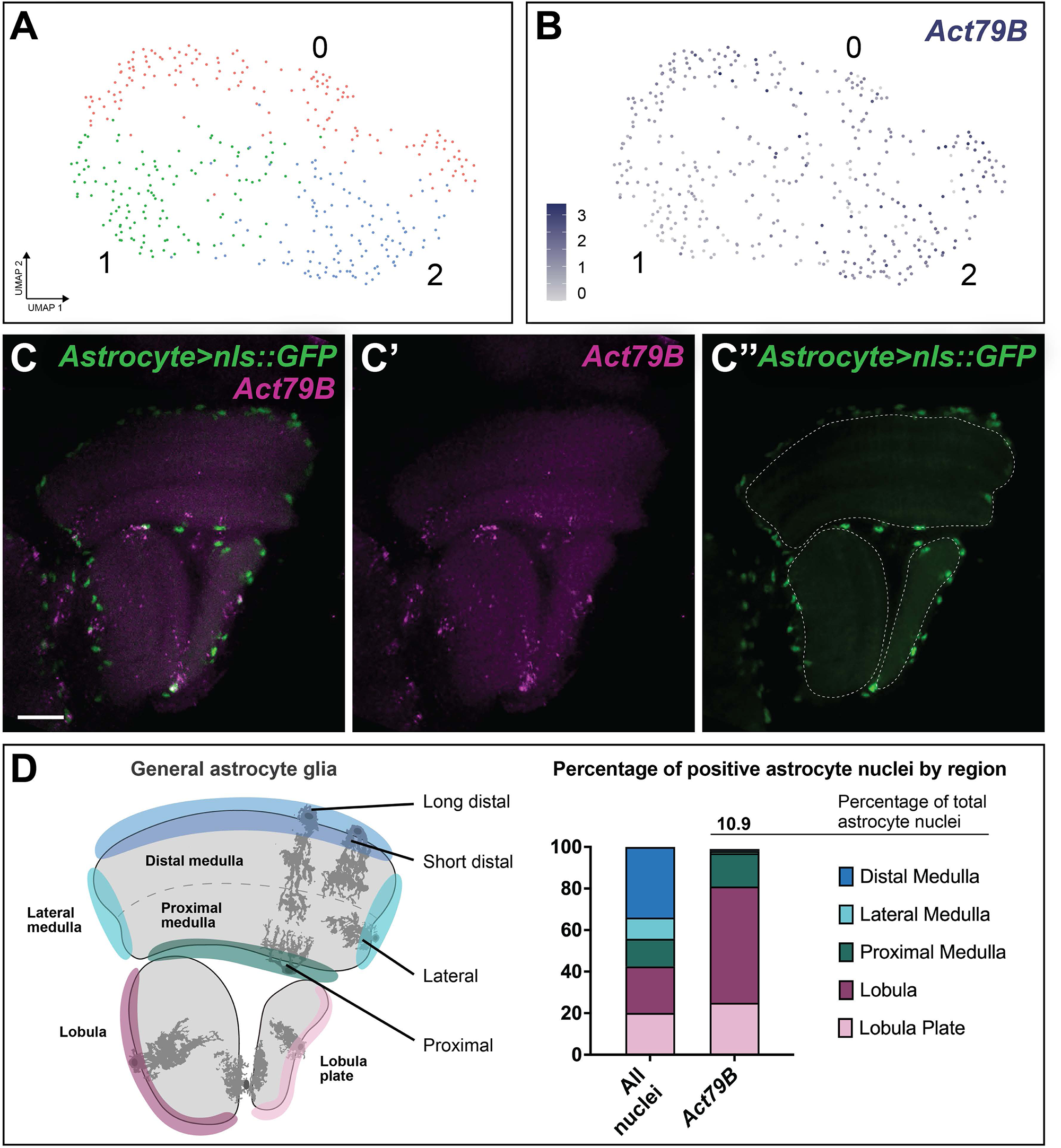
Analysis of clusters obtained from subclustering cluster #0 (Astrocyte_1; homeostatic astrocytes) **(A)** UMAP of the subclusters obtained from the subclustering of Astrocyte_1 (cluster #0). **(B)** *Act79B* expression levels plotted on the same UMAP. **(C)** Maximum projection of *astrocyte(R86E01)>nls::GFP* adult optic lobe where *Act79B* expression (magenta) was detected by *in situ* HCR in a sparse subset of GFP (green) positive astrocyte nuclei of the medulla, lobula, and lobula plate. Dashed lines outline the neuropils and scale bar is 20 µm. **(D)** Quantification of *Act79B* positive astrocyte nuclei (*i.e.* GFP positive nuclei) from (C) Approximately 11% of astrocyte nuclei were positive for *Act79B in vivo*. The percentage of positive nuclei within each indicated region was quantified, showing *Act79B was* preferentially expressed in astrocytes of the lateral and proximal medulla, lobula, and lobula plate (Chi-squared test, *p*<.0001).

### Immune response-related genes are enriched in lamina glia

Lamina glia are the main source of transcriptional diversity in the optic lobes, with more unique cell types associated with the lamina than all other neuropils combined (Figure 8A). Indeed, in addition to its own unique counterparts for the main glial classes, the lamina contains multiple perineurial (fenestrated and chalice), subperineurial (pseudocartridge and carpet), and cortex (distal and proximal satellite) glial subtypes. To investigate lamina glia specialization further, we performed GO enrichment analysis for biological processes on pooled lamina glia and pooled general optic lobe glia (summarised in Figure 8B and Materials and Methods). These comparisons revealed that compared to general glia, lamina glia are enriched for GO terms associated with immune-responses and cell junctions (Figure 8B). Indeed, among the markers for lamina glia we validated *in vivo* in intact optic lobes were *Tsf1, GILT1* and *JhI-21*, which are known to be involved in the antibacterial and/or antifungal immune response (Figure 5 – figure supplement 5D, M,N)(Jin et al., 2008; Kongton et al., 2014; Latsenko et al., 2020; Weber et al., 2022). By contrast, general glia were enriched for terms associated with metabolite transport, intercellular signalling, and migration relative to lamina glia (Figure 8B). Thus, lamina glia appear to be primed to perform immune-related functions relative to their general glial counterparts.

**Figure 8.**
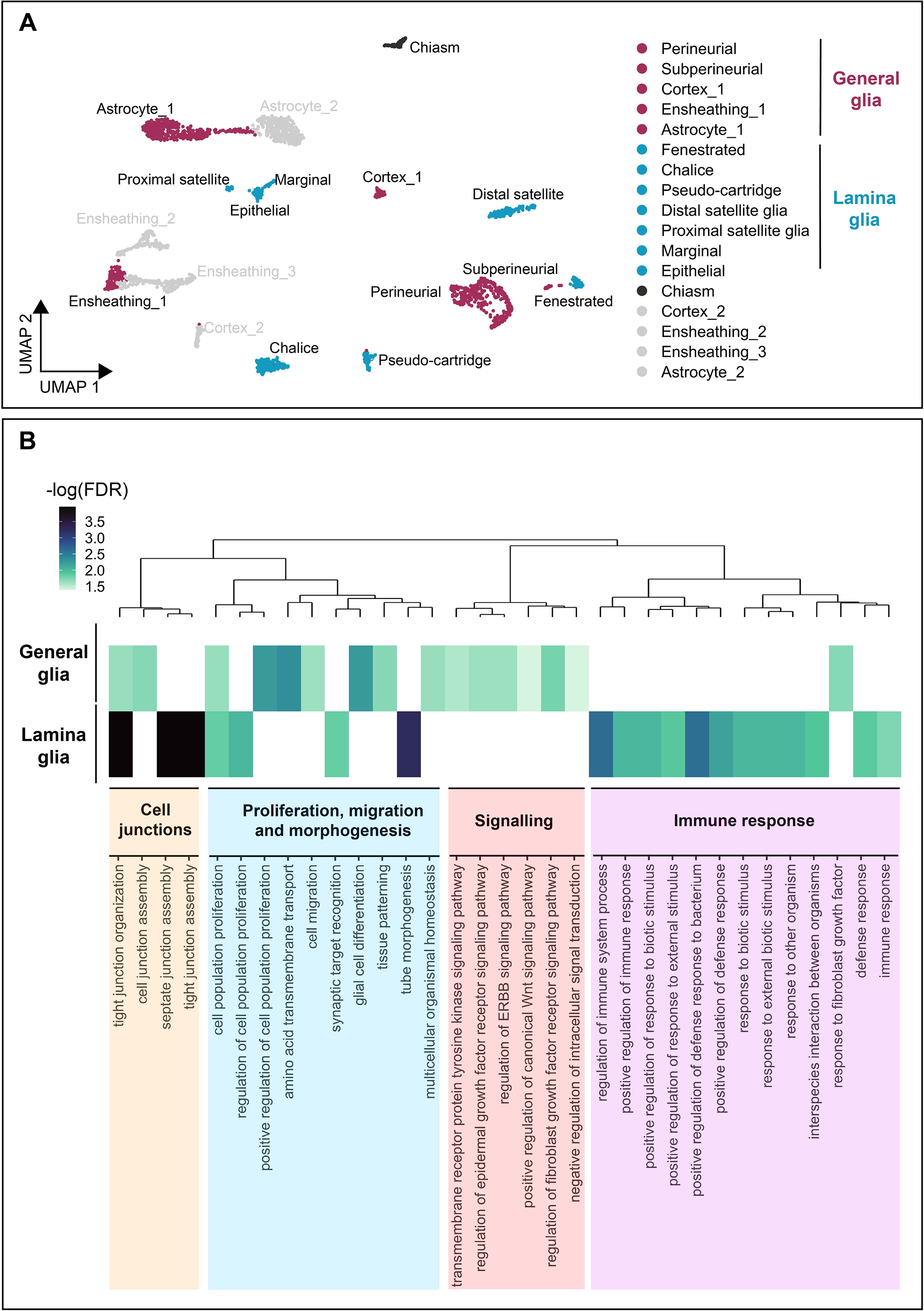
Lamina glia are specialized to perform immune-related functions. **(A)** UMAP of the adult optic lobe glia colour-coded for lamina (maroon) and ‘general’ (medulla, lobula, and lobula plate; blue) glial clusters that were included in the GO analysis in (B). Note that the chiasm glia cluster (dark grey) and non-homeostatic clusters belonging to multiplets (light grey) were excluded from the analysis in (B). **(B)** GO enrichment analysis for biological processes of the lamina glia compared to other (general) optic lobe glia shown as a heatmap. We used a dendrogram to cluster GO terms into groups that were defined by a super-term. The scale indicates the negative log of the false discovery rate (FDR).

### Motor and sensory-associated glia are transcriptionally similar

Having shown that, apart from the lamina, optic lobe glia within each class are transcriptionally similar irrespective of morphology (*e.g.* all astrocyte morphotypes are transcriptionally similar, with the exception of lamina astrocytes), we sought to evaluate how different glia are between the young adult optic lobe and embryo, where developmental stage, circuits and afferent inputs are vastly different. To address this question, we integrated the embryonic dataset with the optic lobe dataset, representing motor and sensory-associated glia, respectively. Note that our embryonic dataset, though generated from whole brains, did not contain any optic lobe glia as they are not born until late larval and pupal development (Chotard and Salecker, 2007). Strikingly, we found that the embryonic clusters and their corresponding adult general glia clusters (*i.e.,* not the lamina clusters) converged on the UMAP visualization (Figure 9A-C). Indeed, several of our validated marker genes were found in both embryonic glia and their counterparts in the adult optic lobe (summarised in Figure 9D). Interestingly, there was no overlap with lamina glia clusters, except for the embryonic channel perineurial glia and the chalice glia, suggesting that they may be functionally analogous (Figure 9B,C).

**Figure 9.**
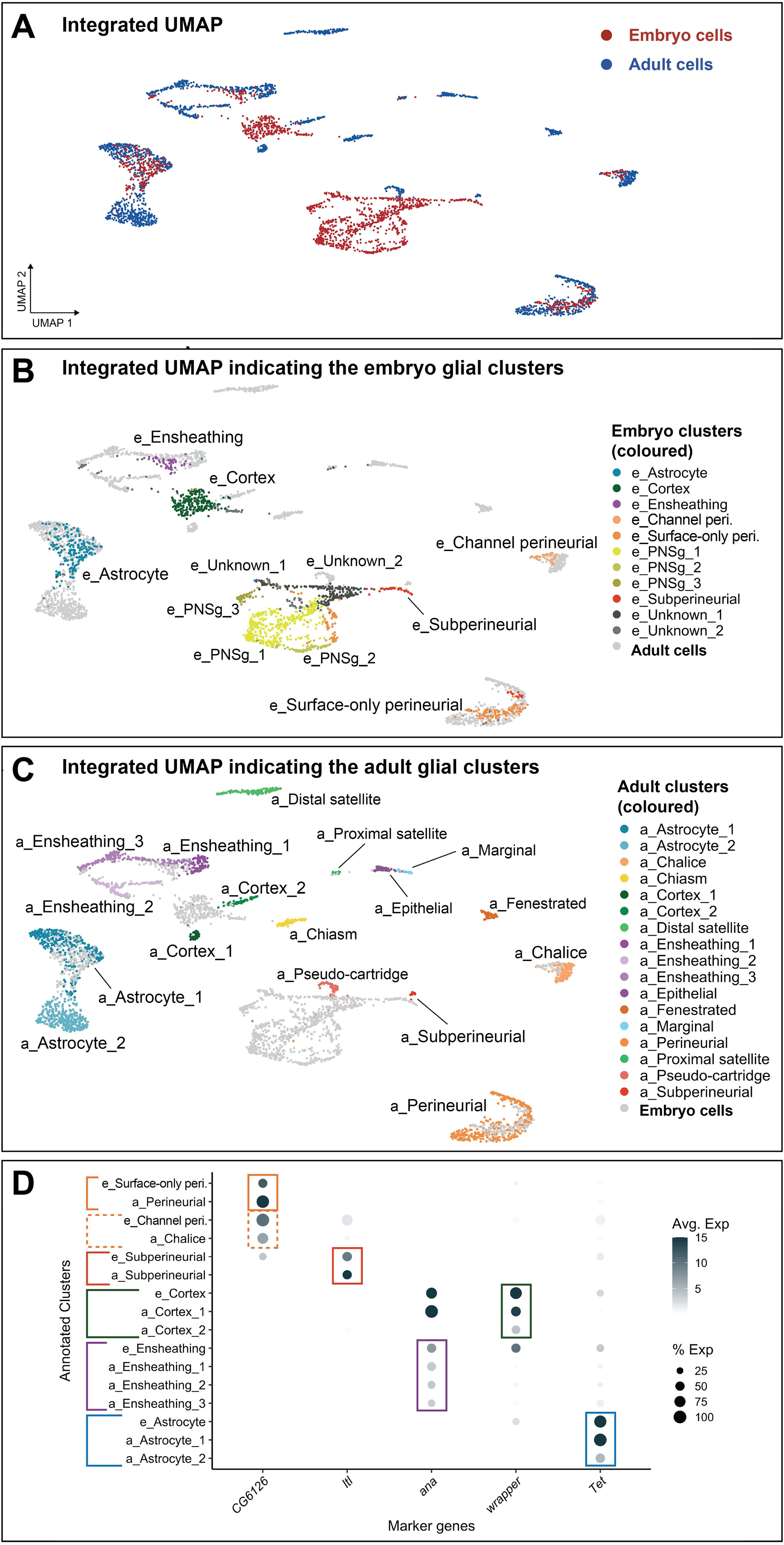
Integration of the embryonic stage 17 CNS and young adult optic lobe glial clusters. **(A)** UMAP of the integrated dataset, highlighting the embryonic (red) and young adult optic lobe (blue) glial clusters. **(B-C)** UMAP of the integrated dataset, highlighting embryonic (B, ‘e_’) or the young adult optic lobe (C, ‘a_’) glial clusters in colours by glial class. Grey marks cells from the young adult optic lobe clusters in (B) and from the embryonic clusters in (C). **(D)** Expression plot of genes with expression in the same glial class of the embryo and adult. The size of the dot represents the percentage of cells with expression in each cluster, while the colour of the dot represents the level of average expression in the cluster. The five genes indicated were validated *in vivo* for both time points by independent strategies: *CG6126* for perineurial (embryo only-surface perineurial, Figure 4– figure supplement 1B,I,J; adult perineurial, Figure 5– figure supplement 4F) and chalice (Figure 5– figure supplement 4F) and channel perineurial (Figure 4E, Figure 4– figure supplement 1I,J), *ltl* for subperineurial (embryo, Figure 4– figure supplement 1E,F,I,K; adult, Figure 5– figure supplement 4C-E), *ana* for ensheathing (embryo, Figure 4– figure supplement 3A; adult, Figure 5– figure supplement 4I), *wrapper* for cortex (embryo, Figure 4F; adult, Figure 5– figure supplement 3B) and *Tet* for astrocyte (embryo, Figure 4– figure supplement 2B,D,E; adult, Figure 5– figure supplement 3A).

Altogether, our analysis uncovered a surprising transcriptional similarity between glia from different circuit types, brain regions, and time points in development. This suggests that, within each general class (astrocyte, cortex, ensheathing, etc.), glia likely fulfil similar functions from circuit to circuit and across neuropils.

### Discussion

We generated, validated, and annotated a single cell transcriptional atlas of *Drosophila* glia spanning embryo to adult. In validating our dataset, we identified many new marker genes for known glial classes and subclasses, and identified a new glial subclass, the channel perineurial glia. We hope that this atlas will serve as a community resource and facilitate functional studies to unveil how distinct glial populations coordinate the development and function of neural circuits.

#### Glial morphological diversity and detectable transcriptional diversity were not correlated

While we must be cautious in interpreting our results given the depth of sequencing constraints associated with scRNA-seq approaches, our data, nevertheless, point to a mismatch between glial morphological and transcriptional diversity. Specifically, glia exhibited more diversity at the morphological level than was detectable at the transcriptional level.

Although we detected striking morphological diversity in embryonic VNC subperineurial glia, this diversity could not be easily divided into discrete categories, as these glia clearly exhibited a continuum of morphologies from one extreme to the other (surface-only to channel-only) (Figure 1H-L). Similarly, embryonic ensheathing glia also exhibited a range of morphologies from neuropil-only association to both neuropil and axon tract association to varying degrees (Figure 1N,O). Nonetheless we detected only one transcriptional cluster corresponding to all subperineurial glia and one transcriptional cluster for all ensheathing glia. Even when glial morphologies could be divided into discreet categories such as for the newly-hatched larval VNC and the adult optic lobe, astrocytes (two and eight discreet morphological categories, respectively; Figure 1A,P-S, Figure 2A,Q-W and Figure 2 – figure supplement 2), detected transcriptional diversity did not match with morphological diversity. While astrocytes of the lamina were transcriptionally distinct, we identified only one transcriptional cluster of true, homeostatic astrocytes that corresponded to all remaining astrocyte morphologies at both developmental stages and brain regions. Indeed, when integrated, embryonic and adult glia co-clustered by class (except for lamina glia), despite the difference in developmental stage and origin, highlighting a striking convergence in their transcriptional profiles. This result was unexpected as the embryonic dataset was enriched for glia within a sensorimotor circuit (Grueber et al., 2007; Landgraf et al., 2003; Mark et al., 2021; Merritt and Whitington, 1995; Zlatic et al., 2009), and the adult dataset was specific to the visual system. Thus, within the present limits of detection, our data suggest that glial classes are not transcriptionally specialised to the circuit types they associate with.

While at first glance it may appear that some glial morphologies of the adult optic lobes were associated with unique transcriptional signatures (*e.g.* lamina astrocytes), the glial morphological categories with unique transcriptional signatures were all associated with the lamina. This suggests that the observed transcriptional specialisation is due to lamina association rather than specialised morphology. Interestingly, the glial transcriptional diversity we defined matches remarkably well with glial diversity described previously by glial-specific drivers from the Janelia GAL4 collection (Kremer et al., 2017; Pfeiffer et al., 2008). These driver lines also did not distinguish between cortex and ensheathing morphotypes of medulla, lobula and lobula plate neuropils (Kremer et al., 2017). It will be interesting to explore if lineage relationships alone account for the transcriptional diversity detected in glia. Glial lineage relationships are characterised with detailed cellular resolution for the embryonic VNC where cell numbers are low, but are still incomplete for the optic lobes (Beckervordersandforth et al., 2008; Edwards et al., 2012).

We hypothesize that glial morphology may be a plastic cellular feature that is set by the local environment and thus may represent a cellular state rather than an underlying transcriptionally defined identity. In the future, one could leverage genetic manipulations that alter neuropil structure to assess how equivalent glia might alter their morphology and transcriptional profiles in distinct environments. For example, one could assess glial morphologies and transcriptional profiles while (i) altering neuronal diversity and composition by manipulating the temporal series in medulla neuroblasts (Erclik et al., 2017; Konstantinides et al., 2021; Li et al., 2013; Zhu et al., 2022), or by blocking lamina neuron differentiation (Fernandes et al., 2017; Prasad et al., 2021), or (ii) by modifying neuropil stratifications with genetic perturbations that disrupt neuronal targeting (Zhang et al., 2023).

#### Specialisation of the Drosophila lamina glia

Transcriptomic studies in mammals have uncovered regional differences in astrocyte transcriptional profiles (*e.g.* between and within the cortex and hippocampus) (Batiuk et al., 2020; Bayraktar et al., 2020). In our optic lobe data, we could only detect within-class transcriptional specialisation for glia associated with the lamina, whereas we could not detect any within-class transcriptional differences between (or within) the other optic lobe regions. This is particularly surprising since others have speculated that circuit complexity (stemming from neuronal diversity) may drive glial specialization (Durkee and Araque, 2019; Khakh and Sofroniew, 2015), however, the lamina is the simplest of all neuropils, while the medulla is the most complex (Fischbach and Dittrich, 1989). Thus, at least for the optic lobes, region-specific neuronal diversity does not appear to be a predictor of glial transcriptional diversity, within the present limits of detection.

Instead, our data revealed an upregulation of GO terms related to immune functions in lamina glia when compared to other optic lobe glia (Figure 8B). The lamina is the first neuropil to receive input from photoreceptor sensory neurons (Apitz and Salecker, 2014; Fischbach and Dittrich, 1989) and, as such, its surface glia are densely perforated to allow photoreceptor axons to enter the lamina (Edwards and Meinertzhagen, 2010). Thus, the lamina may be subject to more pathogenic insults compared to other optic lobe neuropils. This finding opens up the intriguing hypothesis that glia that interface with the environment may be specialised to manage environmental insults (*e.g.* heat, cold, lesions). Recently, a specialised population of cutaneous Schwann cells was identified which sense noxious stimuli and initiate pain sensation (Parfejevs et al., 2018). Furthermore, Schwann cells can migrate away from peripheral nerves to aid in cutaneous wound healing following injury, which is accompanied by an upregulation of genes related to the innate immune response (Abdo et al., 2019). Defining the extent to which other glial cell populations at environmental boundaries adopt immune-related roles will broaden our understanding of the functional properties of glia not only for nervous system function, but organismal health.

#### Limitations of this study

It is possible that all within-class glial morphological categories are transcriptionally distinct, but that the genes that confer this diversity are lowly expressed and therefore were missed due to depth of sequencing limitations. It is also possible that functional specialization may occur locally in micro-domains of the same cell. For example, astrocytes are intimately associated with neuronal synapses. At the synapse, astrocytes regulate synapse strength and turnover of local neurotransmitters. As a single astrocyte supports many distinct types of synapses simultaneously, each requiring different molecular machinery (Durkee and Araque, 2019), astrocytes must express a wide variety of channels and receptors (resulting in transcriptional homogeneity). Functional diversity could therefore be conferred by local synaptic activity. In other words, a single astrocyte may have functionally distinct microdomains dictated by the synapses it supports. Indeed, recent studies show that synapse activity regulates positioning of neurotransmitter receptors in astrocyte processes (Ciappelloni et al., 2017; Murphy-Royal et al., 2015; Muthukumar et al., 2014). Functional studies, including *in vivo* Calcium, Glutamate, or GABA imaging will help to resolve glial functional specialization both within and across cells.

A major anatomical difference between *Drosophila* and mammals is the size of glial domains relative to the size of the brain. Although only 10-15% of the Drosophila brain is made up of glia, astrocytes tile the entire CNS. For example, six individual astrocytes are sufficient to tile a single hemisegment of the larval VNC (Peco et al., 2016). In contrast, thousands of astrocytes are required to tile an analogous section of the mouse spinal cord (Sun et al., 2017). As mammalian astrocytes proportionally tile a much smaller region of the nervous system, it is conceivable that a single mammalian astrocyte can interact with one circuit type. In contrast, Drosophila astrocytes frequently span multiple circuits, which may require more functional flexibility. Thus, it is possible that further exploration of astrocyte diversity in vertebrates will identify unique transcriptional programs that specify morphologically distinct astrocytes with regionalized identities and functions. In sum, our work highlights the necessity for a systematic multimodal approach to characterising glial diversity in other systems (Khakh and Deneen, 2019; Westergard and Rothstein, 2020).

## Materials and Methods

### Drosophila stocks and maintenance

*Drosophila melanogaster* strains and crosses were reared on standard cornmeal medium and raised at 25^°^C unless stated otherwise. We used the following strains in this study (See Supplementary file 4 for more details):

*hs-FLPG5;; 10×UAS(FRT.stop)myr::smGdP*-*HA, 10×UAS(FRT.stop)myr::smGdP-V5, 10×UAS(FRT.stop)myr::smGdP-Flag* (hsMCFO; BDSC 64085), *pBPhsFlp2::PEST in attP3;; HA_V5_FLAG_OLLAS* (BDSC: 64086), *CG6126-Gal4* (BDSC 67505), *CG5080-Gal4* (BDSC 83275), *CG10702/CG17343-Gal4* (BDSC 76234), *Ntan1^Mz97^-Gal4* (BDSC 9488), *PRL-1-Gal4* (BDSC 65566), *R54H02-Gal4* (BDSC 45784), *Eaat2-Gal4* (BDSC 78932), *alrm-Gal4* (BDSC 67031 and 67032), *Rdl-Gal4* (BDSC 65421 and 66509), *moody-Gal4* (BDSC 90883), *R25H07-Gal4* (BDSC 49145), *Tet-Gal4* (BDSC 19427), *pum-Gal4* (BDSC 63368), *w[*] TI{RFP[3xP3.cUa]=TI}Tre1[attP]* (BDSC 84582), *ana-Gal4* (BDSC 86394), *CG9657-Gal4* (BDSC 78971), *CG9449-Gal4* (BDSC 91274), *pippin-Gal4* (BDSC 86401), *ltl-Gal4* (BDSC 76144), *PRL-Gal4* (BDSC 81151), *hs-flp;lexAop-myr::tdTomato; repo-lexA*, *R85G01-Gal4* (BDSC: 40436), *R54C07-Gal4* (BDSC: 50472), *R10C12-Gal4* (BDSC: 47841), *R47G01-Gal4* (BDSC: 45768), *R50A12-Gal4* (BDSC: 47618), *R46H12-Gal4* (BDSC: 50285), *R53H12-Gal4* (BDSC: 50456), *R56F03-Gal4* (BDSC: 39157), *R35E04-Gal4* (BDSC: 48127), *R86E01-Gal4* (BDSC: 45914), *55B03-Gal4* (BDSC: 39101), *Repo-Gal4* (BDSC: 7415), Mi{MIC}CG43795-GFP[MI03737] (BDSC: 41395), 10XUAS-mCD8::GFP (BDSC 32184 and 32186), 10XUAS-IVS-myr::GFP (BDSC 32197) and UAS-nls::GFP (BDSC 4775 and 4776).

### Embryo collections

For timed collections of embryonic and larval stages, crosses were reared in collection bottles fitted with 3% agarose apple caps coated with yeast paste. Embryos were then collected for 1.5-hour (h) windows and reared at 25^°^C until the desired developmental stage.

### Multi Color Flip-Out clonal analyses (See Supplementary file 4 for list of genotypes)

#### Embryonic/larval ventral nerve cord

To generate multi-color flip-out (MCFO) clones, we crossed virgin females of the hsMCFO line to males of Gal4 lines that were known to be expressed (Coutinho-Budd and Freeman, 2013; Doherty et al., 2009; Peco et al., 2016; Schwabe et al., 2005) or found to be enriched in a given transcriptional cluster (see Supplementary file 4 for full genotype list).

Embryos were collected for 1.5 h, aged for 6 hours (embryonic stage 11/12, prior to gliogenesis (Hartenstein et al., 1998)), and then heatshocked at 37^°^C for 15 minutes to induce FLP expression. Embryos were then transferred to 4^°^C for 5 minutes to halt FLP expression and transferred to 25^°^C until hatching. At 25^°^C, hatching occurs at 21 h after egg laying (Crisp et al., 2008).

#### Adult optic lobe

For MCFO experiments of adult brains we utilised Janelia Gal4 driver lines with specific glial expression, as previously described (Kremer et al., 2017) (Figure 2– figure supplement 1). We crossed males of the glial-Gal4 lines with virgin females of a 3-tag (HA, V5 and FLAG) MCFO line, except the cortex and marginal Gal4s which we crossed with a 4-tag (HA, V5, FLAG and OLLAS) MCFO (Supplementary file 4). We raised progeny at 18^°^C, and induced FLP expression in adult flies (0-5 days old) by heat-shocking at 37^°^C. Length of heat shocks varied depending on the Gal4 driver (Supplementary file 4). After heat-shocking, flies were placed back at 18^°^C for at least 1 night, before being dissected and stained as detailed in our immunohistochemistry protocol.

### Immunohistochemistry

#### Larval VNC

For VNC studies, larvae were dissected at 0 h after larval hatching. We dissected larval brains in sterile-filtered, ice-cold 1× PBS. Brains were then mounted on 12 mm #1 thickness poly-d-lysine coated round coverslips (Neuvitro Corporation, GG-12-pdl) and fixed in fresh 4% paraformaldehyde (Electron Microscopy Sciences, 15710) in 1× PBS with 0.3% Triton detergent (0.3% PBST). We then washed the coverslips in 0.3% PBST to remove fixative and blocked overnight at 4°C in 0.3% PBST supplemented with 1% BSA (Fisher, BP1600-100), 1% normal donkey serum and 1% normal goat serum (Jackson ImmunoResearch Laboratories 017-000-121 and 005-000-121). Brains were then incubated in primary antibody overnight at 4 °C, washed overnight at 4 °C with 0.3% PBST, and then incubated in secondary antibodies overnight at 4 °C. We then removed the secondary antibodies, transferred the coverslips to 0.3% PBST overnight, and mounted in DPX. To mount in DPX, brains were dehydrated with an ethanol series: 30%, 50%, 70% and 90%, each for 5 min, then twice in 100% ethanol for 10 min each (Decon Labs, 2716GEA). Samples were then transferred to glass-bottomed depression slides with xylenes (Fisher Chemical, X5-1) for 2 × 10 min. Finally, samples were mounted onto slides containing DPX mountant (Millipore Sigma, 06552), and cured for 1–2 days before imaging.

#### Adult optic lobe

For immunocytochemistry experiments of the adult optic lobe, we dissected whole brains in 1X phosphate buffered saline (PBS), fixed in 4% paraformaldehyde for 30 minutes, then washed with 0.5% PBTx (1X PBS with 0.5% TritonX). We next incubated the samples with primary antibodies diluted in block (5% normal horse serum), for two nights. Samples were washed with 0.5% PBTx, incubated for a further two nights with secondary antibodies diluted in block, washed again and mounted in SlowFade (Life Technologies).

We used the following primary antibodies: Rabbit anti-Pros (1:1000)(Vaessin et al., 1991), Rat anti-HA (1:100; Millipore Sigma, 11867423001), Chicken anti-V5 (1:1000; Bethyl Laboratories, A190-118A), Mouse anti-Repo (1:50, DSHB 8D12), Mouse anti-Cherry (1:500; Clontech, 632543), Rabbit anti-Gat (1:4,000; a gift from M. Freeman)(Stork et al., 2014), rat Anti-FLAG (1:400, Novus NBP1-06712), rabbit Anti-HA-tag (1:400, Cell Signalling Technology, C29F4), rat anti-Elav (1:20, DSHB 7E8A10), mouse anti-V5-Tag:DyLight-550 mouse (1:400, Bio-Rad), chicken anti-GFP (1:400, EMD Millipore), rabbit anti-GFP (1:400, Thermo Fisher A6455), DyLight 405 conjugated Goat Anti-HRP (1:50, Jackson ImmunoResearch, 123-475-021), Rabbit anti-dsRed (1:500, Takara, 632496) and mouse anti-FasIII (1:20, DSHB 7G10).

We used the following secondary antibodies at 1:400: Alexa Fluor Rhodamine Red-X Donkey-Anti Mouse (Jackson ImmunoResearch, 715-295-151), Alexa Fluor 488 Donkey anti-Rabbit (Jackson ImmunoResearch, 711-545-152), DyLight 405 Donkey-anti Rabbit 405 (Jackson ImmunoResearch, 711-475-152), Alexa Fluor 488 Donkey anti-Chicken (Jackson ImmunoResearch, 703-545-155), Alexa Fluor Rhodamine Red-X Donkey-Anti Rat (Jackson ImmunoResearch, 712-295-153), Alexa Fluor 647 Donkey anti-Mouse (Jackson ImmunoResearch, 715-605-151) and Alexa Fluor 647 donkey anti-rat (Jackson Immunolabs, 712-605-153).

### Embryo dissociation for single cell RNA sequencing (scRNA-seq)

We prepared cell dissociates from embryos at 17-18.5 h after egg laying (stage 17). We washed embryos in deionized water before surface sterilizing them in 30% bleach for 2 minutes. We then homogenized them in Chan-Gehring (C + G) medium by six to eight strokes of a loose-fitting dounce. We filtered the cell suspension through a 40 μm Nitex mesh, and pelleted cells in a clinical centrifuge at 4°C (setting 5, IEC). We washed the cell pellet twice by pouring off the supernatant and gently resuspending the pellet in fresh C + G, pelleting between each rinse as above. We determined the cell-survival proportion for each dissociate using the BioRad TC-20 trypan-blue assay. Samples that met a threshold of 80% viability were submitted for sequencing at a concentration of 1000 cells per microliter.

### Single cell RNA sequencing of stage 17 embryos

The University of Oregon Genomics and Cell Characterization core facility (https://gc3f.uoregon.edu/) prepared embryonic cell samples for scRNA-seq. We ran dissociated cells on a 10X Chromium platform using 10X NextGem v3.1 chemistry targeting 10,000 cells per sample. Following cDNA library preparation, we amplified the library with 15 cycles of PCR before sequencing on two separate Illumina Hi-seq lanes, providing two technical replicates. Following examination for batch effects between technical replicates (Supplementary File 3), we merged the datasets using the CellRanger Aggregate function prior to quality control and downstream analysis. Reads were aligned to the Drosophila genome (BDGP6.22) and protein coding reads were counted.

### Initial quality control and cell clustering of glia from embryonic scRNA-seq

We analysed the resulting sequencing data with the 10X CellRanger pipeline, version 3.1.0 (Zheng et al., 2017), R version 3.6.3 and Seurat (Hao et al., 2021) version 3.1.2 using standard quality control, normalization, and analysis steps. We filtered cells by the percentage of mitochondrial genes expressed (relative to the total number of genes expressed), indicating a high stress state. Only cells expressing <10% mitochondrial reads were retained for analysis. Additionally, cells containing reads for <50 and >3000 unique genes were filtered out of downstream analysis. For each gene, expression levels were normalized by total expression, multiplied by a scale factor (10,000) and log-transformed, and the top 3000 variable genes were identified for downstream Principal Component

Analysis (PCA). Clustering was performed using 50 PCs [*FindClusters* resolution 1.0], resulting in a Seurat object containing 52,881 cells. To isolate neuronal and *repo+* glial cell clusters from all somatic tissues, we visualized the expression of *elav*, *repo*, *Gat*, and *alrm* in UMAP space. We then selected embryonic clusters 18, 2, 0, 4, 24, 26, 22, 16, 13, 14, 11 and 27 to be subset for further analysis. The resulting subset of cells were re-clustered using the previous parameters, resulting in the stage 17 embryonic neurons and *repo+* glia dataset (19,600 cells). To isolate embryonic *repo+* glia from embryonic neurons, we again visualized the expression of *elav*, *repo*, *Gat*, and *alrm* in UMAP space. From this object, clusters 23, 26, 34, 38, 17, 41, 27, 32 and 19 were identified as glial/non-neuronal. All the scripts described above are available at https://github.com/AustinSeroka/2022_stage17_glia. The resulting dataset (3221 cells) was exported (.Rds) for subsequent analyses.

Midline glia are known to express *sim* and *wrapper,* but do not express *elav* or *repo* (Jacobs, 2000). Therefore, to identify midline glia from the full dataset we visualized *sim, wrapper, elav* and *repo* expression in UMAP space. We identified a small group of cells meeting these expression criteria, which were clustered together and belonged to a larger cluster (cluster 19). We selected these cells as putative midline glia and defined them as a new cluster within the whole embryonic dataset. We pooled midline glia together with the stage 17 embryonic neurons and *repo+* glia dataset described above and performed hierarchical clustering analysis [*hclust*] on this combined dataset, using average cluster gene expression to produce an euclidean dissimilarity matrix [*dist*]. We used [*FindAllMarkers*] (default parameters, except only.pos = TRUE) to obtain the most highly expressed genes for putative midline glia relative to all other embryonic cells (See Source data file 1).

### Preliminary integration of published optic lobe glial single cell RNA sequencing datasets

Two independent studies published scRNA-seq datasets of the optic lobes throughout pupal development with a focus on neuronal development and therefore did not analyse glial clusters (Kurmangaliyev et al., 2020; Özel et al., 2021). We took advantage of the glial cells within these datasets here. Özel et al. (2021) performed scRNA-seq on optic lobes at specific stages in triplicate, profiling a total of 275,000 cells, whereas Kurmangaliyev et al. (2020) used a novel strategy to perform multiplexed scRNA-seq of many developing brain samples in parallel and profiled 51,000 cells in total. The two datasets (hereafter referred to as the Özel and Kurmangaliyev datasets, respectively) thus varied dramatically in the number of cells they each profiled. Here we chose to focus specifically on optic lobe glia from developmental stages that most closely match the young adult, *i.e.* young adult dataset from Özel et al. (2021) and 96h APF dataset from Kurmangaliyev et al. (2020) We used R (version 4.1.0) and Seurat (version 4.0.3) (Hao et al., 2021) to analyse the optic lobe scRNA-seq datasets from Özel et al. (2021) (NCBI GEO accession GSE142787) and Kurmangaliyev et al. (2020) (NCBI GEO accession GSE156455). [] enclose the specific Seurat functions used.

We converted the Kurmangaliyev 10X Genomics data to an RDS file [*CreateSeuratObject*] and subsetted [*subset*] the cells belonging to the 96h APF timepoint (within metadata, under ‘time’). Glial clusters were subsetted from both the 96h APF-Kurmangaliyev dataset (within metadata, under ‘class’) and adult-Özel dataset based on the original annotation; *i.e. repo* expression. Since the two datasets represent glia from the same structure and from very close developmental stages, we used Seurat’s Integration pipeline to find corresponding cells and batch correct the two datasets (Figure 3). We normalized each glial dataset [*NormalizeData*] and selected the 2000 most variable features [*FindVariableFeatures*]. We defined Anchors [*FindIntegrationAnchors*, dims = 1:30] and integrated the datasets [*IntegrateData*, dims = 1:30]. We then scaled the integrated dataset [*ScaleData*] and ran PCA [*RunPCA*, npcs = 30]. We checked elbow plots [*ElbowPlot*] to determine the number of dimensions to use in [*FindNeighbors*, dims = 1:16], followed by [*FindClusters,* resolution = 0.5]. We obtained 20 clusters and used a Uniform Manifold Approximation and Projection (UMAP) dimensionality reduction [*RunUMAP*, reduction = pca, dims = 1:20] to plot the cell clusters and visualise their distribution as a 2-dimensional representation [*DimPlot*] (Figure 3 and Figure 3-figure supplement 2C,D).

### Further clean-up of young adult optic lobe and embryonic glial datasets

#### Neuronal and hemocyte clean-up

While analysing the adult optic lobe dataset we noticed that most glial clusters contained cells that formed streams that extended towards the center of the UMAP and that cells in these streams expressed high levels of *Rdl*, *Frq1*, *Nckx30C, elav* and *fne* whereas cells belonging to the main body of clusters did not express these genes (Figure 3– figure supplement 2A). *elav* and *fne* are well-documented pan-neuronal marker genes. Indeed, we found that *Rdl, Frq1* and *Nckx30C* genes were also expressed pan-neuronally (Figure 3– figure supplement 2B). Thus, we hypothesized that they likely represented contamination from neurons due to the close association of glia and neurons. We also observed *Rdl, Frq1*, *Nckx30C, elav* and *fne* co-expressing cells in the embryonic glial dataset (mainly from cluster 4) and pan-neuronally (Figure 3– figure supplement 2C,D). We validated that *Frq1* and *Nckx30C* are expressed neurons and not glia by HCR in the adult optic lobes (Figure 3– figure supplement 3A-C), and similarly through MCFO clonal analyses in the embryo found that *Rdl* was exclusively expressed in neurons (Figure 3– figure supplement 3D). We then examined the distribution of cells expressing different levels of *Rdl*, *Frq1* and *Nckx30C* in individual glial clusters and all neuronal clusters averaged [*RidgePlot*] (Figure 3– figure supplement 3E,F), and used these distributions to choose a cut-off to eliminate *Rdl, Frq1* and *Nckx30C* expressing cells from the glial clusters [*subset, Rdl/ Frq1/ Nckx30C* ≤ 1] (5101 cells in total; Figure 3– figure supplement 3G,H). To remove contaminated cells in the embryo, we used the same thresholds (Figure 3– figure supplement 3J-N) as the adult dataset (346 cells in total).

We also eliminated putative hemocytes from both the young adult optic lobe (3 cells) and embryonic (10 cells) glial datasets by removing cells that expressed *Hml*, a hemocyte-specific marker [*subset*, *Hml* ≤ 0].

### Re-clustering the embryonic glial dataset

Following the clean-up described above for the embryonic glial dataset, we normalized the data [*NormalizeData*], selected the 2000 most variable features [*FindVariableFeatures*], scaled the data [*ScaleData*] and ran PCA [*RunPCA*, npcs = 30]. We examined elbow plots [*ElbowPlot*] to determine the number of dimensions to use in [*FindNeighbors*, dims = 1:16], followed by [*FindClusters*, resolution = 0.5]. We obtained 11 clusters and plotted them as a 2-dimensional representation ([*RunUMAP*, reduction = pca, dims = 1:20]; [*DimPlot*]; Figure 3; See also Figure 3– figure supplement 3G,H). We used [*FindAllMarkers*] (default parameters, except only.pos = TRUE) to obtain the most highly expressed genes for the embryonic glia (See source data file 2).

### Re-integrating the young adult optic lobe dataset and eliminating clusters originating from a single dataset

Following the clean-up described above for the young adult optic lobe glial dataset we obtained a list of the cells remaining using [*WhichCells*] (all identities, by default *ident* argument) and isolated these cells from the original 96h APF – Kurmangaliyev and young adult – Özel datasets (using [*subset*], and lists of cells used as *cells* argument). We then integrated these as described previously; [*FindNeighbors*, dims = 1:19] and [*FindClusters*, resolution = 0.5] (Figure 3– figure supplement 2I). We obtained 19 clusters. Since others have reported that glial clustering is sensitive to batch effects (Kurmangaliyev et al., 2020; Özel et al., 2021; Simon and Konstantinides, 2021), we eliminated clusters to which the Kurmangaliyev dataset contributed fewer than 1% of the total number of cells in the cluster (Figure 3– figure supplement 3I; a total of 850 cells were eliminated) and reintegrated the remaining cells as described above (Figure 3). Following this, the young adult optic lobe glial dataset consisted of 15 clusters (Figure 3).

### Comparing cell-type-specific bulk RNA sequencing to scRNA-seq

Cell-type-specific transcriptomes, obtained by Tandem-affinity purification of intact nuclei sequencing, were published for the marginal glia, epithelial glia and the proximal satellite glia (Davis et al., 2020) (NCBI GEO accession #GSE116969). We simulated the FACS-sorted glial-type-specific transcriptomes as single-cell transcriptomes to enable us to compare these deep bulk RNA sequencing datasets with the shallower scRNA-seq dataset, as described in Konstantinides et al., 2018. Briefly, a random number of reads were assigned to each of 900 simulated cells, from a normal distribution with the same mean and standard deviation as the cells in the final Adult-96hAPF dataset. The probability of expression of each gene was obtained by dividing the number of transcripts per million (TPM) of each gene by the total, in each cell-type, and this probability was used to allocate the number of reads of each simulated cell. The simulated matrix of expression was transformed into a RDS object, normalised [*NormalizeData*] and average expression of each gene was calculated [*AverageExpression*]. We then examined the Pearson correlations between the average expression of the simulated single-cell and the average expression of the scRNA-seq clusters to determine the best match.

### Sub-clustering individual glial clusters

To analyse individual young adult optic lobe glial clusters, we used [*WhichCells*] to obtain the list of cells belonging to a specific cluster and subsetted those cells from the original Kurmangaliyev and Özel datasets [*subset*]. We then integrated these as described above. The arguments *dims, k.score* and *k.filter* in [*FindIntegrationAnchors*], and *dims* and *k.weight* in [*IntegrateData*], were assigned a value of 1 unit lower than the smallest number of cells in either the Kurmangaliyev or Özel subsetted cells. Next, we used [*FindNeighbors*] and [*FindClusters*] (see Figure 5– figure supplement 2 for specific dimensions and resolutions used).

### Adding subclusters to main young adult optic lobe glial dataset

Figure 5– figure supplement 2 summarizes the number of subclusters obtained following sub-clustering of individual glial clusters. To account for over-clustering artefacts, we examined the number of genes that were differentially expressed between subclusters. Only the subclusters of clusters 8 and 9 expressed greater than 20 genes differentially (4-fold change or higher), whereas all other subclusters expressed fewer than 15 genes differentially. Therefore, we proceeded to manually divide clusters 8 and 9 accordingly (Figure 5– figure supplement 1F-Q). We obtained lists of cells belonging to each of these sub-clusters using [*WhichCells*] and used these lists to define new cell clusters in the full dataset, thus generating 17 clusters in total (Figure 5A).

### Finding markers for clusters in the embryonic and young adult optic lobe glial datasets

We identified marker genes expressed by each glial cluster using [*FindMarkers*]. To find genes with highly specific expression, we focused on the top 30 differentially expressed transcripts. We then examined their expression within and across clusters [*FeaturePlot*] to choose transcripts that were expressed by most cells in a given cluster (See Figures 4B and 5B for a summary of the marker genes validated *in vivo*). We used [*FindAllMarkers*] (default parameters, except only.pos = TRUE) to obtain the most highly expressed genes for the young adult optic lobe glia (See Source data file 3).

### *In vivo situ* Hybridisation chain reaction (HCR) probe design

To assess the expression of marker genes *in vivo*, we designed hybridization chain reaction (HCR) probes against chosen marker genes (Figure 5B). We designed 6-21 antisense probe pairs against each target gene, tiled along the annotated transcripts but excluding regions of strong sequence similarity to other transcripts (See Source data file 4), with the corresponding initiator sequences for amplifiers B3 and B5 (Choi et al., 2018). We purchased HCR probes as DNA oligos from Thermo Fisher (at 100 μM in water and frozen).

### *In situ* Hybridisation chain reaction (HCR)

We dissected optic lobe – central brain complexes from adult flies (female and male) in 1X phosphate-buffered saline (PBS). We then fixed them in 4% formaldehyde for 35 min at room temperature. We then rinsed (3X) and washed (3X 30mins) them in PBSTx (1X PBS with 0.5% Triton X-100, Fisher BioReagents). Next, we transferred the optic lobe – central brain complexes to 1.5 mL Eppendorf tubes and followed the Multiplexed HCR RNA-FISH protocol (Detection and Amplification stages) for whole-mount fruit fly embryos, from Molecular Instruments (molecularinstruments.com), also described in Choi et al. (2018) with the following modifications: we used 2 pmol of stock probe set and 12 pmol of each hairpin. Before proceeding with the 30min washes with 5X sodium chloride sodium citrate (SSC) with 0.1% Tween 20 (SSCT), we incubated samples with in 5X SSCT (with DAPI) for 2h at room temperature. All HCR buffers and hairpins were purchased from Molecular Instruments. Samples were stored at 4°C and mounted in SlowFade™ Gold Antifade Mountant (Thermo Fisher) within 3 days of completing the protocol.

### Microscopy and image processing

#### Larval VNC

We used a Zeiss LSM700 point-scanning or an Intelligent Imaging Innovations (3i) spinning disc confocal microscope with a 63X objective to image all samples with a step size of .2-.38μm, and the following laser lines: 405, 488, 555, 647. Images were acquired using Zen Black or Slidebook, respectively. Samples were imaged to encompass the entire dorsal-ventral extent of the VNC.

#### Young adult optic lobe

We used a Zeiss LSM800 or LSM880 confocal microscope with 20X or 40X objectives to image whole optic lobes with a maximum step size of 1μm. Individual astrocytes for morphological quantifications were imaged on a Zeiss LSM980 confocal microscope with AiryScan 2. Briefly, we acquired z-stacks (step size 3-4 μm) of whole optic lobes containing isolated MCFO-labelled astrocytes to determine astrocyte cell body position (using anti-Repo) along with information on the neuropil layers occupied by individual astrocytes (Using anti-Brp). We then acquired z-stacks (optimal step size) of individual astrocytes at high resolution.

### Quantifications and statistical analyses

#### Embryonic astrocyte morphological quantifications

To quantify the volume of type 1 versus type 2 astrocytes, we used Bitplane Imaris (version 10.0.0) to render 3D constructions of individual astrocyte MCFO clones (genotype: *hsMCFO + alrm-Gal4*). We used the *[Surfaces]* function with default parameters to measure total cell volume. Primary branches were quantified manually by segmenting astrocytes, 5 microns at a time, using the Imaris [*Ortho Slicer*] function.

#### Adult optic lobe astrocyte volumetric quantifications

We used Bitplane Imaris (version 9.9.0) to render 3D reconstructions of individual astrocytes. We used the [*Surfaces*] function with default parameters to measure total cell volume. We then aligned cells using the [*Frame*] function and cut the object (the cell) by neuropil layers (binning 2 layers together), and measured the volume for each was section of the cell. We quantified 8 cells of each astrocyte morphological category. Primary branches were quantified manually by segmenting astrocytes as detailed above. We used Graphpad Prism 9 to analyse the data for statistical significance as indicated in the main text.

#### Larval CNS

In Bitplane Imaris (version 9.6.1), a 3D projection was created for each individual VNC. For quantification of *repo*+ cells in the VNC versus brain lobes, we used the Imaris “Spots” function to automatically reconstruct and quantify the number of *repo*+ nuclei (xy diameter set to 3µm, z spread set to 5 µm, manual thresholding). We quantified *repo*+ cells in 7 independent VNC samples and 7 independent brain lobe samples at hatching (Genotype: *hsMCFO + fne-Gal4*). VNC enrichment was then determined via a one-way ANOVA in GraphPad Prism 9. For morphometric analyses, subtypes and morphotypes were quantified manually, blinded to genotype, and assessed statistically in Graphpad Prism 9 by Chi-squared statistical tests.

#### Adult optic lobe astrocyte cluster validation

We used the following approach to quantify the proportion of total astrocytes expressing a particular marker gene and their positions within the optic lobe. In Fiji (ImageJ2 version 3.2.0) we chose 35 to 50 optical slices [*Make Substack*] from each Z-stack. We performed a standard background subtraction [*Subtract Background*] (default parameters). We then created an average intensity projection and measured the mean fluorescence intensity of the HCR probe channel to obtain a Normalisation value (see below). We then used Icy (version 2; icy.bioimageanalysis.org) for further analysis. In Icy, we defined neuropil regions (as in Figures 6 and 7 for astrocytes, we then used the *Spot Detector* plugin (spot scale of 13 or 25 pixels in size and 40 to 100 sensitivity) to identify glial nuclei marked by nuclear GFP (*R86E01-Gal4>nls::GFP*) within these neuropil regions. We manually removed nuclei from the central brain that were detected by this method to restrict our analysis to the medulla, lobula and lobula plate only. We then quantified the mean fluorescence intensity of mRNAs detected by HCR in the defined ROIs. We scaled these measures to the normalization value (defined above) and used a threshold of 2.25, above which cells were considered positive for expression of the specific transcript examined. We settled on this threshold empirically by analysing several samples for markers that these cells were known to express or not express. We then calculated the proportion of nuclei positive for a given transcript in each optic lobe (excluding the lamina) as well as the proportion of positive nuclei within different neuropil regions. We used GraphPad Prism 9 to analyse these data with Chi-squared statistical tests.

### Gene Ontology Enrichment Analyses

All gene ontology (GO) enrichment analyses were carried out in R studio (version 1.4.1717), using R (version 4.1.1). We identified differentially expressed genes in our cluster/s of interest using [*FindMarkers*] from Seurat (version 4.1.1) (Hao et al., 2021). For comparisons of one single cluster to another, markers were selected with a log2FC ≥ 0.25 (fold change of 1.2). When comparing larger groups of clusters (*i.e.* the pooled lamina glia clusters to the pooled general glia clusters) a more stringent threshold of 1 log2FC (fold change of 2) was used to select differentially expressed genes. GO enrichment analysis for Biological Processes (BP) were carried out using [enrichGO] from clusterProfiler (version 4.2.2) (Hao et al., 2021). Within this function the adjusted p-values associated with the GO terms were calculated, using the Benjamini-Hochberg (BH) adjustment method for multiple comparisons. Only terms with adjusted p-value < 0.05 were considered. We next used [*simplify*] from clusterProfiler (Yu et al., 2012), with a p-adjust threshold of 0.7, which acts to reduce the redundancy in the enriched GO terms. The top 20 GO terms for each group were selected based on their significance (adjusted p-value), and plotted as a heatmap using ggplot2 (version 3.3.5)(Wickham, 2016). A dendrogram of the enriched GO terms was built using the Ward D agglomerative method with [*hclust; dist*], based on their pairwise similarity calculated with the Wang method using [*pairwise_termsim*] from the enrichplot package (version 1.14.2). This dendrogram was used to arrange and group the GO terms, enabling us to manually define superterms for the enrichment. For several poorly annotated GO terms we examined the annotated genes and manually renamed the GO terms (see R script available on our GitHub page for full details).

## Data and software availability

All raw and processed transcriptome data for the embryonic dataset are available from NCBI GEO (accession GSE208324). The scripts used to process the raw RNA-seq data and extract neuronal and glial clusters from the embryonic dataset are available at https://github.com/AustinSeroka/2022_stage17_glia. All other scripts, including midline glia annotation from the whole embryonic dataset, cleaned-up and annotation of embryonic glial dataset, as well as the integration, cleaned-up and annotation of young adult optic lobe glial dataset are available at https://github.com/VilFernandesLab/2022_DrosophilaGlialAtlas. The cleaned-up and annotated embryonic glial dataset and the integrated cleaned-up and annotated young adult optic lobe glial dataset are included here as source data files 5 and 6, respectively.

## Funding

SW was funded by a Wellcome Investigator Award (104682/Z/14/Z). SDA was funded by the National Institute of Health (K99/R00NS121137). VMF was funded by a Wellcome Trust and the Royal Society Sir Henry Dale Research Fellowship (210472/Z/18/Z).

## Acknowledgements

We thank Gaynor Smith, Marc Amoyel, Kelly Monk, Nathan Woodling, Simon Sprecher, and Chris Doe for helpful comments and critiques of the manuscript. Stocks for this study were obtained from the Bloomington Drosophila Stock Center.

## Author contributions

I.L-B., S.D.A, and V.M.F conceived of the project. I.L-B. performed and analysed experiments related to Figures 2-9, and all associated figure supplements. S.D.A performed and analysed experiments related to Figure 1, Figure 4 and all associated figure supplements. M.C. performed and analysed experiments related to Figures 2, 8 and all associated figure supplements. M.C. and I.L-B also generated the graphics used in Figures 1-2 and associated figure supplements. A.S. analysed experiments related to Figure 4, as well as Supplementary file 3, Figure 3– figure supplement 2, and Figure 3-figure supplement 3. C.T., G.P. and S.W. designed HCR probes and assisted in training I.L-B. and M.C. in HCR hybrid chain ISH. I.L-B., S.D.A., M.C., and V.M.F. wrote the paper and prepared the figures. All authors commented and approved of the manuscript.

---

**Figure 1 – figure supplement 1:**
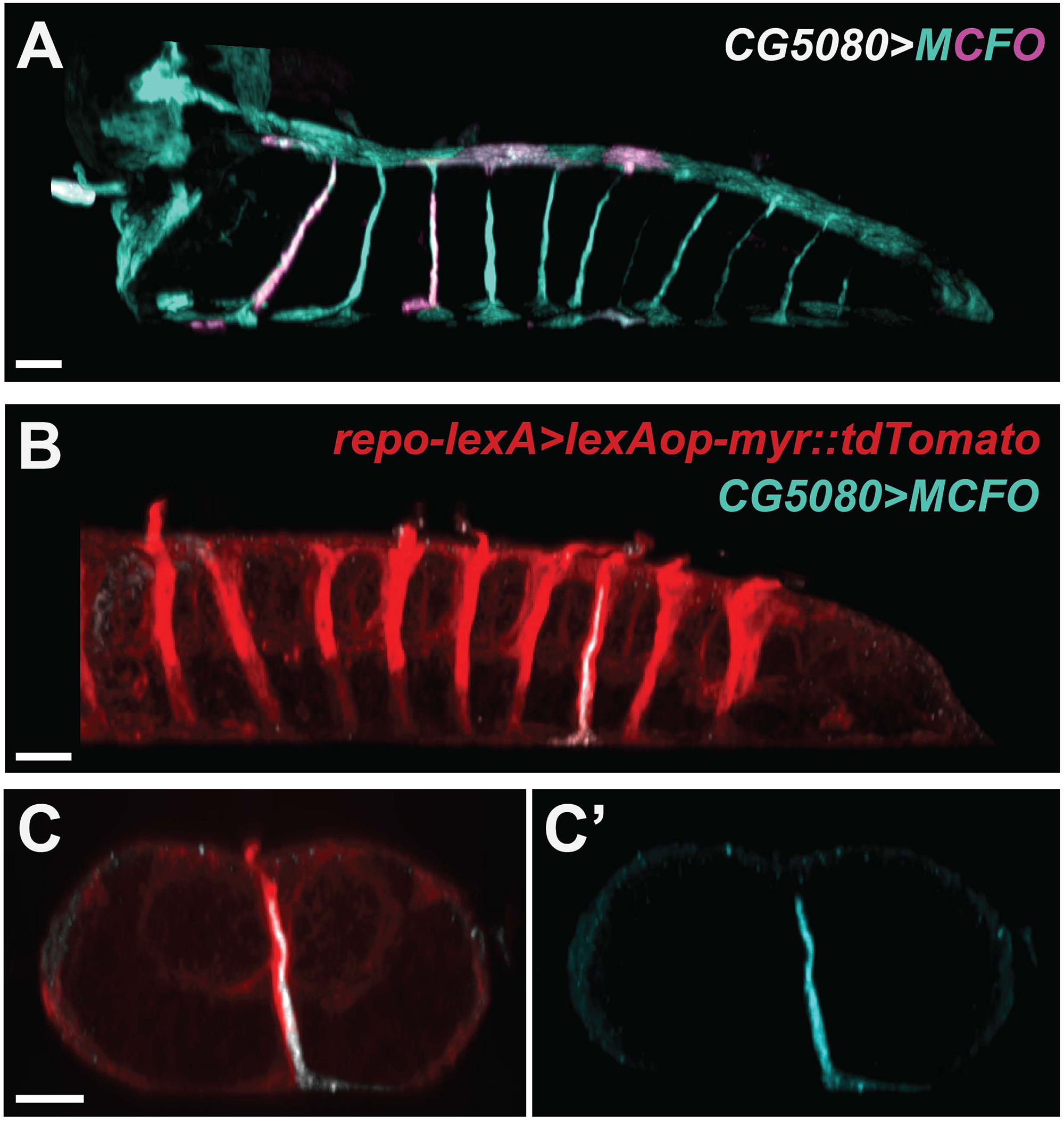
Channel perineurial glia characterisation. **(A)** Lateral view of the VNC at 0 h after larval hatching showing channel perineurial glia (cyan and magenta) on the ventral and dorsal surfaces, each sending a single projection with ventral channel perineurial glia sending longer processes than their dorsal counterparts. **(B-C’)** Lateral view (B) and cross-sectional view (C) of the VNC at 0 h after larval hatching showing a single ventral channel perineurial clone (cyan), and all other glia in red. Note in (B) the red outer glial membranes along the channels that belong to enveloping channel subperineurial glia. Individual channel perineurial glial cells have their surface domains along one side or the other of the midline (C). All scale bars represent 10 µm.

**Figure 1 – figure supplement 2:**
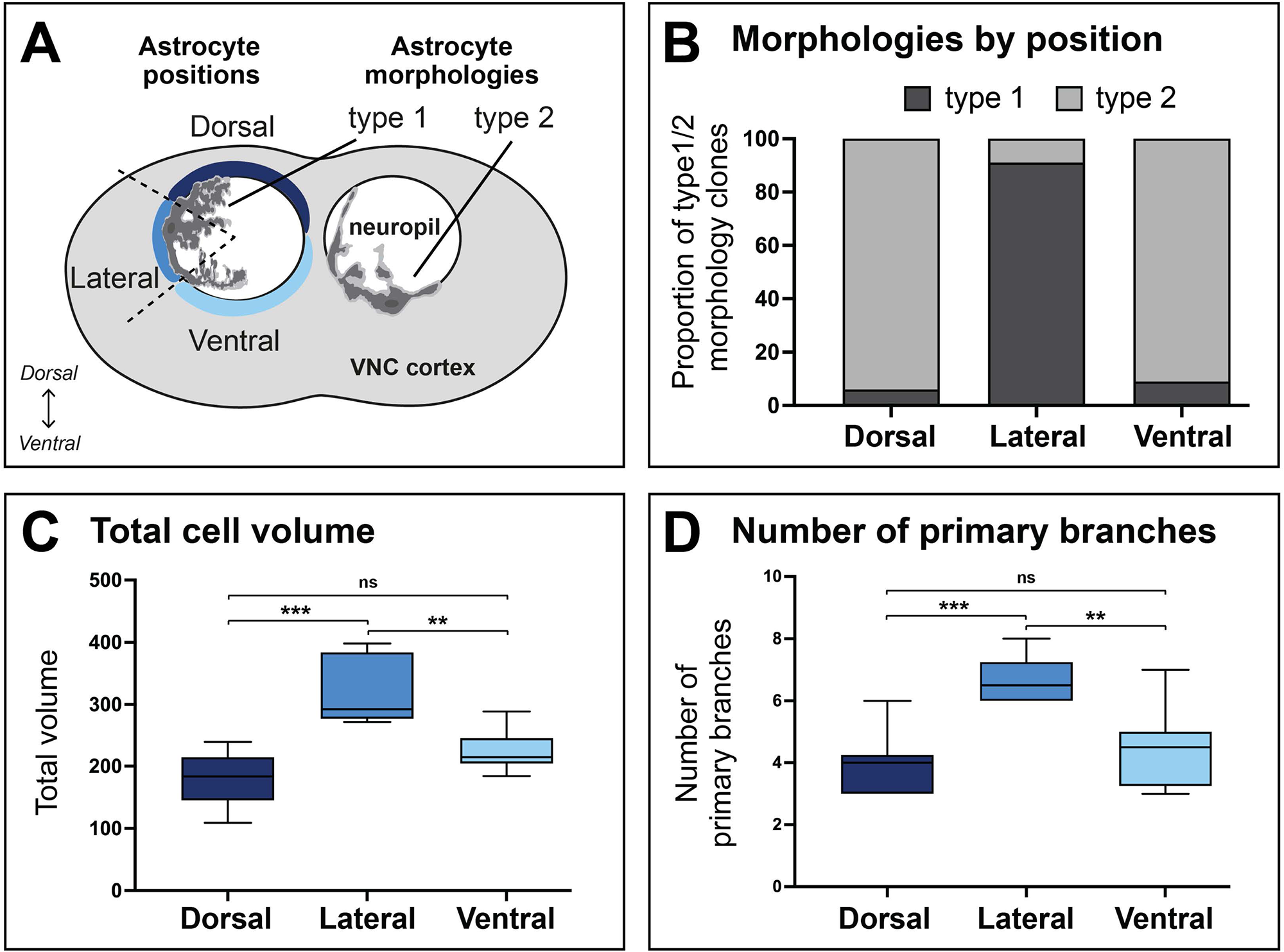
Quantifications of newly-hatched VNC astrocyte morphologies. **(A)** Schematic of the cross-section of the embryonic VNC showing the different astrocyte positions and morphologies. Position types are Dorsal, Lateral and Ventral based on nucleus position, separated by dashed lines (left). Type 1 morphologies are more arborised and type 2 are less arborised. Morphologies described originate from sparse *alrm>MCFO* clones. **(B)** Proportions of type 1 and type 2 morphologies for astrocyte clones at 0 h after larval hatching with nuclei in Dorsal, Lateral, or Ventral positions, as defined in (A). Type 1 astrocytes are more prevalent laterally, while type 2 astrocytes are more prevalent in dorsal and ventral positions. N=117 clones from N=36 brains. **(C-D)** Boxplots of total cell volumes (C) and number of primary branches (D) for astrocyte clones in the dorsal, lateral and ventral positions. Line in the middle represents the mean and limits are minimum and maximum. The lateral astrocytes, mainly type 1, show higher volumes and more primary branches compared to dorsal and ventral astrocytes, mainly type 2. N≥6 clones from N ≥6 brains per neuropil position. Mann-Whitney U-test *p* values indicated (ns, non-significant; ** *p* > 0.01; *** *p* > 0.001).

**Figure 1 - Video 1. 3D projection of newly-hatched VNC lateral astrocyte.**

360°-view of a single laterally-positioned astrocyte (*alrm>MCFO*) clone at 0 h after larval hatching.

**Figure 1 - Video 2. 3D projection of newly-hatched VNC ventral astrocyte.**

360°-view of a single ventrally-positioned astrocyte (*alrm>MCFO*) clone at 0 h after larval hatching.

**Figure 1 - Video 3. 3D projection of newly-hatched VNC dorsal astrocyte.**

360°-view of a single dorsally-positioned astrocyte (*alrm>MCFO*) clone at 0 h after larval hatching.

**Figure 2 – figure supplement 1.**
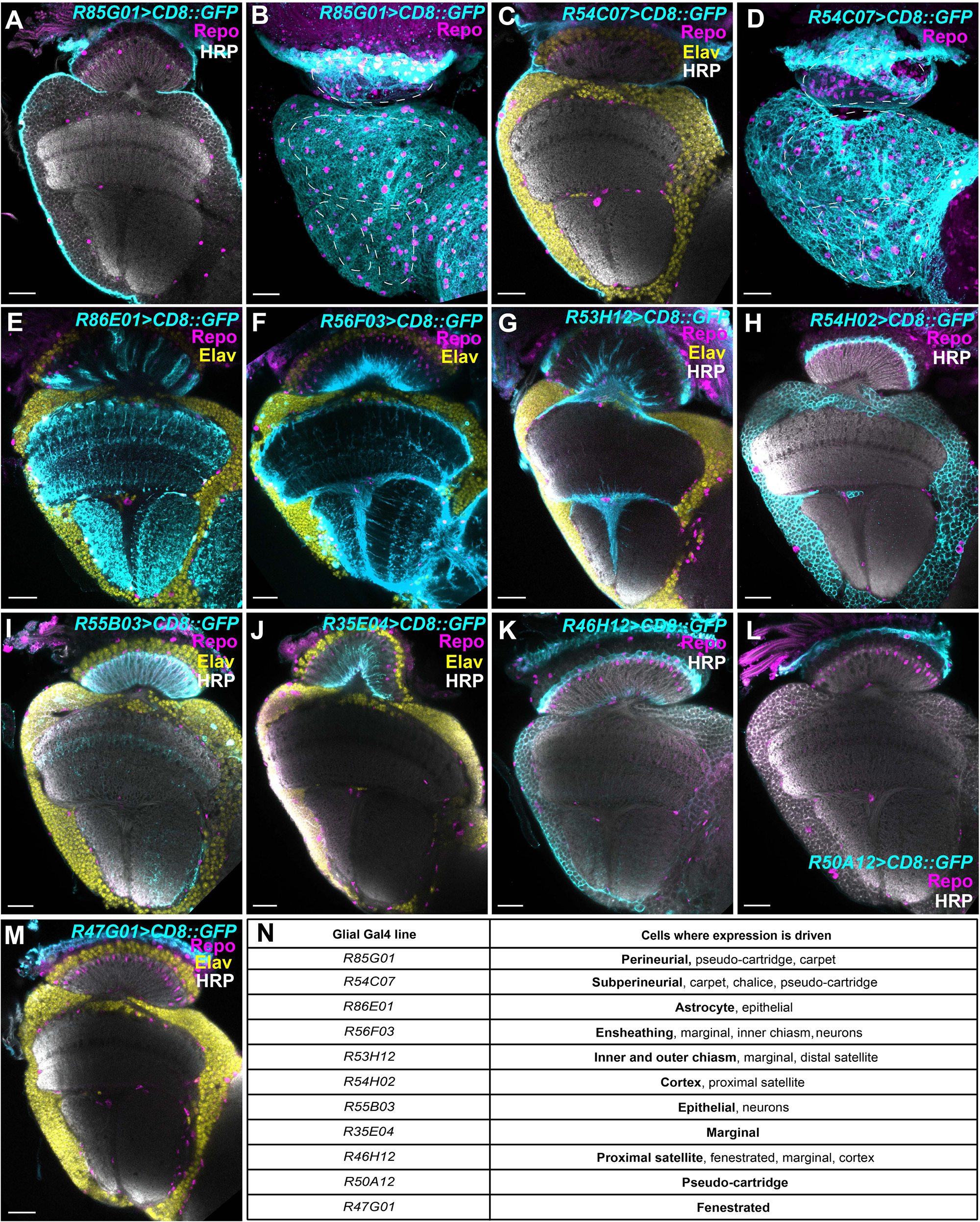
Expression patterns of the glial Gal4s drivers used to evaluate glial morphology in the *Drosophila* adult optic lobe. **(A-M)** GFP expression driven by the indicated glial Gal4 driver (described previously by Kremer et al. (2017) for the **(A,B)** optic lobe perineurial glia, **(C,D)** optic lobe subperineurial glia, **(E)** optic lobe astrocyte glia, **(F)** optic lobe ensheathing glia (not including chiasm glia; also drives expression in a subset of neurons), **(G)** chiasm glia (and marginal glia), **(H)** medulla, lobulla, and lobula plate cortex glia and proximal satellite glia, **(I)** lamina astrocytes (epithelial glia), **(J)** lamina ensheathing glia (marginal glia), **(K)** lamina-specific cortex glia (proximal satellite glia), **(L)** lamina subperineurial glia (pseudo-cartridge glia), and **(M)** lamina perineurial glia (fenestrated glia). **(N)** A table outlining the glial Gal4 lines and the glial subtypes that they drive expression within. Where relevant, cyan marks CD8::GFP driven by the glia-Gal4, magenta marks Repo, yellow labels Elav and HRP labels the neuropils in white. Panels B and D are maximum projections showing the surface of the optic lobe. Dashed lines outline the neuropils. All scale bars represent 20 µm.

**Figure 2 – figure supplement 2:**
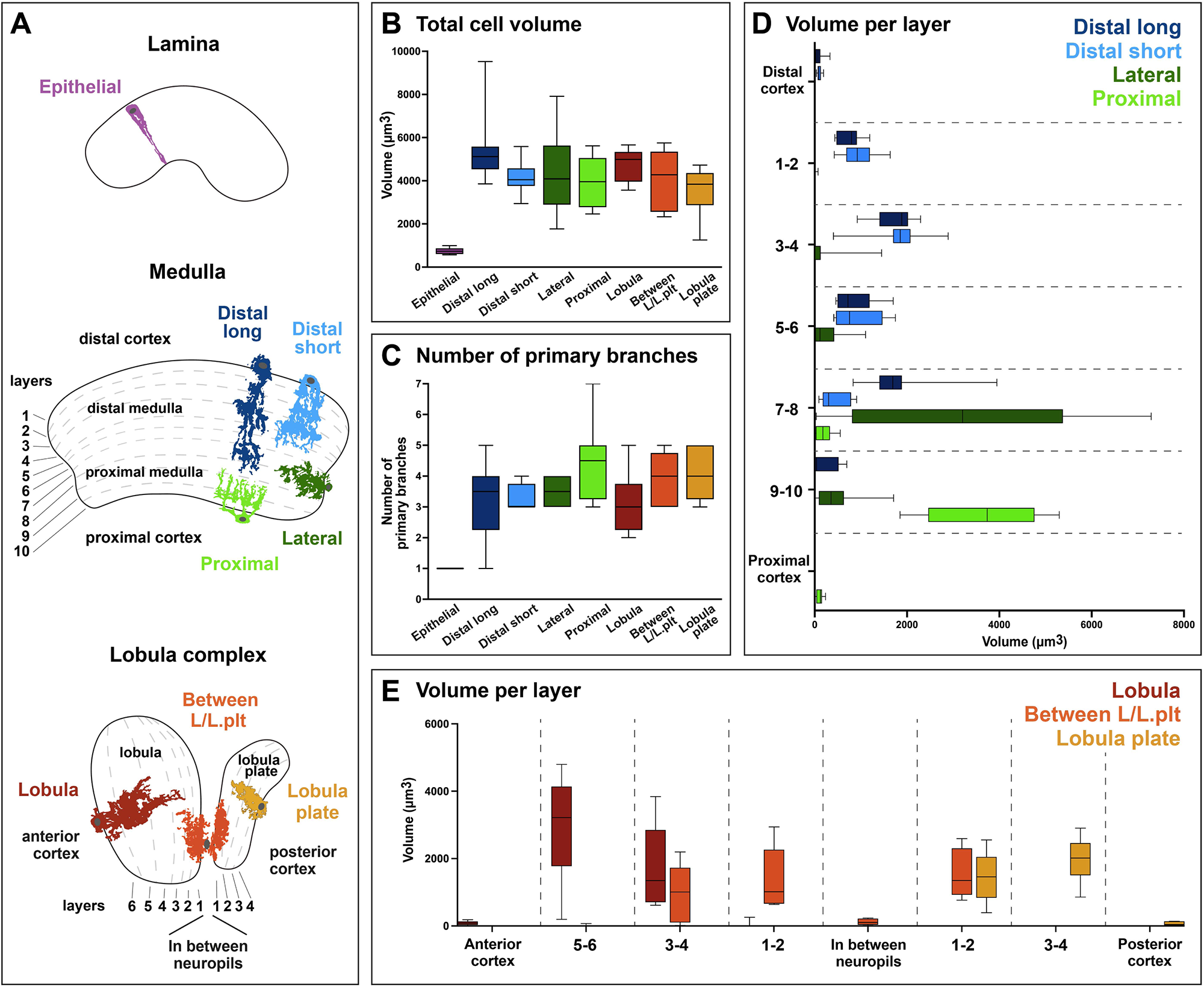
Morphological quantifications of adult optic lobe astrocytes. **(A)** Schematic of the cross-section of the adult optic lobe and its four neuropils: lamina, medulla, lobula and lobula plate, with the different astrocyte morphologies indicated. Morphologies described originate from sparse*Astrocyte(R86E01)>MCFO* clones. Dashed lines indicate the neuropil layers. **(B-C)** Boxplot of total cell volumes (B) and number of primary branches (C) for each astrocyte morphology. Line in the middle represents the mean and limits are minimum and maximum. N=8 clones. **(D-E)** Boxplot of cell volumes within each neuropil layer pair for each astrocyte morphology of the medulla (D) and lobula complex (E). Line in the middle represents the mean and limits are minimum and maximum. N=8 clones.

**Figure 2 - Video 1. 3D projection of an adult epithelial glia (lamina astrocyte)**

360°-view of a single epithelial glia (*repo>MCFO*) clone in a young adult (1-3 days old).

**Figure 2 - Video 2. 3D projection of an adult long distal medulla astrocyte.**

360°-view of a single long distal medulla astrocyte (*R86E01>MCFO*) clone in a young adult (1-3 days old).

**Figure 2 - Video 3. 3D projection of an adult short distal medulla astrocyte.**

360°-view of a single long short medulla astrocyte (*R86E01>MCFO*) clone in a young adult (1-3 days old).

**Figure 2 - Video 4. 3D projection of an adult lateral medulla astrocyte.**

360°-view of a single lateral medulla astrocyte (*R86E01>MCFO*) clone in a young adult (1-3 days old).

**Figure 2 - Video 5. 3D projection of an adult proximal medulla astrocyte.**

360°-view of a single proximal medulla astrocyte also known as a chandelier glia (*R86E01>MCFO*) clone in a young adult (1-3 days old).

**Figure 2 - Video 6. 3D projection of an adult lobula-only astrocyte.**

360°-view of a single lobula-only astrocyte (*R86E01>MCFO*) clone in a young adult (1-3 days old).

**Figure 2 - Video 7. 3D projection of an adult lobula-lobula plate astrocyte.**

360°-view of a single lobula-lobula-plate astrocyte (*R86E01>MCFO*) clone in a young adult (1-3 days old).

**Figure 2 - Video 8. 3D projection of an adult lobula-only astrocyte.**

360°-view of a single lobula plate-only astrocyte (*R86E01>MCFO*) clone in a young adult (1-3 days old).

**Figure 3 – figure supplement 1:**
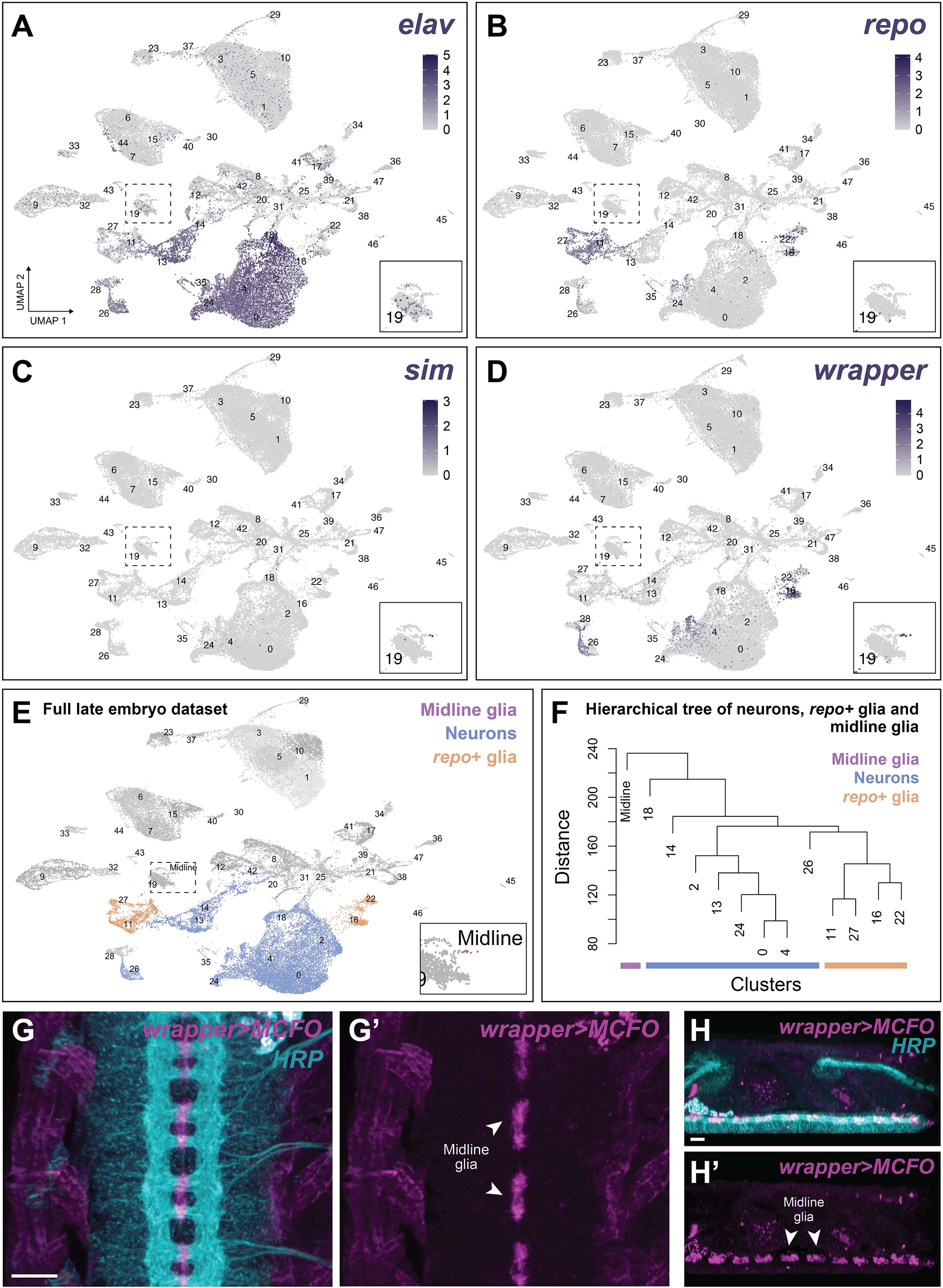
Annotation of the cluster corresponding to the midline glia within the whole embryo dataset. **(A-D)** Expression levels of *elav* (A)*, repo* (B)*, sim* (C) and *wrapper* (D) plotted on the whole embryo UMAP. Each dot represents a single cell, and the colour represents the level of expression as indicated. Zoomed-in details of cluster 19 are shown. **(E)** UMAP of the whole embryo, indicating clusters defined as midline glia (purple), neurons (blue) and *repo+* glia (yellow), based on *elav, repo, sim* and *wrapper* expression. **(F)** Dendrogram of hierarchical clustering average expression of all genes between midline glia (purple), neurons (blue) and *repo+* glia (yellow) clusters. Midline glia form an outgroup to both neuron and *repo+* glia. **(G-H)** Single focal planes of MCFO clones (magenta) of midline glia generated with *wrapper-Gal4*. HRP in cyan. Scale bar is 10 µm.

**Figure 3 – figure supplement 2.**
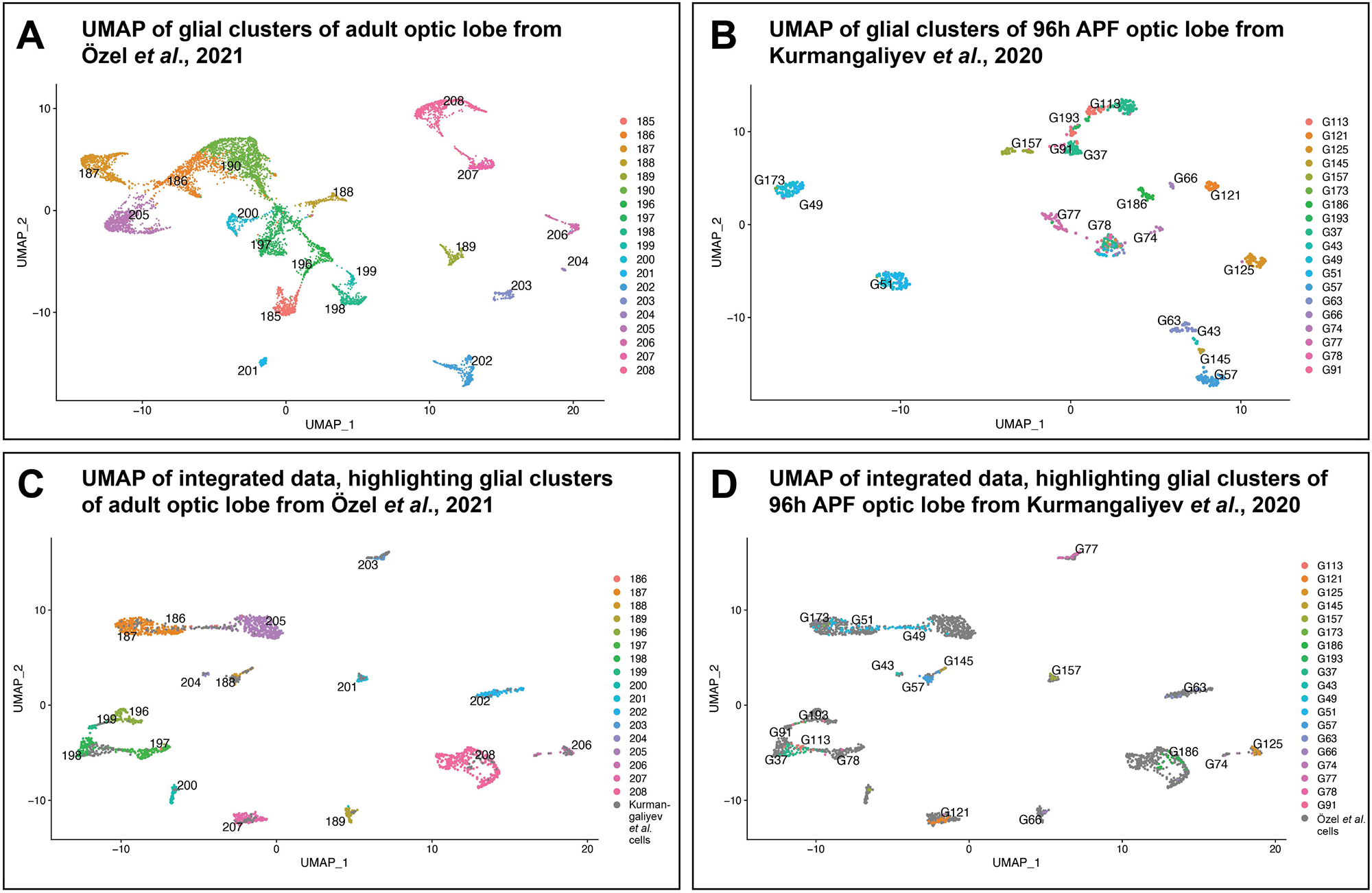
Contribution of the two datasets, 96h APF and 3-day-old adult, to the resulting integrated young adult optic lobe dataset. **(A)** UMAP of the 19 glial clusters from 3-day-old adult optic lobes, from Özel et al. (2021) **(B)** UMAP of the 19 glial clusters from 96h APF optic lobes, from Kurmangaliyev et al. (2020). **(C,D)** UMAP of the integrated young adult optic lobe dataset, highlighting the 3-day-old adult clusters (C) or 96h APF clusters (D) in colour, with 96h APF (C) and 3-day-old adult (D) cells in grey.

**Figure 3 – figure supplement 3.**
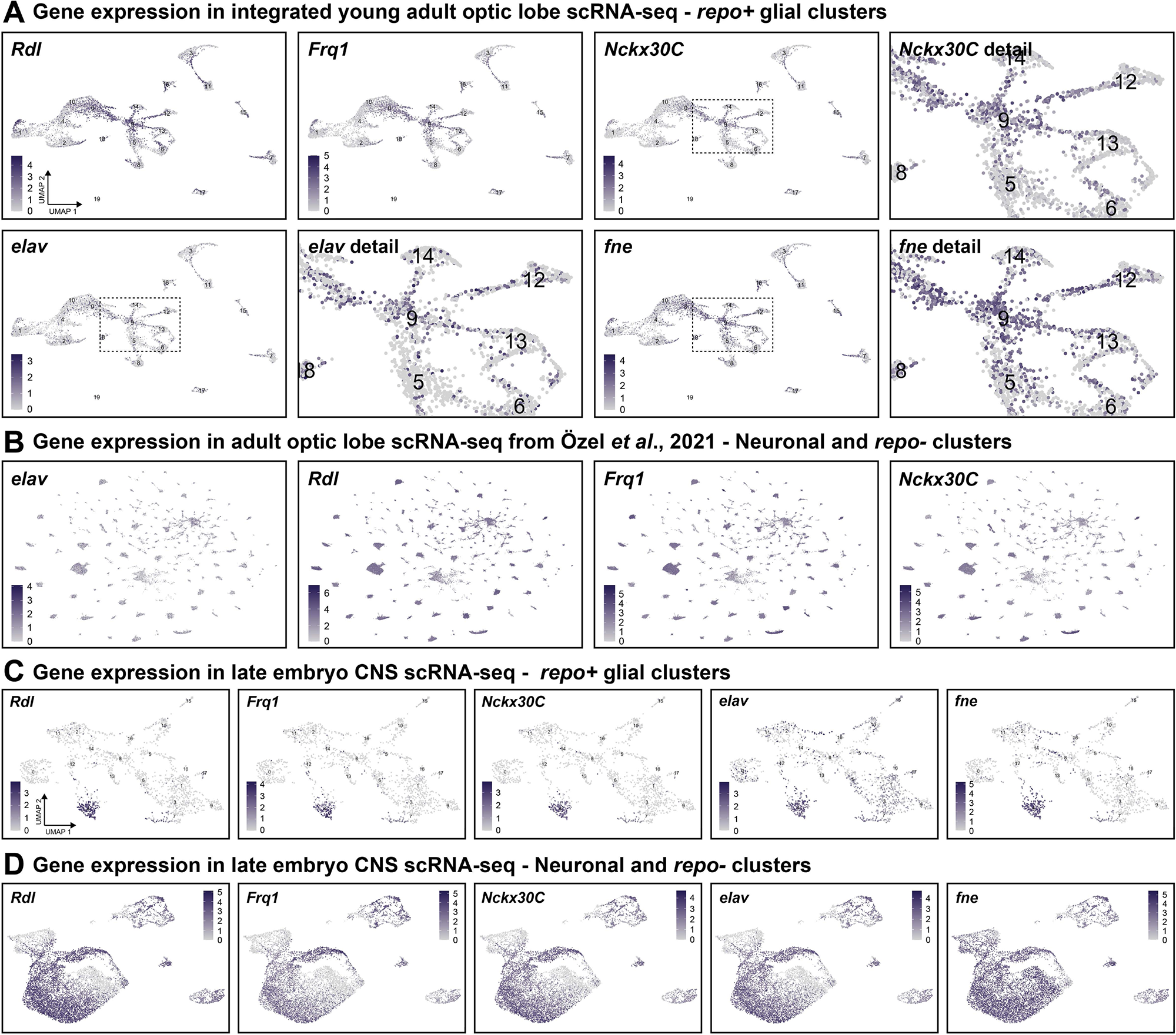
Contamination of glial clusters with cells with typical neuronal profile. **(A)** Expression levels of *Rdl, Frq1, Nckx30C, elav* and *fne* plotted on the young adult optic lobe integrated UMAP, before neuronal clean-up. Each dot represents a single cell, and the colour represents the level of expression as indicated. Zoomed-in details of the centre of the UMAP are shown for *Nckx30C, elav* and *fne*. **(B)** Expression levels of *elav, Rdl, Frq1* and *Nckx30C* plotted on the 3-day-old adult optic lobe UMAP, from Özel et al., 2021, including all clusters except the 19 glial clusters. All four genes showed expression in all clusters, illustrating the pan-neuronal nature of *Rdl, Frq1* and *Nckx30C* expression. **(C)** Expression levels of *Rdl, Frq1*, *Nckx30C* plotted on the embryonic glial UMAP, before neuronal clean-up. The expression of these three genes overlapped with the expression of *elav* and *fne*, mainly in cluster #4. **(D)** Expression levels of *Rdl*, *Frq1* and *Nckx30C elav* and *fne* plotted on the UMAP of the embryonic nervous system clusters.

**Figure 3 – figure supplement 4.**
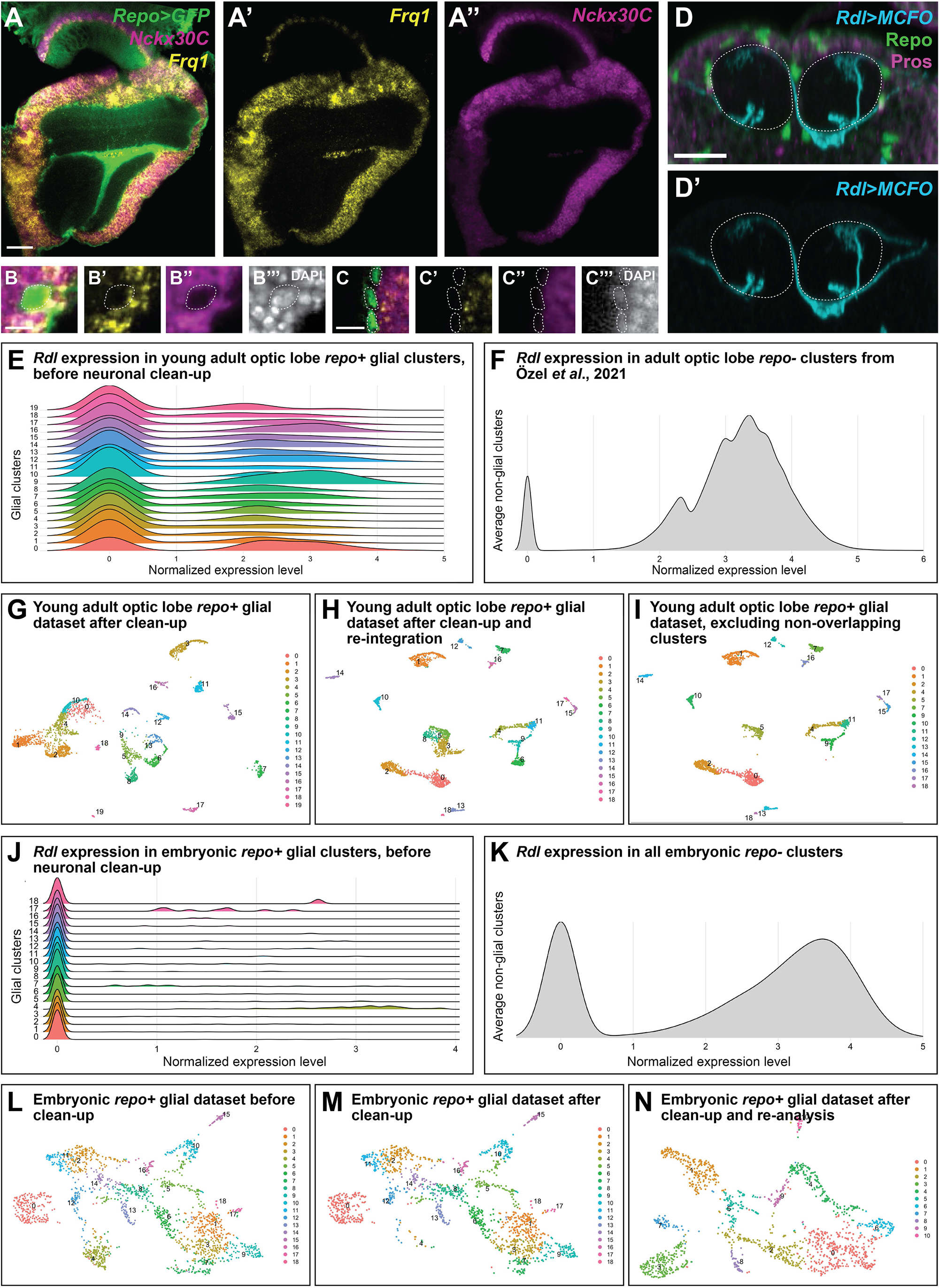
Clean-up of contamination of glial clusters with cells with typical neuronal profile. **(A)** *Frq1* (yellow) and *Nckx30C* (magenta) expression were detected by *in situ* HCR in the cortex area of the optic lobe. GFP labelled all glial cells in green. Single focal plane. Scale bar is 20 µm. **(B,C)** Single focal planes showing glial somas (green) with *Frq1* (yellow) and *Nckx30C* (magenta) expression. DAPI marks all nuclei in white. Dashed lines outline glial somas. Scale bars are 5 µm. **(D)** Single focal planes of MCFO clones (cyan) generated with *Rdl-Gal4*. Repo in green and Prospero in magenta. Dashed lines outline the neuropil. Scale bar is 7 µm. **(E)** The distribution of *Rdl* expression levels in each glial cluster of the young adult optic lobe integrated dataset, before neuronal clean-up. **(F)** The distribution of *Rdl* expression levels in all cells of the 3-day-old adult optic lobe UMAP, from Özel et al., 2021, except glial clusters. **(G)** UMAP of the 20 glial clusters obtained from the first integration of optic lobe datasets **(H)** UMAP of 19 glial clusters after clean-up of potential neurons by excluding cells with normalised expression of *Rdl, Frq1* and *Nckx30C* >1, and cells with *Hml* expression (>0 normalised expression) as potential hemocytes. **(I)** Cluster #3, #6 and #8 were excluded from the UMAP in (H) since less than 1% of the cells contained in them originated from the Kurmangaliyev et al. (2020) dataset (see Materials and Methods for details). **(J)** The distribution of *Rdl* expression in each glial cluster of the embryonic dataset, before neuronal clean-up. **(K)** The distribution of *Rdl* expression in all cells of the embryonic nervous system, excluding cells belonging to the 19 glial clusters. **(L)** UMAP of the 19 initial glial clusters of the embryonic nervous system, before neuronal clean-up. **(M)** Same UMAP as in (L) after exclusion of potential neurons (normalised expression >1 of *Rdl, Frq1* and *Nckx30C*) and potential hemocytes (normalised expression >0 of *Hml* expression). **(N)** UMAP of the remaining cells in (M) after reanalysis and reclustering (see Materials and Methods for details).

**Figure 4 – figure supplement 1.**
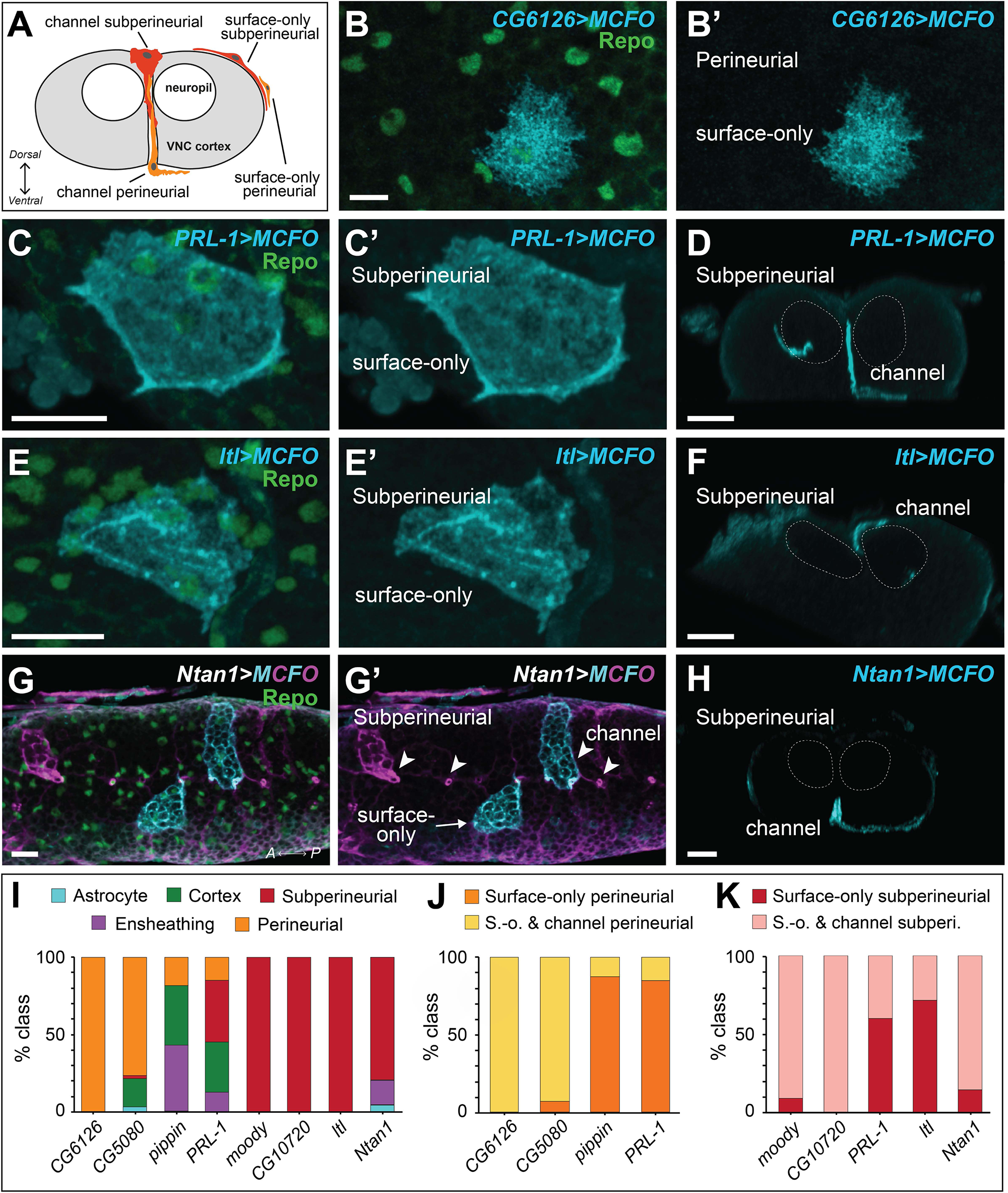
Validation and analysis of new markers for *Drosophila* VNC surface glia. **(A)** Schematic of the cross-section of the embryonic VNC showing the different surface glial types, channel and surface-only perineurial or subperineural glia. **(B-H)** Surface and cross-sectional views of MCFO clones at 0 h after larval hatching generated with the Gal4 lines indicated, belonging to marker genes with high expression in the surface glia clusters: *CG6126* (N=372 clones from N=11 brains), *PRL-1* (N=154 clones from N=17 brains), *ltl* (N=19 clones from N=7 brains) and *Ntan1* (N=68 clones from N=14 brains). All MCFO clones labelled in cyan and magenta, with Repo in green. **(I)** Quantification of the frequency of clones recovered by glial class for the indicated driver. **(J)** Quantification of perineurial glia morphotype frequency by driver line. Channel perineurial glia were detected in 100% of *CG6126* MCFO brains (N=11 brains total) and 92.9% of *CG5080* brains (N=13 brains total), compared to 7.7% of *pippin* MCFO brains (N=13 brains total) and 5.7% of *PRL-1* MCFO brains (N=15 brains total). **(K)** Quantification of subperineurial glia morphotype frequency by driver line. Channel subperineurial glia were detected in 90.9% of *moody* MCFO brains (N=12 brains total), 28.6% of *ltl* MCFO brains (N=7 brains total), 85.7% of *Ntan1* MCFO brains (N=15 brains total), 100% of *CG10702* MCFO brains (N=15 brains total), and 40% of *PRL* MCFO brains (N=15 brains total). Dashed lines outline the neuropil. All scale bars are 10 µm.

**Figure 4 – figure supplement 2.**
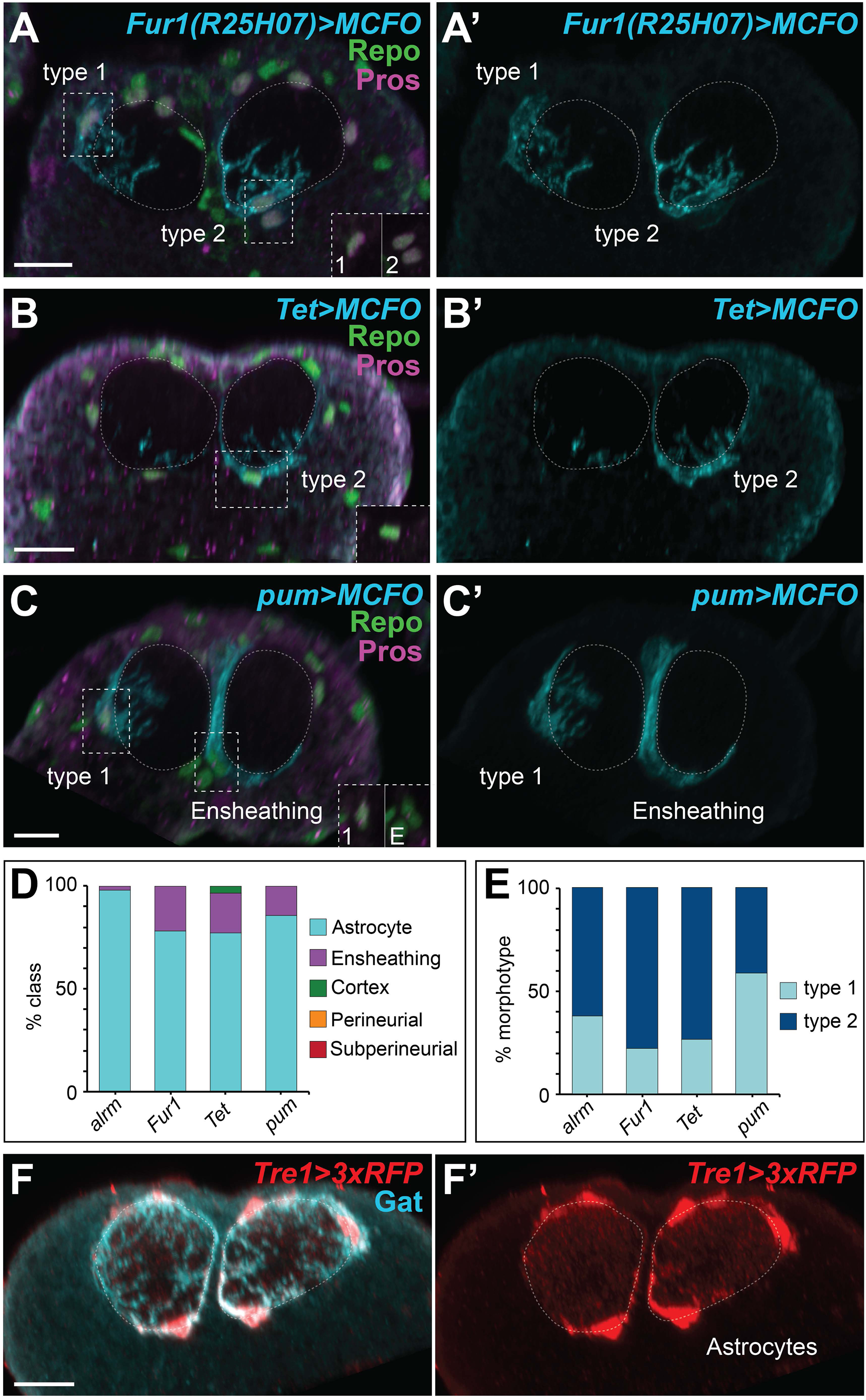
Validation and analysis of new markers for *Drosophila* VNC astrocytes. **(A-C)** MCFO clones (VNC cross sections) at 0 h after larval hatching generated with the Gal4 lines indicated, belonging to marker genes with high expression in the astrocyte cluster: *Fur1* (N=27 clones from N=9 brains), *Tet* (N=71 clones from N=10 brains), and *pum* (N=25 clones from N=9 brains). All MCFO clones labelled in cyan, with Repo in green and Prospero in magenta. Insets in (A-C) show Prospero and Repo in glial nuclei; only astrocyte clones were positive for Prospero. Dashed lines outline the neuropils. **(D)** Quantification of the frequency of glial type clone for each indicated driver line: *alrm* (N=117 clones from N=36 brains), other Ns noted above. **(E)** Quantification of astrocyte morphotype frequency by driver line. **(F)** VNC cross section showing colocalization of the astrocyte marker Gat (cyan) and a gene-trap line where 3xRFP has been swapped for the Tre1 locus (red). Dashed lines outline the neuropil. Scale bars are 10 µm.

**Figure 4 – figure supplement 3.**
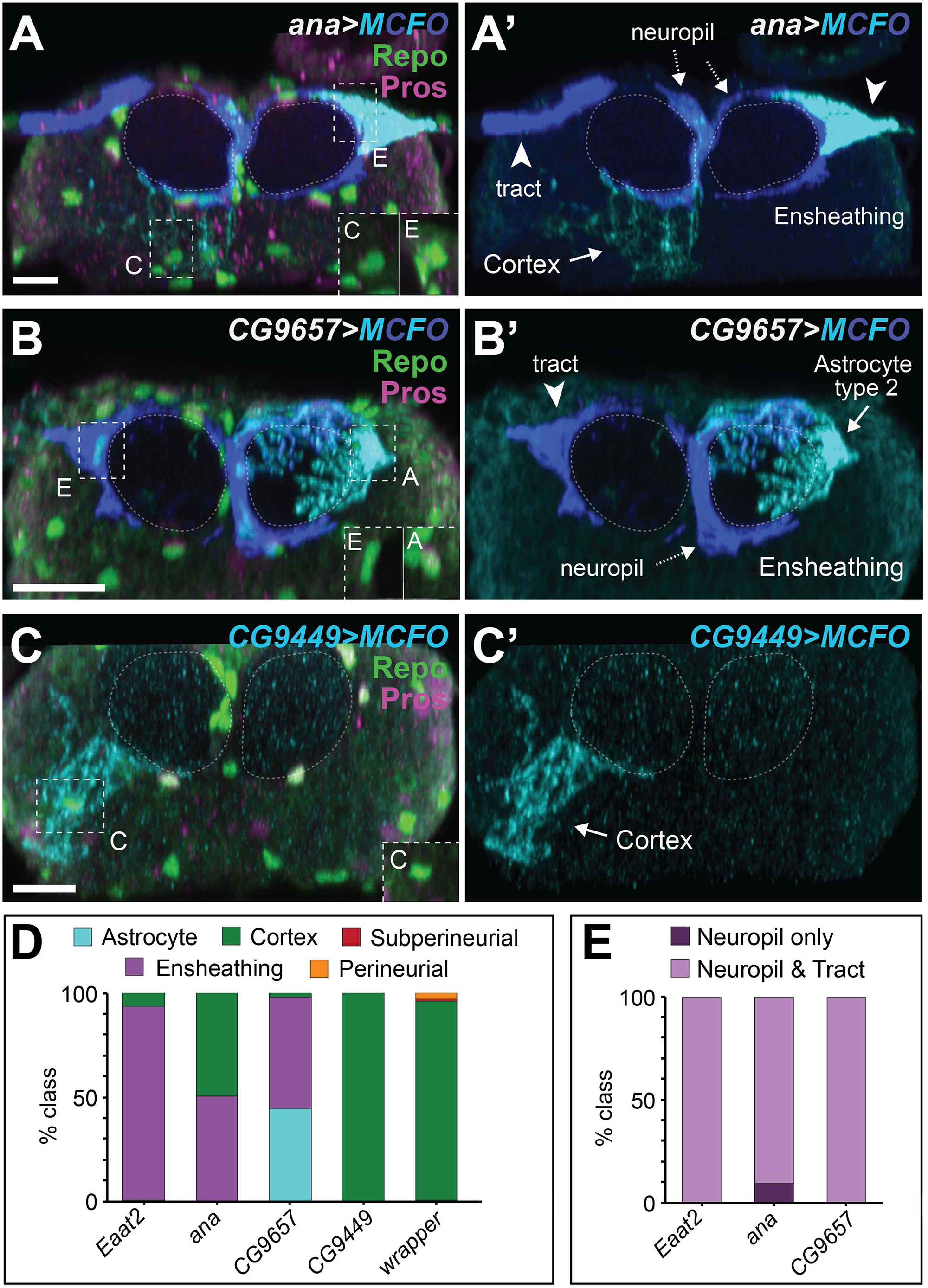
Validation and analysis of new markers for *Drosophila* VNC ensheathing and cortex glia. **(A-C)** MCFO clones (VNC cross sections) at 0 h after larval hatching generated with the Gal4 lines indicated, belonging to marker genes with high expression in the ensheathing or cortex clusters: *ana* (N=388 clones from N=11 brains), *CG9657* (N=287 clones from N=10 brains), and *CG9449* (N=5 clones from N=3 brains). Tract and neuropil ensheathing types are indicated. All MCFO clones labelled in cyan and blue, with Repo in green and Prospero in magenta. Insets in (A,B,C) show Prospero and Repo in glial nuclei, where only astrocyte clones were positive for Prospero. Dashed lines outline the neuropils. **(D)** Quantification of the frequency of glial type clone for each indicated driver line: *wrapper* (N=13 brains), other Ns noted above. **(E)** Quantification of the frequency of brains with clones of both ensheathing types, tract and neuropil, or only neuropil clones, for each indicated driver line. Dashed lines outline the neuropil. Scale bars are 10 µm.

**Figure 5 – figure supplement 1.**
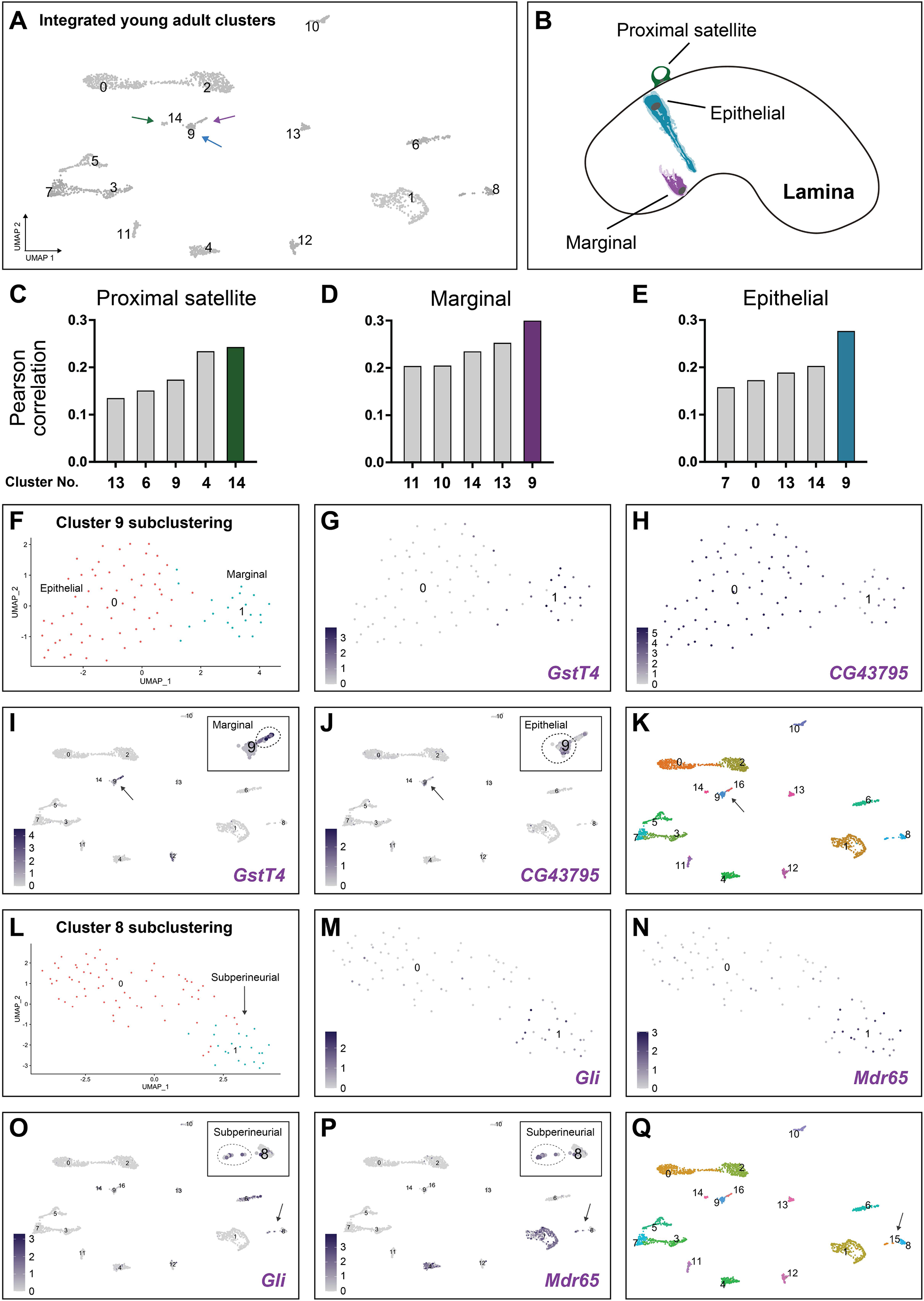
Subclustering of young adult optic lobe glial clusters to find hidden rare subtypes. **(A)** UMAP of the 15 glial clusters of the integrated young adult dataset. **(B)** Schematic of the adult optic lobe lamina with marginal, epithelial and proximal satellite glia represented. **(C-E)** Pearson correlation values between the bulk-RNA-seq transcriptomes of the proximal satellite, marginal and epithelial glia and the top five glial cluster matches from the young adult dataset. **(F)** Subclustering of cluster #9 generated two subclusters. **(G)** *GstT4* (a gene highly expressed marginal glia bulk RNA-seq data) showed high expression in subcluster #1. **(H)** *CG43795* (a gene highly expressed in epithelial glia bulk RNA-seq) showed high expression in subcluster #0. **(I,J)** Expression of GstT4 (I) and CG43795 (J) in the UMAP of the 15 glial clusters of the integrated young adult dataset. Both genes showed mutually exclusive expression patterns within cluster #9. **(K)** UMAP showing the original cluster #9 split into new clusters #9 and #16 (see Materials and Methods). **(L)** Subclustering cluster #8 generated two subclusters. **(M,N)** *Gli (Gliotactin)* and *Mdr65 (Multi drug resistance 65)*, both known markers of subperineurial glia (Mayer et al., 2009), were expressed in subcluster #1. **(O,P)** Expression of *Gli* and *Mdr65* on the UMAP of the 16 glial clusters of the integrated young adult dataset. Both genes showed overlapped expression in a group of cells belonging to cluster #8. **(Q)** UMAP showing the original cluster #8 split into new clusters #8 and #15 (see Materials and Methods).

**Figure 5 – figure supplement 2.**
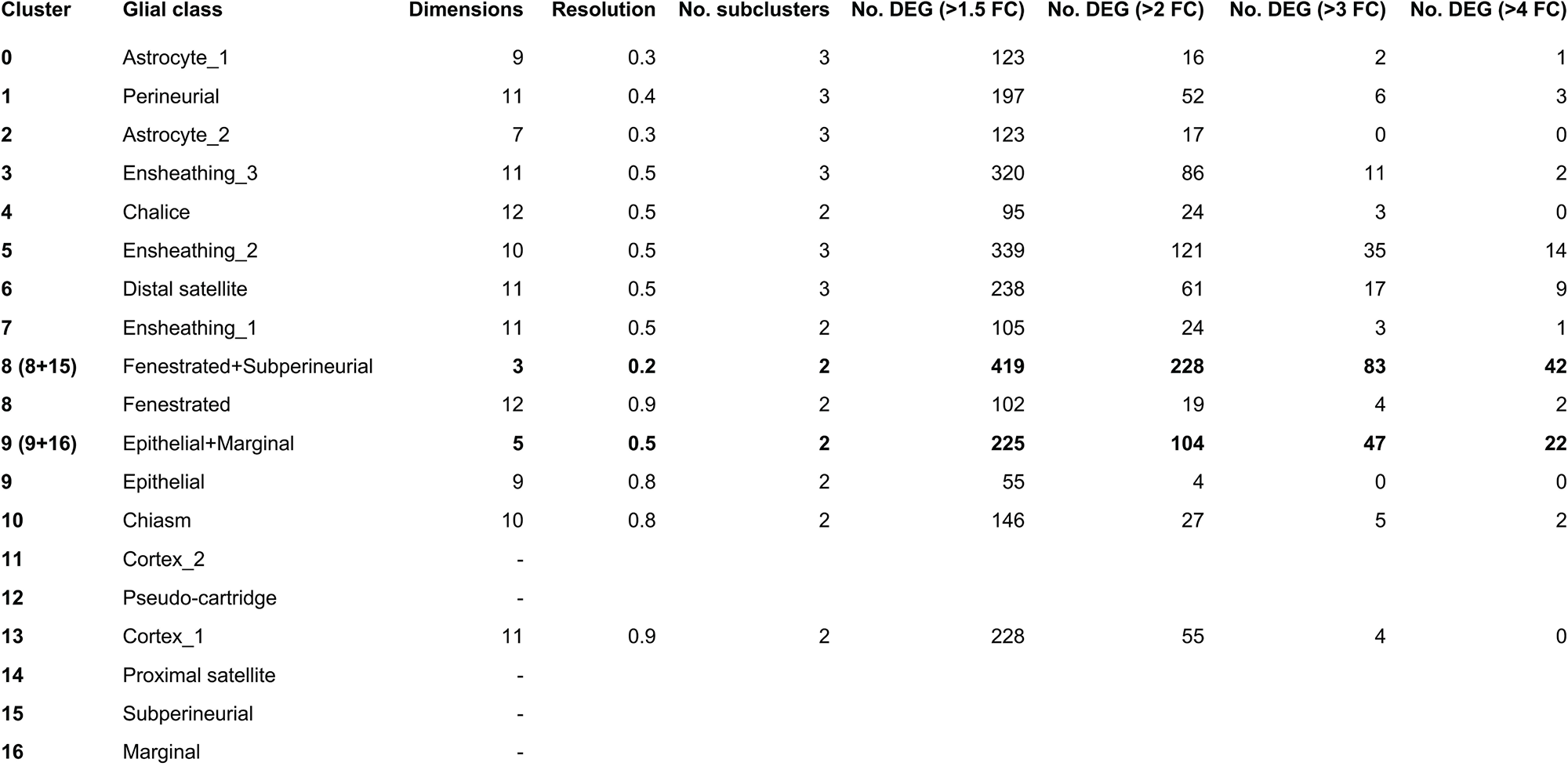
Summary of the number of differentially expressed genes (DEGs) obtained for different fold-change (FC) cutoffs between subclusters within each of the 15 glial clusters post clean-up of the young adult glial dataset.

**Figure 5 – figure supplement 3.**
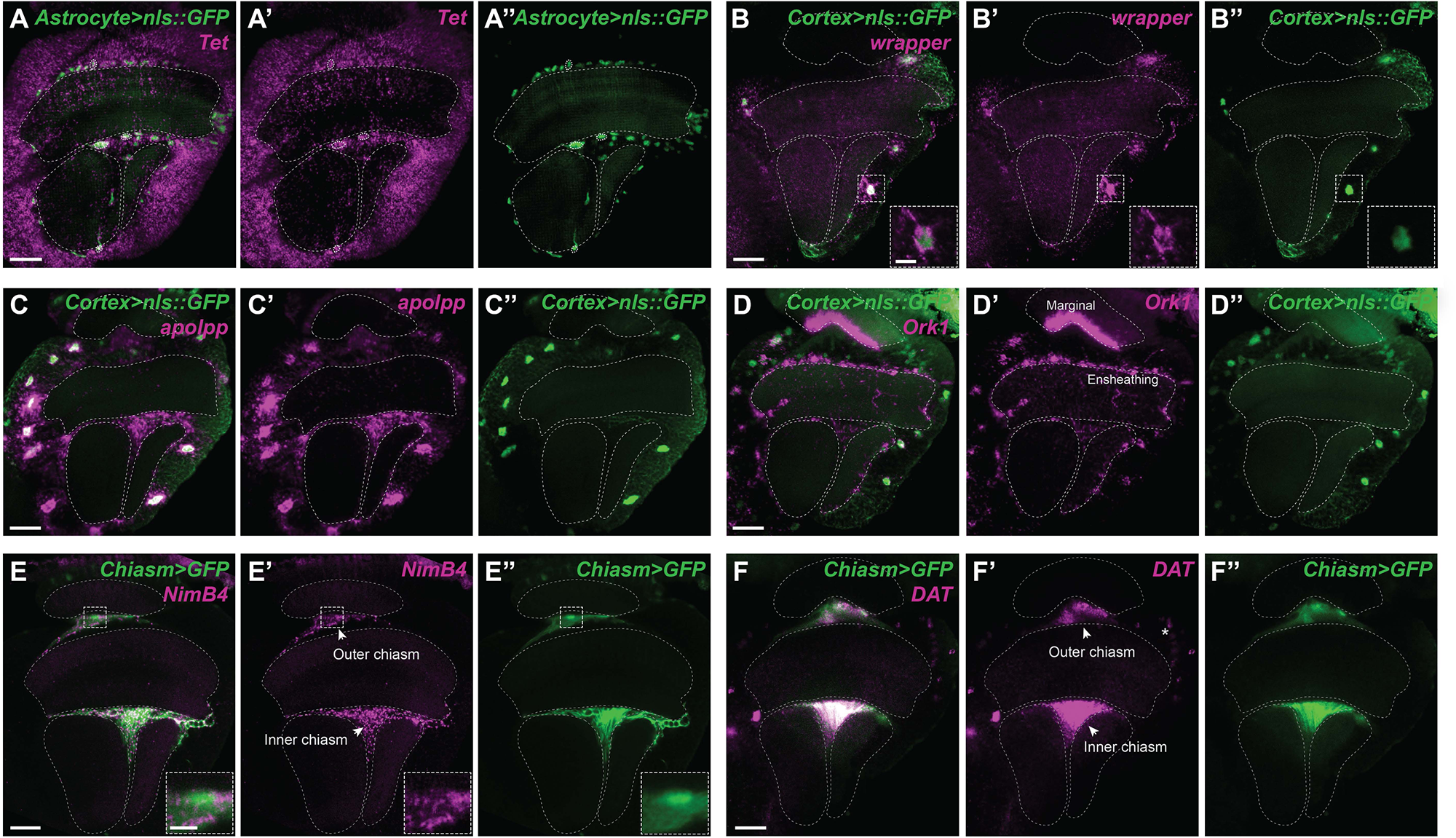
Validation of marker genes for astrocyte, cortex and chiasm glia of the young adult optic lobe. **(A)** *Astrocyte(R86E01)>nls::GFP* adult optic lobe where *Tet (Ten-Eleven Translocation (TET) family protein)* expression (magenta) was detected by *in situ* HCR throughout the cortex, including in many GFP positive astrocyte (green) nuclei (outlined by thicker dashed lines). See Figure 5 and 6 for additional *in vivo* astrocyte marker gene validation. **(B)** *Cortex(R54H02)>nls::GFP* adult optic lobe where *wrapper* expression (magenta) was detected by *in situ* HCR in many GFP positive cortex glia (green) nuclei. Inset of zoomed in nuclei (scale bar is 5 µm). **(C)** *Cortex(R54H02)>nls::GFP* adult optic lobe where *apolpp (apolipophorin)* expression (magenta) was detected by *in situ* HCR in most GFP positive (green) nuclei. A few nuclei adjacent to the neuropil were also positive (likely ensheathing glia based on the scRNA-seq data). **(D)** *Cortex(R54H02)>nls::GFP* adult optic lobe where *Ork1 (Open rectifier K+ channel 1)* expression (magenta) was detected by *in situ* HCR in some GFP positive cortex glia (green) nuclei. Many nuclei adjacent to the neuropil were also positive (likely ensheathing glia based on the scRNA-seq data), as well as in marginal glia. **(E)** *Chiasm(R53H12)>GFP* adult optic lobe where *NimB4 (Nimrod B4)* expression (magenta) was detected by *in situ* HCR in most GFP positive chiasm glia (green) nuclei. Inset of zoomed in nuclei (scale bar is 5 µm). **(F)** *Chiasm(R53H12)>GFP* adult optic lobe where *DAT (Dopamine transporter)* expression (magenta) was detected by *in situ* HCR in most GFP positive chiasm glia (green) nuclei. Some nuclei in the cortex region around the medulla were positive for *DAT* (asterisk), consistent neuronal expression in the scRNA-seq data from Özel et al., 2021. Single focal planes in (A,E,F) and maximum projections of 11-12 focal planes (1 μm each) in (B-D). Dashed lines outline the neuropils and scale bars are 20 µm.

**Figure 5 – figure supplement 4.**
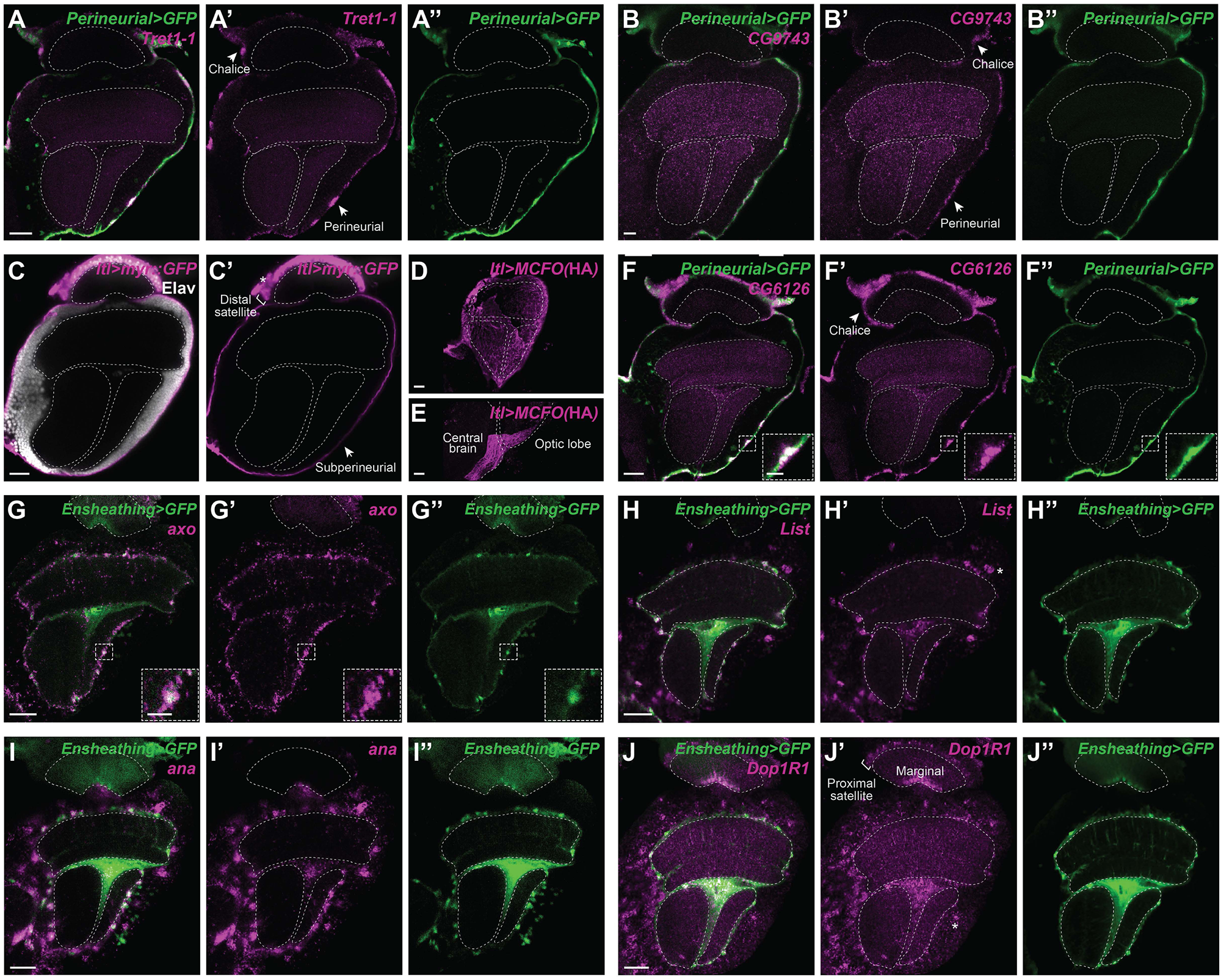
Validation of marker genes for surface and ensheathing glia of the young adult optic lobe. **(A,B)** *Perineurial(R85G01)>GFP* adult optic lobe where **(A)** *Tret1-1 (Trehalose transporter 1-1)* and **(B)** *CG9743* expression (magenta) was detected by *in situ* HCR. For both genes, most GFP positive (green) nuclei of the general perineurial glia (surrounding the medulla, lobula and lobula plate) were positive, in addition to the chalice glia (as predicted by the scRNA-seq data). *Tret1-1* was shown to be expressed in perineurial glia (Volkenhoff et al., 2015). **(C)** Adult optic lobes expressing myrGFP driven by *ltl(larval translucida)-Gal4*, gene trap Trojan line, showing *ltl* expression in general surface glia (arrow), distal satellite (bar), and lamina surface (asterisk; fenestrated glia as predicted by the scRNA-seq data). **(D,E)** Maximum projection of *ltl-*Gal4 labelled MCFO clones showing subperineurial morphology (see Figure 2). **(F)** *Perineurial(R85G01)>GFP* adult optic lobe where *CG6126* expression (magenta) was detected by *in situ* HCR. Most GFP positive (green) nuclei of the general perineurial glia were positive, in addition to the chalice and fenestrated glia, as predicted by the scRNA-seq data. **(G)** *Ensheathing(R56F03)>GFP* adult optic lobe where *axo* (axotactin) expression (magenta) was detected by *in situ* HCR in most GFP positive (green) nuclei adjacent to the medulla, lobula and lobula plate neuropils. Inset of zoomed-in nuclei (scale bar is 5 µm). **(H)** *Ensheathing(R56F03)>GFP* adult optic lobe where *List* expression (magenta) was detected by *in situ* HCR in most GFP positive (green) nuclei adjacent to the medulla, lobula and lobula plate neuropils, and in some GFP negative nuclei in the cortex area of the same neuropils (asterisk) (general cortex glia as predicted by the scRNA-seq data). **(I)** *Ensheathing(R56F03)>GFP* adult optic lobe where *ana* (*anachronism*) expression (magenta) was detected by *in situ* HCR in most GFP positive (green) nuclei adjacent to the neuropil as well as many nuclei in the cortex area, which were probably cortex glia based on the large size of the nuclei (predicted by scRNA-seq data). **(J)** *Ensheathing(R56F03)>GFP* adult optic lobe where *Dop1R1 (Dopamine 1-like receptor 1)* expression (magenta) was detected by *in situ* HCR in many GFP positive (green) nuclei adjacent to the medulla, lobula and lobula plate neuropils as well as nuclei in the cortex area of the medulla, lobula and lobula plate, marginal glia and proximal satellite glia (predicted by scRNA-seq data). All panels are single focal planes unless stated otherwise. Dashed lines outline the neuropils and scale bars are 20 µm.

**Figure 5 – figure supplement 5.**
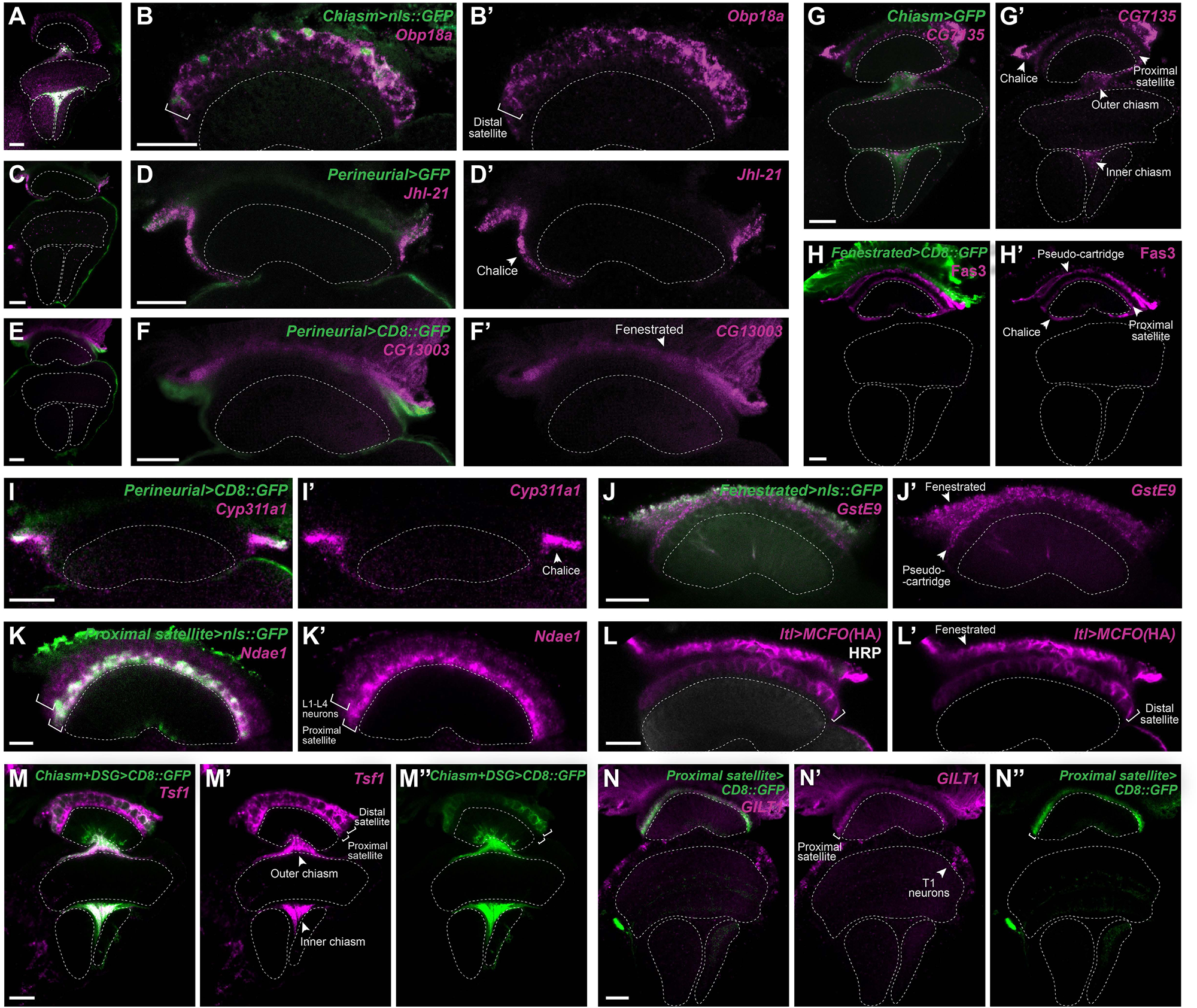
Validation of marker genes for lamina-specific glial subtypes of the young adult optic lobe. **(A,B)** *Chiasm(R53H12)>GFP* adult optic lobe, showing expression in chiasm and distal satellite glia. *Obp18a* expression (magenta) was detected by *in situ* HCR in most GFP positive (green) nuclei of chiasm and distal satellite glia. **(C,D)** *Perineurial(R85G01)>GFP* adult optic lobe where *JhI-21 (Juvenile hormone Inducible-21)*expression (magenta) was detected by *in situ* HCR, specifically in chalice glia. **(E,F)** *Perineurial(R85G01)>CD8::GFP* adult optic lobe where CG13003 expression (magenta) was detected by *in situ* HCR in fenestrated glia. **(G)** *Chiasm(R53H12)>GFP* adult optic lobe where *CG7135* expression (magenta) was detected by *in situ* HCR in chiasm, chalice, and proximal satellite glia. **(H)** Fas3 (magenta) antibody staining of *Fenestrated(R47G01)>CD8::GFP* adult optic lobe, showed expression in the lamina surface glia layer below the fenestrated glia, pseudo-cartridge glia, in the chalice, and proximal satellite glia. **(I)** *Perineurial(R85G01)>CD8::GFP* adult optic lobe where *Cyp311a1 (Cytochrome P450 311a1)* expression (magenta) was detected by *in situ* HCR, specifically in chalice glia. **(J)** *Fenestrated(R47G01)>nls::GFP* adult optic lobe where *GstE9 (Glutathione S transferase E9)* expression (magenta) was detected by *in situ* HCR in fenestrated and pseudo-cartridge glia. **(K)** *Proximal satellite(R46H12)>nls::GFP* adult optic lobe where *Ndae1 (Na+-driven anion exchanger 1)* expression (magenta) was detected by *in situ* HCR in proximal satellite glia, as well as in L1, L2, L3 and L4 lamina neurons, as predicted by the neuronal scRNA-seq data from Özel et al., 2021. **(L)** Large MCFO clones labelled by *ltl-Gal4*, marked distal satellite (bar) and fenestrated glia (arrow). HRP in white. **(M)** *Chiasm(R53H12)>CD8::GFP* adult optic lobe where *Tsf1* expression (magenta) was detected by *in situ* HCR in proximal and distal satellite and chiasm glia. **(N)** *Proximal satellite(R46H12)>CD8::GFP* adult optic lobe where *GILT1* expression (magenta) was detected by *in situ* HCR in proximal satellite glia, as well as in T1 medulla neurons, as predicted by the neuronal scRNA-seq data from Özel et al., 2021. All panels are single focal planes. Dashed lines outline the neuropils and scale bars are 20 µm.

**Figure 6 – figure supplement 1.**
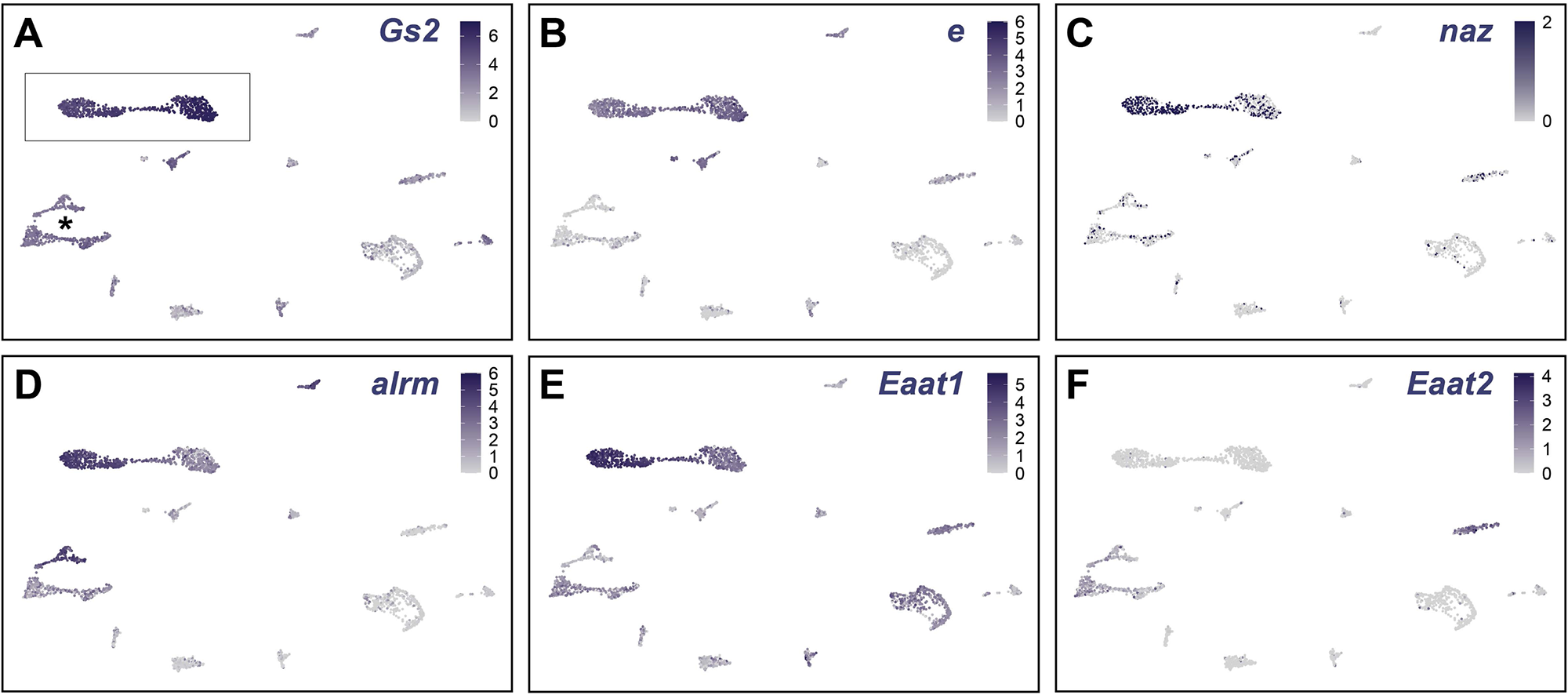
Expression levels of known astrocytic markers, and *Eaat2* ensheathing marker, in the young adult optic lobe glial clusters. **(A-C)** Plots showing *Gs2*, *e*, and *naz* expression levels in young adult optic lobe glial clusters. Each dot represents a single cell, and the colour represents the level of expression as indicated. As previously described (Kato et al., 2020), *Gs2* was expressed in both astrocytes (clusters indicated in a box in A) and ensheathing (clusters indicated with asterisk in A), while *e* and *naz* are expressed exclusively in astrocytes. **(D)** *alrm*, a known astrocyte marker (Edwards et al., 2012), showed expression in the astrocyte clusters (box in A). **(E)** *Eaat1*, a known astrocyte marker (Peco et al., 2016), showed expression in the astrocyte clusters (box in A). **(F)** *Eaat2*, a known ensheathing marker (Peco et al., 2016), showed expression in the ensheathing clusters (asterisk in A) and no expression in the astrocyte clusters (box in A).

**Figure 6 – figure supplement 2.**
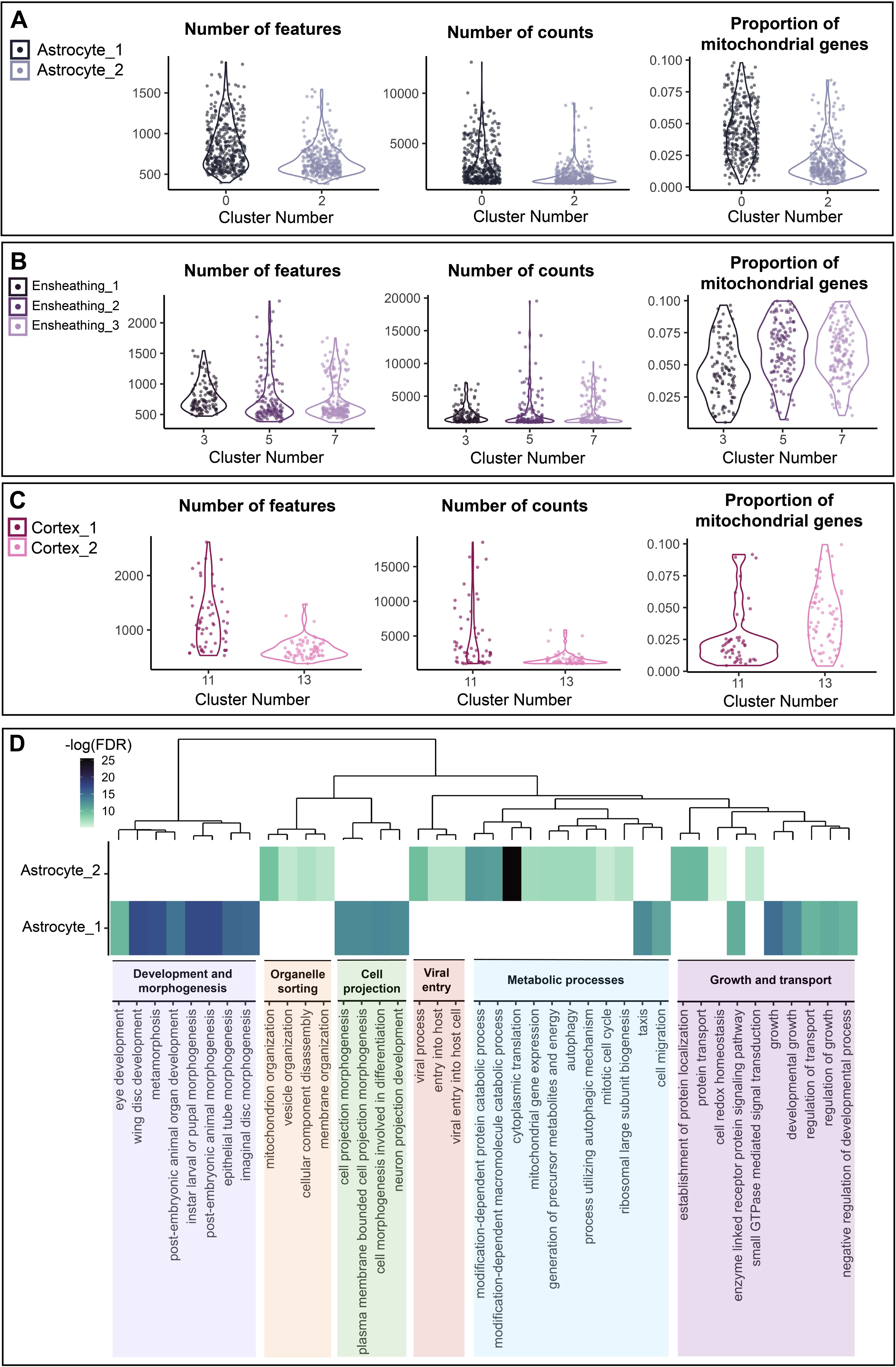
Assessment of the transcriptome quality of ‘multiplet’ clusters. **(A-C)** Violin plots outlining the number of features, number of counts and proportion of mitochondrial genes, for the astrocyte **(A)**, ensheathing **(B)**, and cortex **(C)** glia cluster multiplets. Cluster identity is colour-coded and the cluster number is indicated on the x-axes. **(A)** Astrocyte multiplets showed no overall trend in transcriptome quality. **(B)** Ensheathing glia showed a decrease in transcriptome quality in clusters #5 and #7, compared to #3. **(C)** Cortex glia cluster #13 appears to be of a lower quality than #11. **(D)** GO analysis indicating the enriched biological processes in Astrocyte_1 (cluster #0) and Astrocyte_2 (cluster #2).

## Supplementary file 1

This file is a classification key to distinguish between Embryonic Repo+ glial classes and morphological categories based on association with different brain regions.

## Supplementary file 2

This file is a classification key to distinguish between adult optic lobe glial classes, subclasses and morphological categories based on association with different brain regions.

## Supplementary file 3

### Examining batch effects between technical and biological replicates for embryonic scRNA-seq

**(A)** Schematic of data analysis for stage 17 whole embryo. 10X libraries were prepared from four biological replicates, each sequenced across two separate lanes. Following examination for batch effects **(B-F)** using the 8 sets of resulting CellRanger outputs, CellRanger Aggregate was used to merge the data. The resulting CellRanger outputs (Sarah_aggr_trial1_barcodes.tsv, Sarah_aggr_trial1_features.tsv, Sarah_aggr_trial1_matrix.mtx) were then read into Seurat for downstream processing (see Materials and Methods).

**(B)** Schematic of data analysis to check for batch effects between sequencing lanes. As this analysis predates Seurat integration functionalities, the stage 17 embryo data generated from sequencing lane 1 (biological replicates 1-4) and sequencing lane 2 (biological replicates 1-4) were combined into 2 Seurat objects, merged and clustered **(C)**. The number of cells derived from each sequencing lane were calculated for each cluster in UMAP space, and the correlation between the two sequencing lanes was calculated (R^2^ = 0.995) (see “220706 glia paper supp fig 1.Rmd”).

**(D)** Lane-of-origin was plotted onto the clusters in UMAP space, showing an even distribution across all clusters. Together, these analyses demonstrate the absence of batch effects between sequencing lanes.

**(E)** Schematic of data analysis for examining batch effects between biological replicates. Reads for each biological replicate were combined across both sequencing lanes. These four objects were then merged using Seurat, and clustered using standard methods. Biological-replicate-of-origin for each cell was plotted onto the clusters in UMAP space. Cells derived from each of the four biological replicates contributed equally to the neuronal (illustrated by *elav* expression) and glial (illustrated by *repo* expression) clusters of the UMAP **(F)**, demonstrating the absence of batch effects across the replicates.

## Supplementary file 4

List of specific genotypes and conditions used by figure panel

## Source data file 1

List of genes showing enriched expression in midline glia relative to all other embryonic cells

## Source data file 2

Lists of genes showing enriched expression in each embryonic *repo+* glial cluster relative to the others

## Source data file 3

Lists of genes showing enriched expression in each adult glial cluster relative to the others

## Source data file 4

Lists of HCR probe sequences used in this study

**Supporting Zip Documents:**

- Source data file 5
- Source data file 6

## Notes

https://www.ncbi.nlm.nih.gov/geo/query/acc.cgi?acc=GSE208324

https://github.com/VilFernandesLab/2022_DrosophilaGlialAtlas

https://www.ncbi.nlm.nih.gov/geo/query/acc.cgi?acc=GSE142787

https://www.ncbi.nlm.nih.gov/geo/query/acc.cgi?acc=GSE156455

https://www.ncbi.nlm.nih.gov/geo/query/acc.cgi?acc=GSE116969

https://github.com/AustinSeroka/2022_stage17_glia

### Competing Interest Statement

The authors have declared no competing interest.

### Summary of Updates

The main suggestions from the reviewers related to: (1) Providing a more thorough evaluation of the new surface glial cell type to determine its identity as a perineurial vs. subperineurial glia while amending terminology to be consistent with previous reports. (2) Helping non-specialists know how to categorize cells by providing quantifications of cell shape. We are grateful to the reviewers for their constructive feedback. We have incorporated their suggestions and addressed their concerns in our revised manuscript. We provide a detailed response outlining the changes we have made in the attached Response to Reviewers document. To summarise, we confirmed the identity of the new surface glial type as belonging to the perineurial glial class but also identified an additional subperineurial glial subclass, which had been described previously in the literature based on lineage relationships, nuclear positioning and molecular markers, but whose morphology had not been described previously. These findings add clarity to embryonic glial classification and will be very valuable for the community. We also quantified astrocyte morphologies in the embryonic VNC and the adult optic lobe in detail, provide high resolution 3-dimensional reconstructions of the various astrocyte morphological categories and include classification keys to help readers distinguish between glial classes and morphological categories. All these changes have substantially improved the quality of the manuscript and further support our finding that glial morphological diversity exceeds detectable transcriptional diversity. Nonetheless, we have been cautious to draw attention to the limitations of our study as suggested by the reviewers and emphasize the value of this work as a community resource.

## References

Abdo H, Calvo-enrique L, Lopez JM, Song J, Zhang M, Usoskin D, Manira A El, Adameyko I, Hjerling-leffler J, Ernfors P. 2019. Specialized cutaneous Schwann cells initiate pain sensation. Science (80*-)* 365:695–699.

Allen AM, Neville MC, Birtles S, Croset V, Treiber CD, Waddell S, Goodwin SF. 2020. A single-cell transcriptomic atlas of the adult \textit{Drosophila} ventral nerve cord. Elife 9:e54074. doi:10.7554/eLife.54074

Allen NJ, Lyons DA. 2018. Glia as architects of central nervous system formation and function. Science (80*-)* 362:181–185.

Apitz H, Salecker I. 2014. A challenge of numbers and diversity: neurogenesis in the Drosophila optic lobe. J Neurogenet 28:1–35. doi:10.3109/01677063.2014.922558

Banerjee S, Mino RE, Fisher ES, Bhat MA. 2017. A versatile genetic tool to study midline glia function in the Drosophila CNS. Dev Biol 429:35–43. doi:https://doi.org/10.1016/j.ydbio.2017.06.010

Batiuk MY, Martirosyan A, Wahis J, Vin F De, Marneffe C, Kusserow C, Koeppen J, Viana JF, Oliveira JF, Voet T, Ponting CP, Belgard TG, Holt MG. 2020. Identification of region-specific astrocyte subtypes at single cell resolution 1–15. doi:10.1038/s41467-019-14198-8

Bayraktar OA, Bartels T, Holmqvist S, Kleshchevnikov V, Martirosyan A, Polioudakis D, Haim L Ben, Young AMH, Batiuk MY, Prakash K, Brown A, Roberts K, Paredes MF, Kawaguchi R, Stockley JH, Sabeur K, Chang SM, Huang E, Hutchinson P, Ullian EM, Hemberg M, Coppola G, Holt MG, Geschwind DH, Rowitch DH. 2020. Astrocyte layers in the mammalian cerebral cortex revealed by a single-cell in situ transcriptomic map. Nat Neurosci 23. doi:10.1038/s41593-020-0602-1

Beckervordersandforth RM, Rickert C, Altenhein B, Technau GM. 2008. Subtypes of glial cells in the Drosophila embryonic ventral nerve cord as related to lineage and gene expression. Mech Dev 125:542–57. doi:10.1016/j.mod.2007.12.004

Bittern J, Pogodalla N, Ohm H, Schirmeier S, Klämbt C, Brüser L, Kottmeier R. 2021. Neuron – glia interaction in the Drosophila nervous system. Dev Neurobiol 81:438–452. doi:10.1002/dneu.22737

Chai H, Diaz-castro B, Shigetomi E, Whitelegge JP, Coppola G, Khakh BS. 2017. Neural Circuit- Specialized Astrocytes : Transcriptomic, Proteomic, Morphological, and Functional Evidence. Neuron 95:531–549. doi:10.1016/j.neuron.2017.06.029

Choi HMT, Schwarzkopf M, Fornace ME, Acharya A, Artavanis G, Stegmaier J, Cunha A, Pierce NA. 2018. Third-generation in situ hybridization chain reaction : multiplexed, quantitative, sensitive, versatile, robust. Development 145:dev165753. doi:10.1242/dev.165753

Chotard C, Salecker I. 2007. Glial cell development and function in the Drosophila visual system. Neuron Glia Biol 3:17–25. doi:10.1017/S1740925X07000592

Ciappelloni S, Murphy-Royal C, Dupuis JP, Oliet SHR, Groc L. 2017. Dynamics of surface neurotransmitter receptors and transporters in glial cells: Single molecule insights. Cell Calcium 67:46–52. doi:https://doi.org/10.1016/j.ceca.2017.08.009

Corrales M, Cocanougher BT, Kohn AB, Wittenbach JD, Long XS, Lemire A, Cardona A, Singer RH, Moroz LL, Zlatic M. 2022. A single-cell transcriptomic atlas of complete insect nervous systems across multiple life stages. Neural Dev 17:8. doi:10.1186/s13064-022-00164-6

Coutinho-Budd J, Freeman MR. 2013. Probing the enigma: Unraveling glial cell biology in invertebrates. Curr Opin Neurobiol 23:1073–1079. doi:10.1016/j.conb.2013.07.002

Coutinho-Budd JC, Sheehan AE, Freeman MR. 2017. The secreted neurotrophin Spätzle 3 promotes glial morphogenesis and supports neuronal survival and function. Genes Dev 31:2023–2038. doi:10.1101/gad.305888.117.GENES

Crews ST, Thomas JB, Goodman CS. 1988. The Drosophila single-minded gene encodes a nuclear protein with sequence similarity to the per gene product. Cell 52:143–151.

Crisp S, Evers JF, Fiala A, Bate M. 2008. The development of motor coordination in Drosophilaembryos. Development 135:3707–3717. doi:10.1242/dev.026773

Davis FP, Nern A, Picard S, Reiser MB, Rubin GM, Eddy SR, Henry GL. 2020. A genetic, genomic, and computational resource for exploring neural circuit function. Elife 9:e50901.

Dimou L, Gallo V. 2015. NG2-Glia and Their Functions in the Central Nervous System. Glia 63:1429– 1451. doi:10.1002/glia.22859

Doherty J, Logan MA, Taşdemir ÖE, Freeman MR. 2009. Ensheathing Glia Function as Phagocytes in the Adult Drosophila Brain. J Neurosci 29:4768–4781. doi:10.1523/JNEUROSCI.5951-08.2009

Dohn TE, Cripps RM. 2018. Absence of the Drosophila Jump Muscle Actin Act79B is Compensated by Up-regulation of Act88F. Dev Dyn 247:642–649. doi:10.1002/dvdy.24616

Durkee CA, Araque A. 2019. Diversity and Specificity of Astrocyte–neuron Communication. Neuroscience 396:73–78. doi:https://doi.org/10.1016/j.neuroscience.2018.11.010

Edwards TN, Meinertzhagen IA. 2010. The functional organisation of glia in the adult brain of Drosophila and other insects. Prog Neurobiol 90:471–497. doi:10.1016/j.pneurobio.2010.01.001.The

Edwards TN, Nuschke AC, Nern A, Meinertzhagen I a. 2012. Organization and metamorphosis of glia in the Drosophila visual system. J Comp Neurol 520:2067–2085. doi:10.1002/cne.23071

Erclik T, Li X, Courgeon M, Bertet C, Chen Z, Baumert R, Ng J, Koo C, Arain U, Behnia R, Del A, Rodriguez V, Senderowicz L, Negre N, Kevin P. 2017. Integration of temporal and spatial patterning generates neural diversity. Nat Publ Gr 541:365–370. doi:10.1038/nature20794

Escartin C, Galea E, Laktos A, O’Callaghan JP, Carmignoto G, Agarwal A, Allen NJ, Araque A, Barbeito L, Quintana FJ, Ransohoff RM, Riquelme-perez M, Robel S. 2021. Reactive astrocyte nomenclature, definitions, and future directions. Nat Neurosci 24:312–325. doi:10.1038/s41593-020-00783-4

Evans IR, Hu N, Skaer H, Wood W. 2010. Interdependence of macrophage migration and ventral nerve cord development in Drosophila embryos. Development 137:1625–1633. doi:10.1242/dev.046797

Fernandes VM, Chen Z, Rossi AM, Zipfel J, Desplan C. 2017. Glia relay differentiation cues to coordinate neuronal development in Drosophila. Science (80-) 357:886–891.

Fischbach KF, Dittrich APM. 1989. The optic lobe of Drosophila melanogaster. I: A. Golgi analysis of wild-type structure. Cell Tissue Res 258:441–475. doi:doi: 10.1007/BF00218858

Foerster S, Hill MFE, Franklin RJM. 2019. Diversity in the oligodendrocyte lineage : Plasticity or heterogeneity ? Glia 67:1797–1805. doi:10.1002/glia.23607

Freeman MR. 2015. Drosophila Central Nervous System Glia. Cold Spring Harb Perspect Biol 7:a020552.

Freeman MR, Doherty J. 2006. Glial cell biology in Drosophila and vertebrates. Trends Neurosci 29:82–90. doi:10.1016/j.tins.2005.12.002

Goto A, Kadowaki T, Kitagawa Y. 2003. Drosophila hemolectin gene is expressed in embryonic and larval hemocytes and its knock down causes bleeding defects. Dev Biol 264:582–591. doi:10.1016/j.ydbio.2003.06.001

Grabert K, Michoel T, Karavolos MH, Clohisey S, Baillie JK, Stevens MP, Freeman TC, Summers KM, Mccoll BW. 2016. Microglial brain region − dependent diversity and selective regional sensitivities to aging. Nat Neurosci 19:504–516. doi:10.1038/nn.4222

Grueber WB, Ye B, Yang C-H, Younger S, Borden K, Jan LY, Jan Y-N. 2007. Projections of Drosophila multidendritic neurons in the central nervous system: links with peripheral dendrite morphology.

Hao Y, Hao S, Andersen-nissen E, Gottardo R, Smibert P, Hao Y, Hao S, Andersen-nissen E, Iii WMM, Zheng S, Butler A, Papalexi E, Mimitou EP, Jain J, Srivastava A, Stuart T, Fleming LM, Yeung B. 2021. Resource Integrated analysis of multimodal single-cell data ll ll Integrated analysis of multimodal single-cell data. Cell 184:3573–3587. doi:10.1016/j.cell.2021.04.048

Hartenstein V. 2011. Morphological diversity and development of glia in Drosophila. Glia 59:1237– 1252.

Hartenstein V, Nassif C, Lekven A. 1998. Embryonic Development of the Drosophila Brain. II. Pattern of Glial Cells. J Comp Neurol 402:32–47.

Harty B, Monk K. 2017. Unwrapping the unappreciated: recent progress in Remak Schwann cell biology. Curr Opin Neurobiol 47:131–137.

Hidalgo A. 2003. Neuron–glia interactions during axon guidance in Drosophila. Biochem Soc Trans 31:50–55.

Ho T, Wu W, Hung S, Liu T, Lee Y. 2019. Expressional Profiling of Carpet Glia in the Developing Drosophila Eye Reveals Its Molecular Signature of Morphology Regulators. Front Neurosci 13:244. doi:10.3389/fnins.2019.00244

Ito K, Urban J, Technau GM. 1995. Distribution, classification, and development of Drosophila glial cells in the late embryonic and early larval ventral nerve cord. Roux’s Arch Dev Biol 204:284–307. doi:10.1007/BF02179499

Jacobs JR. 2000. The Midline Glia of Drosophila: a molecular genetic model for the developmental functions of Glia. Prog Neurobiol 62:475–508. doi:https://doi.org/10.1016/S0301-0082(00)00016-2

Jin LH, Shim J, Yoon JS, Kim B, Kim J, Kim-Ha J, Kim Y-J. 2008. Identification and Functional Analysis of Antifungal Immune Response Genes in Drosophila. PLOS Pathog 4:e1000168.

Kamen Y, Pivonkova H, Evans KA, Káradóttir RT. 2022. A Matter of State : Diversity in Oligodendrocyte Lineage Cells. doi:10.1177/1073858420987208

Kato K, Orihara-ono M, Awasaki T. 2020. Multiple lineages enable robust development of the neuropil-glia architecture in adult Drosophila. doi:10.1242/dev.184085

Kearney JB, Wheeler SR, Estes P, Parente B, Crews ST. 2004. Gene expression profiling of the developing Drosophila CNS midline cells. Dev Biol 275:473–492.

Khakh BS, Deneen B. 2019. The Emerging Nature of Astrocyte Diversity.

Khakh BS, Sofroniew M V. 2015. Diversity of astrocyte functions and phenotypes in neural circuits. doi:10.1038/nn.4043

Kongton K, McCall K, Phongdara A. 2014. Identification of gamma-interferon-inducible lysosomal thiol reductase (GILT) homologues in the fruit fly Drosophila melanogaster. Dev Comp Immunol 44:389–396. doi:https://doi.org/10.1016/j.dci.2014.01.007

Konstantinides N, Rossi AM, Escobar A, Dudragne L, Chen Y-C, Tran T, Martinez A, Özel MN, Simon F, Shao Z, Nadejda M, Fullard JF, Walldorf U, Roussos P, Konstantinides N, Jaimes AM, Özel MN, Simon F, Shao Z, Tsankova NM, Fullard JF, Walldorf U, Roussos P, Desplan C. 2021. A comprehensive series of temporal transcription factors in the fly visual system. bioRxiv 06:2021.06.13.448242.

Kosman D, Ip YT, Levine M, Arora K. 1991. Establishment of the Mesoderm-Neuroectoderm Boundary in the Drosophila Embryo. Science (80-) 254:118–122. doi:10.1126/science.1925551

Kremer MC, Jung C, Batelli S, Rubin GM, Gaul U. 2017. The Glia of the Adult Drosophila Nervous System. Glia 65:606–638. doi:10.1002/glia.23115

Kurmangaliyev YZ, Yoo J, Valdes-aleman J, Sanfilippo P, Zipursky SL, Kurmangaliyev YZ, Yoo J, Valdes-aleman J, Sanfilippo P, Zipursky SL. 2020. Transcriptional Programs of Circuit Assembly in the Drosophila Visual System. Neuron 108:1045–1057. doi:10.1016/j.neuron.2020.10.006

Lago-Baldaia I, Fernandes VM, Ackerman SD, Czopka T, Smith CJ, Schirmeier S, Ackerman SD. 2020. More Than Mortar : Glia as Architects of Nervous System Development and Disease 8:611269. doi:10.3389/fcell.2020.611269

Lancichinetti A, Fortunato S. 2011. Limits of modularity maximization in community detection AND SPLITTING CLUSTERS. Phys Rev E 84:066122. doi:10.1103/PhysRevE.84.066122

Landgraf M, Sánchez-Soriano N, Technau GM, Urban J, Prokop A. 2003. Charting the Drosophila neuropile: a strategy for the standardised characterisation of genetically amenable neurites. Dev Biol 260:207–225.

Lanjakornsiripan D, Pior B-J, Kawaguchi D, Furutachi S, Tahara T, Katsuyama Y, Suzuki Y, Fukazawa Y, Gotoh Y. 2018. Layer-specific morphological and molecular differences in neocortical astrocytes and their dependence on neuronal layers. Nat Commun 9:1623. doi:10.1038/s41467-018-03940-3

Lassetter AP, Corty MM, Barria R, Sheehan AE, Hill J, Aicher SA, Fox AN, Freeman MR. 2021. Glial TGFβ activity promotes axon survival in peripheral nerves. bioRxiv 2021.09.02.458753. doi:10.1101/2021.09.02.458753

Latsenko I, Marra A, Boquete J-P, Peña J, Lemaitre B. 2020. Iron sequestration by transferrin 1 mediates nutritional immunity in Drosophila melanogaster. Proc Natl Acad Sci 117:7317–7325. doi:10.1073/pnas.1914830117

Li X, Erclik T, Bertet C, Chen Z, Voutev R, Venkatesh S, Morante J, Celik A, Desplan C. 2013. Temporal patterning of Drosophila medulla neuroblasts controls neural fates. Nature 498:456–462. doi:10.1038/nature12319

Mark B, Lai S, Zarin AA, Manning L, Pollington HQ, Litwin-kumar A, Cardona A, Truman JW, Doe CQ. 2021. A developmental framework linking neurogenesis and circuit formation in the Drosophila CNS. Elife 10:e67510.

Marques S, Van Bruggen D, Vanichkina DP, Hjerling-leffler J, Taft RJ, Marques S, Bruggen D Van, Vanichkina DP, Floriddia EM. 2018. Transcriptional Convergence of Oligodendrocyte Lineage Progenitors during Development Resource Transcriptional Convergence of Oligodendrocyte Lineage Progenitors during Development. Dev Cell 46:504–517. doi:10.1016/j.devcel.2018.07.005

Marsh SE, Walker AJ, Kamath T, Dissing-olesen L, Hammond TR, Soysa TY De, Young AMH, Murphy S, Abdulraouf A, Nadaf N, Dufort C, Walker AC, Lucca LE, Kozareva V, Vanderburg C, Hong S, Bulstrode H, Hutchinson PJ, Gaffney DJ, Hafler DA, Franklin RJM, Macosko EZ. 2022. Dissection of artifactual and confounding glial signatures by single-cell sequencing of mouse and human brain. Nat Neurosci 25:306–316. doi:10.1038/s41593-022-01022-8

Mayer F, Mayer N, Chinn L, Pinsonneault RL, Kroetz D, Bainton RJ. 2009. Evolutionary Conservation of Vertebrate Blood – Brain Barrier Chemoprotective Mechanisms in Drosophila. J Neurosci 29:3538–3550. doi:10.1523/JNEUROSCI.5564-08.2009

Merritt DJ, Whitington PM. 1995. Central projections of sensory neurons in the Drosophila embryo correlate with sensory modality, soma position, and proneural gene function. J Neurosci 15:1755–1767.

Murphy-Royal C, Dupuis JP, Varela JA, Panatier A, Pinson B, Baufreton J, Groc L, Oliet SHR. 2015. Surface diffusion of astrocytic glutamate transporters shapes synaptic transmission. Nat Neurosci 18:219–226. doi:10.1038/nn.3901

Muthukumar AK, Stork T, Freeman MR. 2014. Activity-dependent regulation of astrocyte GAT levels during synaptogenesis. Nat Neurosci 17:1340–1350. doi:10.1038/nn.3791

Noordermeer JN, Kopczynski CC, Fetter RD, Bland KS, Chen W-Y, Goodman CS. 1998. Wrapper, a novel member of the Ig superfamily, is expressed by midline glia and is required for them to ensheath commissural axons in Drosophila. Neuron 21:991–1001.

Ohshima S, Villarimo C, Gailey DA. 1997. Reassessment of 79B actin gene expression in the abdomen of adult Drosophila melanogaster. Insect Mol Biol 6:227–231.

Özel MN, Simon F, Jafari S, Holguera I, Chen Y, Benhra N, El-danaf RN, Kapuralin K, Malin JA, Konstantinides N, Desplan C. 2021. Neuronal diversity and convergence in a visual system developmental atlas. Nature 589:88–95. doi:10.1038/s41586-020-2879-3

Parfejevs V, Debbache J, Shakhova O, Schaefer SM, Glausch M, Wegner M, Suter U, Riekstina U, Werner S, Sommer L. 2018. Injury-activated glial cells promote wound healing of the adult skin in mice. Nat Commun 9:236. doi:10.1038/s41467-017-01488-2

Peco E, Davla S, Camp D, Stacey SM, Landgraf M, Meyel DJ Van. 2016. Drosophila astrocytes cover specific territories of the CNS neuropil and are instructed to differentiate by Prospero, a key effector of Notch. Development 143:1170–1181. doi:10.1242/dev.133165

Perz MJ. 1994. The midline glial cell lineage in the post embryonic fruit fly Drosophila melanogaster.

Pfeiffer BD, Jenett A, Hammonds AS, Ngo TB, Misra S, Murphy C, Scully A, Carlson JW, Wan KH, Laverty TR, Mungall C, Svirskas R, Kadonaga JT, Doe CQ, Eisen MB, Celniker SE, Rubin GM. 2008. Tools for neuroanatomy and neurogenetics in Drosophila 105:9715–9720.

Pogodalla N, Winkler B, Klämbt C. 2022. Glial Tiling in the Insect Nervous System. Front Cell Neurosci 16:825695. doi:10.3389/fncel.2022.825695

Ramón y Cajal S. 1899. Histología del sistema nervioso del hombre y de los vertebrados, 1st ed. Madrid: Agencia Estatal Boletín Oficial del Estado.

Prasad AR, Lago-Baldaia I, Bostock MP, Housseini Z, Fernandes VM. 2021. Differentiation signals from glia are fine-tuned to set neuronal numbers during development. bioRxiv 2021.12.13.

Richier B, Vijandi CDM, Mackensen S, Salecker I. 2017. Lapsyn controls branch extension and positioning of astrocyte-like glia in the Drosophila optic lobe. Nat Commun 8:317. doi:10.1038/s41467-017-00384-z

Schwabe T, Bainton RJ, Fetter RD, Heberlein U, Gaul U, Francisco S. 2005. for Blood-Brain Barrier Formation in Drosophila. Cell 123:133–144. doi:10.1016/j.cell.2005.08.037

Simon F, Konstantinides N. 2021. Single-cell transcriptomics in the Drosophila visual system : Advances and perspectives on cell identity regulation, connectivity, and neuronal diversity evolution. Dev Biol 479:107–122. doi:10.1016/j.ydbio.2021.08.001

Sonnenfeld MJ, Jacobs JR. 1995a. Macrophages and glia participate in the removal of apoptotic neurons from the Drosophila embryonic nervous system. J Comp Neurol 359:644–652.

Sonnenfeld MJ, Jacobs JR. 1995b. Apoptosis of the midline glia during Drosophila embryogenesis: a correlation with axon contact. Development 121:569–578.

Spitzer SO, Sitnikov S, Kamen Y, Faria O De, Agathou S, Spitzer SO, Sitnikov S, Kamen Y, Evans KA, Kronenberg-versteeg D. 2019. Oligodendrocyte Progenitor Cells Become Regionally Diverse and Heterogeneous with Age Article Oligodendrocyte Progenitor Cells Become Regionally Diverse and Heterogeneous with Age. Neuron 101:459–471. doi:10.1016/j.neuron.2018.12.020

Stollewerk A, Klämbt C, Cantera R. 1996. Electron microscopic analysis of Drosophila midline glia during embryogenesis and larval development using β-galactosidase expression as endogenous cell marker. Microsc Res Tech 35:294–306.

Stork T, Sheehan A, Tasdemir-yilmaz OE, Freeman MR. 2014. Glia Interactions through the Heartless FGF Receptor Signaling Pathway Mediate Morphogenesis of Drosophila Astrocytes. Neuron 83:388–403. doi:10.1016/j.neuron.2014.06.026

Stork T, Thomas S, Rodrigues F, Silies M, Naffin E, Wenderdel S, Klämbt C. 2009. Drosophila Neurexin IV stabilizes neuron-glia interactions at the CNS midline by binding to Wrapper. Development 136:1251–1261. doi:10.1242/dev.032847

Sun W, Cornwell A, Li J, Peng S, Osorio MJ, Aalling N, Wang S, Benraiss A, Lou N, Goldman SA, Nedergaard M. 2017. SOX9 Is an Astrocyte-Specific Nuclear Marker in the Adult Brain Outside the Neurogenic Regions. J Neurosci 37:4493 LP – 4507. doi:10.1523/JNEUROSCI.3199-16.2017

Thomas JB, Crews ST, Goodman CS. 1988. Molecular Genetics of the single-minded Locus: A Gene Involved in the Development of the Drosophila Nervous System. Cell 52:133–141.

Vaessin H, Grell E, Wolff E, Bier E, Jan LY, Jan YN. 1991. prospero Is Expressed in Neuronal Precursors and Encodes a Nuclear Protein That Is Involved in the Control of Axonal Outgrowth in Drosophila. Cell 67:941–953.

Vasenkova I, Luginbuhl D, Chiba A. 2006. Gliopodia extend the range of direct glia–neuron communication during the CNS development in Drosophila. Mol Cell Neurosci 31:123–130.

Volkenhoff A, Weiler A, Letzel M, Stehling M, Klämbt C, Schirmeier S. 2015. Glial Glycolysis Is Essential for Neuronal Survival in Article Glial Glycolysis Is Essential for Neuronal Survival in Drosophila. Cell Metab 22:437–447. doi:10.1016/j.cmet.2015.07.006

Weber JJ, Brummett LM, Coca ME, Tabunoki H, Kanost MR, Ragan EJ, Park Y, Gorman MJ. 2022. Phenotypic analyses, protein localization, and bacteriostatic activity of Drosophila melanogaster transferrin-1. Insect Biochem Mol Biol 147:103811. doi:https://doi.org/10.1016/j.ibmb.2022.103811

Westergard T, Rothstein JD. 2020. Astrocyte Diversity : Current Insights and Future Directions. Neurochem Res 45:1298–1305. doi:10.1007/s11064-020-02959-7

Wheeler SR, Kearney JB, Guardiola AR, Crews ST. 2006. Single-cell mapping of neural and glial gene expression in the developing Drosophila CNS midline cells. Dev Biol 294:509–524.

Wickham H. 2016. ggplot2: Elegant Graphics for Data Analysis. New York: Springer-Verlag.

Yu G, Wang L-G, Han Y, He Q-Y. 2012. clusterProfiler : an R Package for Comparing Biological Themes Among Gene Clusters. Omi A J Integr Biol 16:284–287. doi:10.1089/omi.2011.0118

Zhang Y, Lowe S, Li X. 2023. Notch-dependent binary fate choice regulates the Netrin pathway to control axon guidance of Drosophila visual projection neurons. Cell Rep 42:112143.

Zheng GXY, Terry JM, Belgrader P, Ryvkin P, Bent ZW, Wilson R, Ziraldo SB, Wheeler TD, Mcdermott GP, Zhu J, Gregory MT, Shuga J, Montesclaros L, Underwood JG, Masquelier DA, Nishimura SY, Schnall-levin M, Wyatt PW, Hindson CM, Bharadwaj R, Wong A, Ness KD, Beppu LW, Deeg HJ, Mcfarland C, Loeb KR, Valente WJ, Ericson NG, Stevens EA, Radich JP, Mikkelsen TS, Hindson BJ, Bielas JH. 2017. Massively parallel digital transcriptional profiling of single cells. Nat Commun 8:14049. doi:10.1038/ncomms14049

Zhou B, Xia Y, Ruo Z, Jiang T. 2019. Astrocyte morphology : Diversity, plasticity, and role in neurological diseases 665–673. doi:10.1111/cns.13123

Zhu H, Zhao SD, Ray A, Zhang Y, Li X. 2022. A comprehensive temporal patterning gene network in Drosophila medulla neuroblasts revealed by single-cell RNA sequencing. Nat Commun 13:1247. doi:10.1038/s41467-022-28915-3

Zlatic M, Li F, Strigini M, Grueber W, Bate M. 2009. Positional Cues in the Drosophila Nerve Cord : Semaphorins Pattern the Dorso-Ventral Axis. PLoS Biol 7:e1000135. doi:10.1371/journal.pbio.1000135

