## Supplementary figures and images for "A *Drosophila* glial cell atlas reveals a mismatch between detectable transcriptional diversity and morphological diversity"

### Supplementary file 3

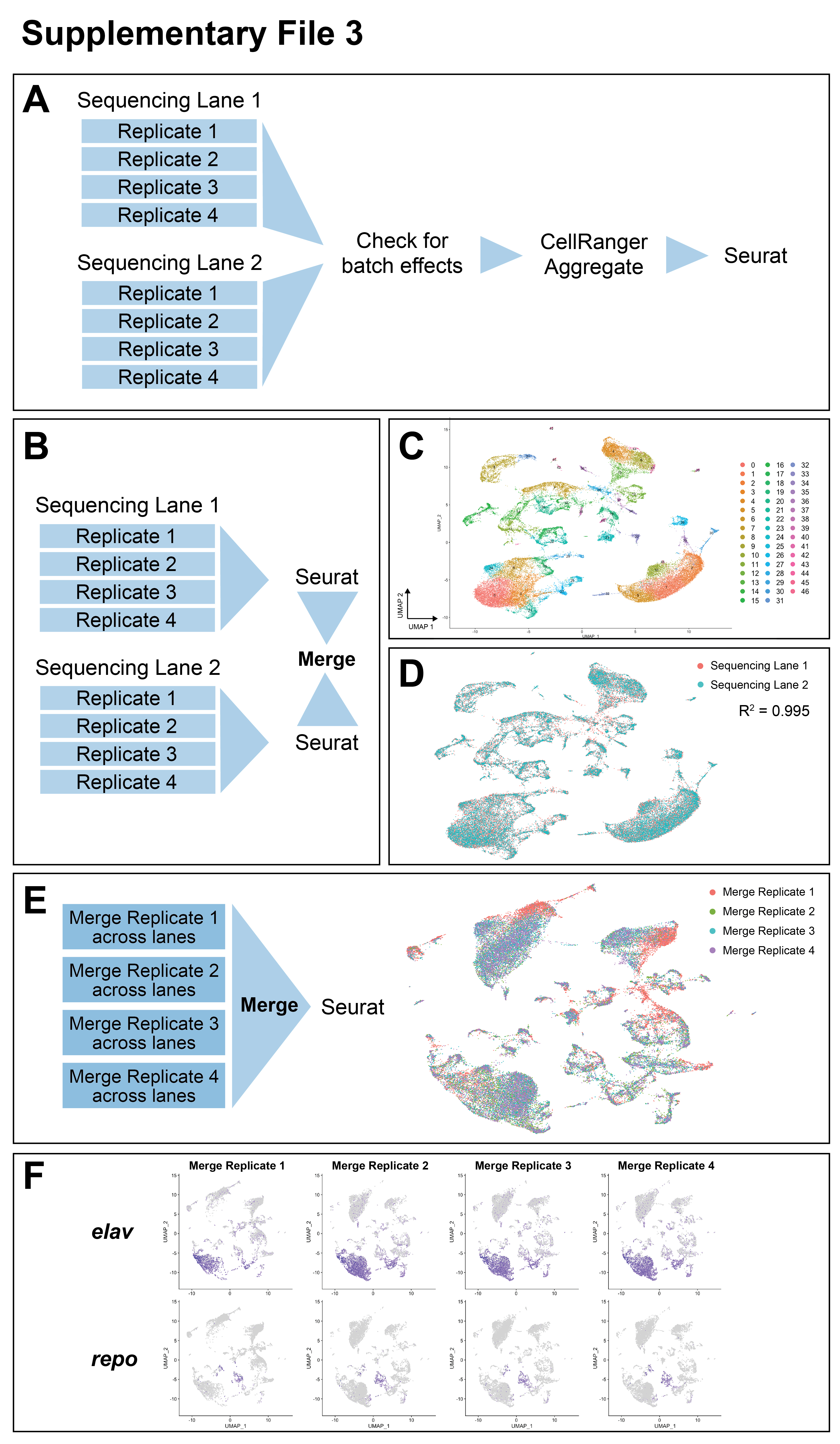
