## Supplementary file 4 for "A *Drosophila* glial cell atlas reveals a mismatch between detectable transcriptional diversity and morphological diversity"

**Supplementary file 4: List of specific genotypes and conditions used by figure panel**

| **Fig.** | **Panel** | **Glial Subtype** | **Genotype** | **Conditions** |
| --- | --- | --- | --- | --- |
| 1 | B, C, D, E, F | Embryonic channel and surface-only perineurial | *w[1118]/pBPhsFlp2;* *TI{GFP[3xP3.cLa]=CRIMIC.TG4.2}CG5080[CR01346-TG4.2]/+; UAS-HA_V5_FLAG/+* | Raised at 25^o^C before and after heat shocking. 6 hour after egg laying embryos were heat stocked at 37^o^C for 15 minutes, quenched at 4^o^C for 10 minutes, and reared at 25^o^C until larval hatching. |
| 1 - S1 | A |  |  |  |
| 1 | G, H, I, J, K, L | Embryonic surface and channel subperineurial | *w[1118]/pBPhsFlp2;* *moody-GAL4.SPG; UAS-HA_V5_FLAG/+* | Raised at 25^o^C before and after heat shocking. 6 hour after egg laying embryos were heat stocked at 37^o^C for 15 minutes, quenched at 4^o^C for 10 minutes, and reared at 25^o^C until larval hatching. |
| 1 | M | Embryonic cortex | *w[1118]/pBPhsFlp2; sp/+; R54H02-Gal4/UAS- HA_V5_FLAG* | Raised at 25^o^C before and after heat shocking. 6 hour after egg laying embryos were heat stocked at 37^o^C for 15 minutes, quenched at 4^o^C for 10 minutes, and reared at 25^o^C until larval hatching. |
| 1 | N, O | Embryonic ensheathing | *w[1118]/pBPhsFlp2;* *TI{CRIMIC.TG4.2}Eaat2[CR00503-TG4.2]/+; UAS-HA_V5_FLAG/+* | Raised at 25^o^C before and after heat shocking. 6 hour after egg laying embryos were heat stocked at 37^o^C for 15 minutes, quenched at 4^o^C for 10 minutes, and reared at 25^o^C until larval hatching. |
| 1 | P, Q, R, S | Embryonic astrocyte | *w[1118]/pBPhsFlp2;; alrm-Gal4/ UAS-HA_V5_FLAG* | Raised at 25^o^C before and after heat shocking. 6 hour after egg laying embryos were heat stocked at 37^o^C for 15 minutes, quenched at 4^o^C for 10 minutes, and reared at 25^o^C until larval hatching. |
| 1 - S1 | B, C, C’ | Embryonic channel and surface-only perineurial | *w[1118]/pBPhsFlp2;* *TI{GFP[3xP3.cLa]=CRIMIC.TG4.2}CG5080[CR01346-TG4.2]/lexAop-myr::tdTomato; UAS-HA_V5_FLAG/RepoLexA* | Raised at 25^o^C before and after heat shocking. 6 hour after egg laying embryos were heat stocked at 37^o^C for 15 minutes, quenched at 4^o^C for 10 minutes, and reared at 25^o^C until larval hatching. |
| 1 - S2 | B, C, D | Embryonic astrocyte | *w[1118]/pBPhsFlp2;; alrm-Gal4/ UAS-HA_V5_FLAG* | Raised at 25^o^C before and after heat shocking. 6 hour after egg laying embryos were heat stocked at 37^o^C for 15 minutes, quenched at 4^o^C for 10 minutes, and reared at 25^o^C until larval hatching. |
| 2 | B | Perineurial | *w[1118]/pBPhsFlp2;; R85G01-Gal4/UAS-HA_V5_FLAG* | Raised at 18^o^C before and after heat shocking. 0-5 day-old adults were heat shocked at 37^o^C for 2 minutes. |
| 2 | C | Subperineurial | *w[1118]/pBPhsFlp2;; R54C07-Gal4/UAS-HA_V5_FLAG* | Raised at 18^o^C before and after heat shocking. 0-5 day-old adults were heat shocked at 37^o^C for 3 minutes. |
| 2 | D | Chalice | *w[1118]/pBPhsFlp2;; R10C12-Gal4/UAS-HA_V5_FLAG* | Raised at 18^o^C before and after heat shocking. 0-5 day-old adults were heat shocked at 37^o^C for 5 minutes. |
| 2 | E | Carpet | *w[1118]/pBPhsFlp2;; R54C07-Gal4/UAS-HA_V5_FLAG* | Raised at 18^o^C before and after heat shocking. 0-5 day-old adults were heat shocked at 37^o^C for 1 minutes. |
| 2 | F | Fenestrated | *w[1118]/pBPhsFlp2;; R47G01-Gal4/UAS-HA_V5_FLAG* | Raised at 18^o^C before and after heat shocking. 0-5 day-old adults were heat shocked at 37^o^C for 3 minutes. |
| 2 | G | Pseudo-cartridge | *w[1118]/pBPhsFlp2;; R54C07-Gal4/UAS-HA_V5_FLAG* | Raised at 18^o^C before and after heat shocking. 0-5 day-old adults were heat shocked at 37^o^C for 2 minutes. |
| 2 | H | Cortex | *w[1118]/pBPhsFlp2; sp/+; R54H02-Gal4/UAS- HA_V5_FLAG_OLLAS* | Raised at 18^o^C before and after heat shocking. 0-5 day-old adults were heat shocked at 37^o^C for 4 minutes. |
| 2 | I | Distal satellite and chiasm | *w[1118]/pBPhsFlp2;; R53H12-Gal4/UAS- HA_V5_FLAG* | Raised at 18^o^C before and after heat shocking. 0-5 day-old adults were heat shocked at 37^o^C for 4 minutes. |
| 2 | J | Proximal satellite | *w[1118]/pBPhsFlp2; sp/+; R54H02/UAS_HA_V5_FLAG_OLLAS*  ♂︎  *pBPhsflp2; sp/+; Repo-Gal4/ UAS- HA_V5_FLAG* | Raised at 18^o^C before and after heat shocking. 0-5 day-old adults were heat shocked at 37^o^C for **6** minutes. |
| 2 | K | Chiasm | *w[1118]/pBPhsFlp2;; R53H12-Gal4/UAS- HA_V5_FLAG* | Raised at 18^o^C before and after heat shocking. 0-5 day-old adults were heat shocked at 37^o^C for 4 minutes. |
| 2 | L | Ensheathing | ♀︎ *yw, hsflp[122]/ pBPhsflp2; sp/+; R56F03-Gal4/UAS- HA_V5_FLAG* | Raised at 18^o^C before and after heat shocking. 0-5 day-old adults were heat shocked at 37^o^C for 2 minutes. |
| 2 | M, N | Ensheathing | ♂︎  *pBPhsflp2; sp/+; R56F03-Gal4/UAS- HA_V5_FLAG* | Raised at 18^o^C before and after heat shocking. 0-5 day-old adults were heat shocked at 37^o^C for 5 minutes. |
| 2 | O | Ensheathing | ♂︎  *pBPhsflp2; sp/+; R56F03-Gal4/UAS- HA_V5_FLAG* | Raised at 18^o^C before and after heat shocking. 0-5 day-old adults were heat shocked at 37^o^C for 4 minutes. |
| 2 | P | Marginal | *w[1118]/pBPhsFlp2; R35E04-Gal4/+; UAS- HA_V5_FLAG_OLLAS/+* | Raised at 18^o^C before and after heat shocking. 0-5 day-old adults were heat shocked at 37^o^C for 2 minutes. |
| 2 | Q | Astrocyte | ♀︎ *yw, hsflp[122]/pBPhsFlp2;; R86E01-Gal4/UAS-HA_V5_FLAG* | Raised at 18^o^C before and after heat shocking. 0-5 day-old adults were heat shocked at 37^o^C for 2 minutes. |
| 2 | R, S, T, U, V | Astrocyte | ♂︎  *pBPhsFlp2;; R86E01-Gal4/UAS-HA_V5_FLAG* | Raised at 18^o^C before and after heat shocking. 0-5 day-old adults were heat shocked at 37^o^C for 5 minutes. |
| 2 | W | Epithelial | ♂︎  *pBPhsflp2; sp/+; Repo-Gal4/ UAS- HA_V5_FLAG* | Raised at 18^o^C before and after heat shocking. 0-5 day-old adults were heat shocked at 37^o^C for 3 minutes. |
| 2 - S1 | A, B | Perineurial | *w[1118]; UAS-CD8::GFP; R85G01-Gal4* | Raised at 25^o^C. Adults dissected between 0-7 days old. |
| 5 - S4 | E, F, F’, I, I’ |  |  |  |
| 2 - S1 | C, D | Subperineurial | *w[1118]; UAS-CD8::GFP; R54C07-Gal4* | Raised at 25^o^C. Adults dissected between 0-7 days old. |
| 2 - S1 | E | Astrocyte | *w[1118]; UAS-CD8::GFP; R86E01-Gal4* | Raised at 25^o^C. Adults dissected between 0-7 days old. |
| 2 - S1 | F | Ensheathing | *w[1118]; UAS-CD8::GFP; R56F03-Gal4* | Raised at 25^o^C. Adults dissected between 0-7 days old. |
| 2 - S1 | G | Chiasm | *w[1118]; UAS-CD8::GFP; R53H12-Gal4* | Raised at 25^o^C. Adults dissected between 0-7 days old. |
| 5 - S4 | M, M’, M’’ |  |  |  |
| 2 - S1 | H | Cortex | *w[1118]; UAS-CD8::GFP; R54H02-Gal4* | Raised at 25^o^C. Adults dissected between 0-7 days old. |
| 2 - S1 | I | Epithelial | *w[1118]; UAS-CD8::GFP; R55B03-Gal4* | Raised at 25^o^C. Adults dissected between 0-7 days old. |
| 2 - S1 | J | Marginal | *w[1118]; UAS-CD8::GFP; R35E04-Gal4* | Raised at 25^o^C. Adults dissected between 0-7 days old. |
| 2 - S1 | K | Proximal satellite | *w[1118]; UAS-CD8::GFP; R46H12-Gal4* | Raised at 25^o^C. Adults dissected between 0-7 days old. |
| 5 - S4 | N, N’, N’’ |  |  |  |
| 2 - S1 | L | Pseudo-cartridge | *w[1118]; UAS-CD8::GFP; R50A12-Gal4* | Raised at 25^o^C. Adults dissected between 0-7 days old. |
| 2 - S1 | M | Fenestrated | *w[1118]; UAS-CD8::GFP; R47G01-Gal4* | Raised at 25^o^C. Adults dissected between 0-7 days old. |
| 5 - S4 | H, H’ |  |  |  |
| 2 - S2 | B, C, D, E | Astrocyte | *yw, hsflp[122]/pBPhsFlp2;; R86E01- Gal4/UAS-HA_V5_FLAG* | Raised at 18oC before and after heat shocking. 0-5 day- old adults were heat shocked at 37oC for 3 minutes. |
| 5 | C, D, D’ | Epithelial | *y[1]w[*];; CG43795-GFP* | Raised at 25^o^C. Adults dissected between 0-7 days old. |
| 5 | E, F, F’ | Marginal | *w[1118]; UAS-nls::GFP; R35E04-Gal4* | Raised at 25^o^C. Adults dissected between 0-7 days old. |
| 5 | G, H, H’ | Proximal satellite | *w[1118]; UAS-nls::GFP; R46H12-Gal4* | Raised at 25^o^C. Adults dissected between 0-7 days old. |
| 5 - S4 | K, K’ |  |  |  |
| 6 | B, B’, B’’,D, D’, D’’, F, F’, F’’ | Astrocyte | *w[1118]; UAS-nls::GFP; R86E01-Gal4* | Raised at 25^o^C. Adults dissected between 0-7 days old. |
| 7 | C, C’, C’’ |  |  |  |
| 5 - S2 | A, A’, A’’ |  |  |  |
| 3 - S1 | G, G’, H, H’ | Embryonic midline glia | *w[1118]/pBPhsFlp2; wrapper-gal4/+; UAS-HA_V5_FLAG/+* | Raised at 25^o^C before and after heat shocking. 6 hour after egg laying embryos were heat stocked at 37^o^C for 15 minutes, quenched at 4^o^C for 10 minutes, and reared at 25^o^C until embryonic stage 17. |
| 3 - S4 | A, A’, A’’, B, B’, B’’, B’’’, C, C’, C’’, C’’’ | All glia | *w[1118]; UAS-nls::GFP; Repo-Gal4* | Raised at 25^o^C. Adults dissected between 0-7 days old. |
| 3 - S4 | D, D’ | Neurons | *w[1118]/pBPhsFlp2;; Mi{Trojan-GAL4.1}Rdl[MI02957-TG4.1]/ UAS-HA_V5_FLAG* | Raised at 25^o^C before and after heat shocking. 6 hour after egg laying embryos were heat stocked at 37^o^C for 15 minutes, quenched at 4^o^C for 10 minutes, and reared at 25^o^C until larval hatching. |
| 4 | C, C’ | Embryonic surface-only perineurial glia | *w[1118]/pBPhsFlp2;* *TI{CRIMIC.TG4.2}pippin[CR01460-TG4.2]/+; UAS-HA_V5_FLAG/+* | Raised at 25^o^C before and after heat shocking. 6 hour after egg laying embryos were heat stocked at 37^o^C for 15 minutes, quenched at 4^o^C for 10 minutes, and reared at 25^o^C until larval hatching. |
| 4 | D, D’ | Embryonic channel and surface-only perineurial glia | *w[1118]/pBPhsFlp2;; Mi{Trojan-GAL4.1}CG6126[MI12300-TG4.1]/ UAS-HA_V5_FLAG* | Raised at 25^o^C before and after heat shocking. 6 hour after egg laying embryos were heat stocked at 37^o^C for 15 minutes, quenched at 4^o^C for 10 minutes, and reared at 25^o^C until larval hatching. |
| 4 - S1 | B, B’ |  |  |  |
| 4 | E, E’ | Embryonic channel and surface-only subperineurial glia | *w[1118]/pBPhsFlp2;Mi{Trojan-GAL4.0}CG10702[MI15239-TG4.0] CG17343[MI15239-TG4.0-X]/+; UAS-HA_V5_FLAG/+* | Raised at 25^o^C before and after heat shocking. 6 hour after egg laying embryos were heat stocked at 37^o^C for 15 minutes, quenched at 4^o^C for 10 minutes, and reared at 25^o^C until larval hatching. |
| 4 | F, F’ | Embryonic cortex glia | *w[1118]/pBPhsFlp2; +/+; R54H02(Wrapper)-Gal4/UAS- HA_V5_FLAG* | Raised at 25^o^C before and after heat shocking. 6 hour after egg laying embryos were heat stocked at 37^o^C for 15 minutes, quenched at 4^o^C for 10 minutes, and reared at 25^o^C until larval hatching. |
| 4 | G, G’ | Embryonic ensheathing glia | *w[1118]/pBPhsFlp2;* *TI{CRIMIC.TG4.2}Eaat2[CR00503-TG4.2]/+; UAS-HA_V5_FLAG/+* | Raised at 25^o^C before and after heat shocking. 6 hour after egg laying embryos were heat stocked at 37^o^C for 15 minutes, quenched at 4^o^C for 10 minutes, and reared at 25^o^C until larval hatching. |
| 4 | H, H’ | Embryonic astrocytes | *w[1118]/pBPhsFlp2;; alrm-Gal4/ UAS-HA_V5_FLAG* | Raised at 25^o^C before and after heat shocking. 6 hour after egg laying embryos were heat stocked at 37^o^C for 15 minutes, quenched at 4^o^C for 10 minutes, and reared at 25^o^C until larval hatching. |
| 4 - S1 | C, C’ D | Embryonic channel and surface-only subperineurial glia | *w[1118]/pBPhsFlp2;* *TI{CRIMIC.TG4.2}PRL-1[CR00881-TG4.2]/+; UAS-HA_V5_FLAG/+* | Raised at 25^o^C before and after heat shocking. 6 hour after egg laying embryos were heat stocked at 37^o^C for 15 minutes, quenched at 4^o^C for 10 minutes, and reared at 25^o^C until larval hatching. |
| 4 - S1 | E, E’, F | Embryonic channel and surface-only subperineurial glia | *w[1118]/pBPhsFlp2;; Mi{Trojan-GAL4.2}ltl[MI02191-TG4.2] / UAS- HA_V5_FLAG* | Raised at 25^o^C before and after heat shocking. 6 hour after egg laying embryos were heat stocked at 37^o^C for 15 minutes, quenched at 4^o^C for 10 minutes, and reared at 25^o^C until larval hatching. |
| 4 - S1 | G, G’, H | Embryonic channel and surface-only subperineurial glia | *w[1118]/pBPhsFlp2;P{w[+mW.hs]=GawB}Ntan1[Mz97] P{w[+mC]=UAS-Stinger}2/+; UAS- HA_V5_FLAG/+* | Raised at 25^o^C before and after heat shocking. 6 hour after egg laying embryos were heat stocked at 37^o^C for 15 minutes, quenched at 4^o^C for 10 minutes, and reared at 25^o^C until larval hatching. |
| 4 - S2 | A, A’ | Embryonic astrocyte glia | *w[1118]/pBPhsFlp2;; P{y[+t7.7] w[+mC]=GMR25H07-GAL4} / UAS- HA_V5_FLAG* | Raised at 25^o^C before and after heat shocking. 6 hour after egg laying embryos were heat stocked at 37^o^C for 15 minutes, quenched at 4^o^C for 10 minutes, and reared at 25^o^C until larval hatching. |
| 4 - S2 | B, B’ | Embryonic astrocyte and ensheathing glia | *w[1118]/pBPhsFlp2; sp/+; PBac{GAL4D,EYFP}Tet[PL00243]/ UAS- HA_V5_FLAG* | Raised at 25^o^C before and after heat shocking. 6 hour after egg laying embryos were heat stocked at 37^o^C for 15 minutes, quenched at 4^o^C for 10 minutes, and reared at 25^o^C until larval hatching. |
| 4 - S2 | C, C’ | Embryonic astrocyte and ensheathing glia | *w[1118]/pBPhsFlp2; sp/+; PBac{IT.GAL4}pum[0508-G4]/ UAS- HA_V5_FLAG* | Raised at 25^o^C before and after heat shocking. 6 hour after egg laying embryos were heat stocked at 37^o^C for 15 minutes, quenched at 4^o^C for 10 minutes, and reared at 25^o^C until larval hatching. |
| 4 - S2 | F, F’ | Embryonic astrocyte glia | *w[1118] TI{RFP[3xP3.cUa]=TI}Tre1[attP];;* | Raised at 25^o^C, dissected at 0 hours after larval hatching. |
| 4 - S3 | A, A’ | Embryonic ensheathing and cortex glia | *w[1118]/pBPhsFlp2;* *TI{CRIMIC.TG4.2}ana[CR01446-TG4.2]/+; UAS-HA_V5_FLAG/+* | Raised at 25^o^C before and after heat shocking. 6 hour after egg laying embryos were heat stocked at 37^o^C for 15 minutes, quenched at 4^o^C for 10 minutes, and reared at 25^o^C until larval hatching. |
| 4 - S3 | B, B’ | Embryonic ensheathing and astrocyte glia | *pBPhsFlp2/TI{CRIMIC.TG4.2}CG9657[CR00636-TG4.2];; UAS-HA_V5_FLAG/+* | Raised at 25^o^C before and after heat shocking. 6 hour after egg laying embryos were heat stocked at 37^o^C for 15 minutes, quenched at 4^o^C for 10 minutes, and reared at 25^o^C until larval hatching. |
| 4 - S3 | C, C’ | Embryonic cortex glia | *w[1118]/pBPhsFlp2;; TI{CRIMIC.TG4.0}CG9449[CR01691-TG4.0]/ UAS- HA_V5_FLAG* | Raised at 25^o^C before and after heat shocking. 6 hour after egg laying embryos were heat stocked at 37^o^C for 15 minutes, quenched at 4^o^C for 10 minutes, and reared at 25^o^C until larval hatching. |
| 5 - S2 | B, B’, B’’, C, C’, C’’, D, D’, D’’ | Cortex | *w[1118]; UAS-nls::GFP; R54H02-Gal4* | Raised at 25^o^C. Adults dissected between 0-7 days old. |
| 5 - S2 | E, E’, E’’, F, F’, F’’ | Chiasm | *w[1118]; UAS-nls::GFP; R53H12-Gal4* | Raised at 25^o^C. Adults dissected between 0-7 days old. |
| 5 - S4 | A, B, B’, G, G’ |  |  |  |
| 5 - S3 | A, A’, A’’, B, B’, B’’, F, F’, F’’ | Perineurial | *w[1118]; UAS-nls::GFP; R85G01-Gal4* | Raised at 25^o^C. Adults dissected between 0-7 days old. |
| 5 - S4 | C, D, D’ |  |  |  |
| 5 - S3 | G, G’, G’’, H, H’, H’’, I, I’, I’’, J, J’, J’’ | Ensheathing | *w[1118]; UAS-nls::GFP; R56F03-Gal4* | Raised at 25^o^C. Adults dissected between 0-7 days old. |
| 5 - S3 | C, C’ | *ltl* | *y[1]w[*]/w[*]; ; ltl-Gal4/UAS-myr::GFP* | Raised at 25^o^C. Adults dissected between 0-7 days old. |
| 5 - S3 | D, E | *ltl* | *w[*]/pBPhsFlp2;; ltl-Gal4/UAS-HA_V5_FLAG* | Raised at 18^o^C before and after heat shocking. 0-5 day-old adults were heat shocked at 37^o^C for 1 minute. |
| 5 - S4 | L, L’ |  |  |  |
| 5 - S4 | J, J’ | Fenestrated | *w[1118]; UAS-nls::GFP; R47G01-Gal4* | Raised at 25^o^C. Adults dissected between 0-7 days old. |
